# Adaptive Disruption of Carotenoid Metabolism in Cavefish via Loss of Beta-carotene oxygenase 2a (Bco2a)

**DOI:** 10.1101/2025.10.06.680734

**Authors:** David Peréz Guerra, Sepalika Bandara, Francisca Leal, Johannes von Lintig, Misty R. Riddle

**Affiliations:** University of Nevada, Reno, Reno Nevada; Case Western Reserve University, Cleveland, Ohio; Harvard Medical School, Boston, MA

## Abstract

Vitamin A is an essential micronutrient that supports vision, immunity, and development, yet excess vitamin A can be toxic^1,2^. Most vertebrates maintain tightly regulated vitamin A levels by cleaving dietary carotenoids into retinoids via specialized enzymes, including the mitochondrial carotenoid oxygenase Bco2^3,4^. To what extent genetic changes in carotenoid-metabolism enzymes like Bco2 drive natural variation in vitamin A homeostasis remains unclear. Here we show that multiple cave-adapted populations of the Mexican tetra (*Astyanax mexicanus*) have evolved loss-of-function mutations in *bco2a*, leading to impaired carotenoid cleavage and dramatic carotenoid and vitamin A accumulation. These mutations yield enzymatically inactive Bco2a proteins, are under selection in cavefish, and may provide a physiological advantage under inconsistent carotenoid supply. Contrary to expectations from mammalian models where Bco2 loss induces oxidative damage^5,6^, cavefish appear to tolerate and even benefit from carotenoid accumulation, potentially through enhanced antioxidant capacity and modified mitochondrial metabolism. Our findings reveal that the vitamin A regulatory system is evolutionarily flexible, and that the adaptive loss of Bco2a function may enable survival in nutrient-variable, low-oxygen cave environments. This work provides a framework for understanding how animals balance essential nutrient thresholds in extreme environments.

## Main

The Mexican tetra, *Astyanax mexicanus*, is a species of small freshwater fish that includes both river-dwelling surface fish and multiple independently evolved eyeless cavefish populations that have adapted to life in dark, nutrient-limited underground caves^7^. Cavefish form two genetically and geographically distinct lineages that were established from two different surface fish populations around the same time^8,9^: lineage 1 in the Northern Sierra de Guatemala (e.g. Molino), and lineage 2 in the Southern Sierra de El Abra (e.g., Pachón and Tinaja, Fig 1A). This evolutionary history creates natural replicates for examining the parallel evolution of traits across independently derived cavefish lineages.

**Figure 1.**
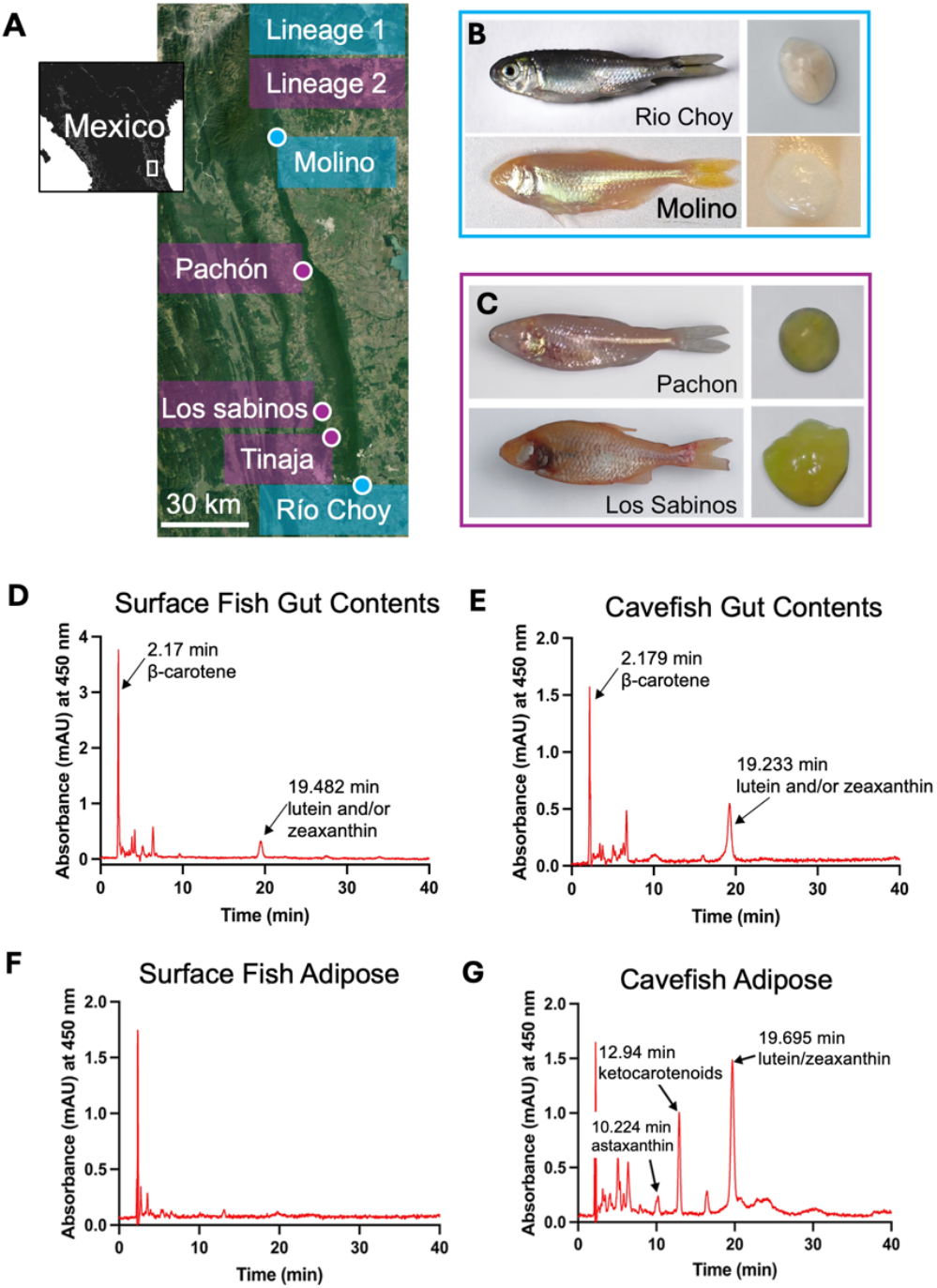
*Astyanax mexicanus* cavefish accumulate carotenoids in the visceral adipose tissue. **A**, Map of Northeastern Mexico showing collection sites for field samples, including surface fish from Río Choy (n = 8) and cavefish from Pachón (n = 9) and Los Sabinos (n = 18) used for carotenoid content analysis. Additional cavefish populations referenced in the study are also indicated, with symbols denoting their geographic locations and lineage assignment (Lineage 1 in blue or Lineage 2 in purple). **B**, Representative images of lineage 1 Río Choy surface fish (field collected) and Molino cavefish (laboratory raised) alongside their visceral adipose tissue. **C**, Representative images of lineage 2 Pachón and Los Sabinos cavefish (field collected) alongside their visceral adipose tissue. **D-G**, High-performance liquid chromatography (HPLC) chromatograms displaying the separation of carotenoids extracted from field-collected surface fish and Pachón cavefish gut contents (D, E) and VAT (F, G) respectively. Peaks are identified based on retention time and spectral characteristics of standards (see Supplemental Figure 1). The sharp unlabeled peaks at ∼2min corresponds to the solvent front (injection peak).

Remarkably, even when fed the same diet in the lab, southern cavefish (lineage 2) accumulate high levels of carotenoids in their tissues, producing a vibrant yellow color with no apparent signs of toxicity^10^. Cavefish from both lineages have lost their eyes, yet carotenoid accumulation is not observed in Northern lineage 1 cavefish (Fig 1B,C, C) suggesting that, despite the importance of vitamin A in vision, it is not simply a byproduct of regressive eye evolution and may represent a lineage-specific metabolic adaptation. Here, we investigated the genetic basis and physiological consequences of carotenoid and vitamin A accumulation.

### *Astyanax mexicanus* cavefish consume and accumulate carotenoids in the wild

Because carotenoids are synthesized by plants, algae, and photosynthetic bacteria, they are most abundant in ecosystems with light-driven primary production ^11,12^. Cave environments are perpetually dark, leading to the expectation that they would be naturally low in carotenoids. To assess carotenoid consumption and storage in wild populations of *A. mexicanus*, we analyzed gut contents and visceral adipose tissue (VAT) of field-collected surface fish from the Río Choy River, and southern lineage 2 cavefish from Pachón and Los Sabinos Caves (Fig. 1). Interestingly, the gut contents of both Río Choy surface fish (n = 7) and Pachón cavefish (n = 8) contained β-carotene, β-cryptoxanthin, lutein, and zeaxanthin, suggesting that both populations encounter and ingest similar dietary carotenoids in the wild (Fig. 1D, E, Supplemental Fig. 1). However, only the VAT of Pachón cavefish contained detectable levels of carotenoids, including β-carotene, β-cryptoxanthin, astaxanthin, and other ketocarotenoids (Fig. 1F, G, Supplemental Fig. 2). These results suggest that while cavefish and surface fish consume similar carotenoids in their natural habitat, only cavefish accumulate and store these compounds in their tissues. In line with this observation, the VAT of field-collected Pachón and Los Sabinos appears bright yellow (Fig. 1C).

### Quantitative trait loci (QTL) mapping identifies *bco2a* as a candidate gene underlying cavefish carotenoid accumulation

To investigate the genetic basis of carotenoid accumulation in cavefish, we performed QTL mapping using a binary score of visceral adipose tissue (VAT) color as a proxy for carotenoid level (Fig. 2). Surface/Pachón F2 hybrids fed the same carotenoid-rich diet were categorized as having yellow (n = 23) or non-yellow (n = 102) VAT at the time of dissection. We identified a significant QTL associated with VAT color on chromosome 1 (Fig. 2A). The peak marker explained 19.86% of phenotypic variance (LOD = 6.01; threshold = 4.57, 5% significance, 1000 permutations), and individuals homozygous for the cavefish allele (C/C) were more likely to display yellow VAT (8/10, 80%) compared to heterozygotes (5/50, 10%) or surface fish homozygotes (10/49, 20.4%, Fig 2B,C), supporting a recessive effect of the cavefish allele. A QTL peak was also detected using a semi-quantitative measure of VAT color (normalized red intensity), mapping to the same region with a broader support interval and lower LOD (4.25, Fig. 2D).

**Figure 2.**
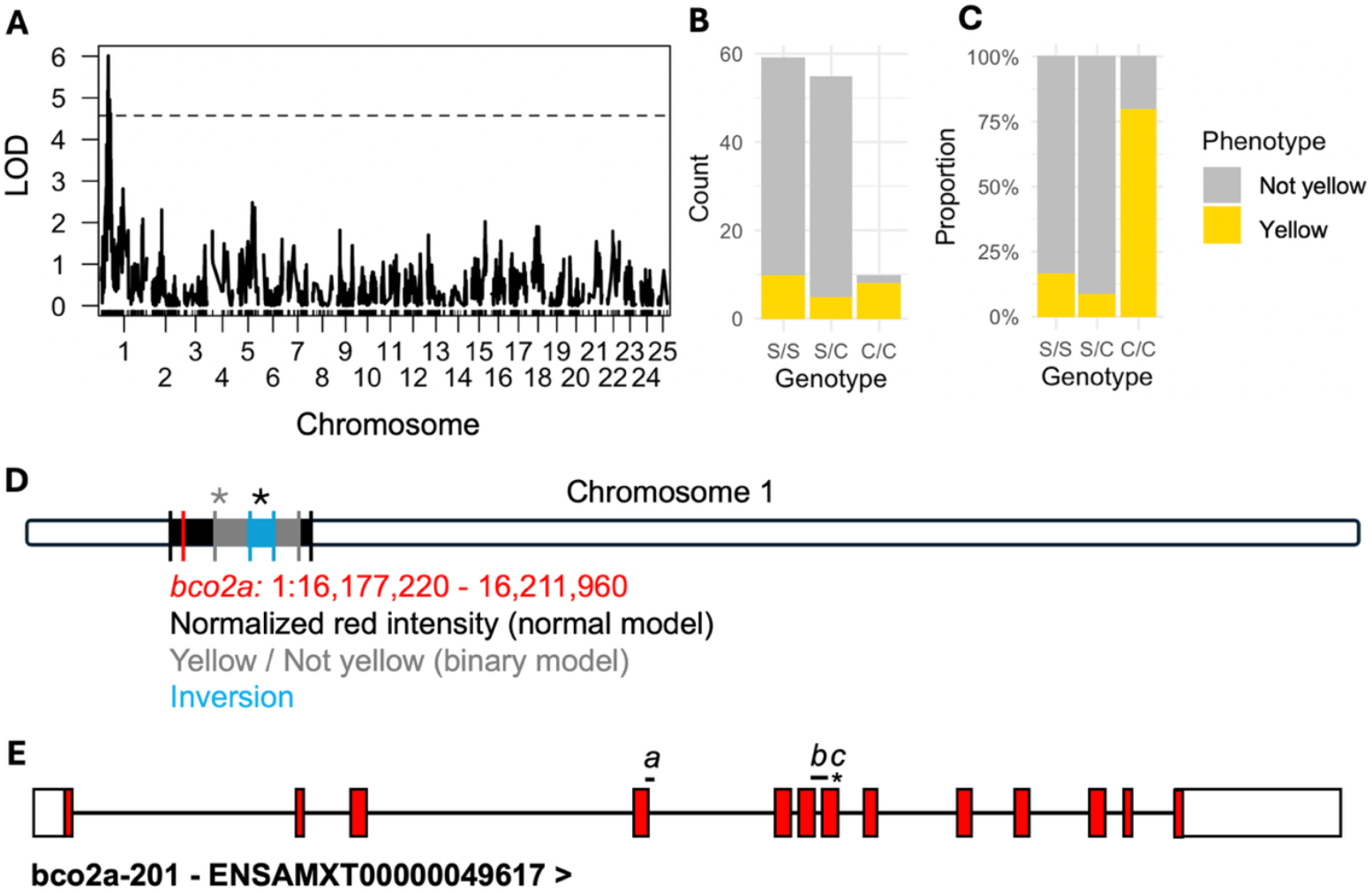
Genetic mapping of visceral adipose tissue (VAT) color in *Astyanax mexicanus* reveals *bco2a* as candidate gene for cavefish carotenoid accumulation. **A**, Genome wide scan for quantitative trait loci associated with visceral adipose tissue color in F2 surface/Pachón hybrids. Tissue was recorded as yellow (n=23) or not yellow (n=102) by visual inspection at the time of dissection. LOD score based on Haley Knott regression and binary model. Dotted line indicates LOD threshold (5%, 4.57) based on 1,000 permutations. Peak marker LOD 6.01 (rs82) explains 19.86% of the variance. **B, C**, Number and proportion of F2 hybrids displaying the indicated genotype/phenotype combinations (S: surface allele, C: Cave allele). **D**, Position of the 1.5 LOD support interval for the VAT color QTL (black: normal model, grey: binary model), *bco2a* locus (red), and Pachón cavefish inversion (blue) on chromosome 1 of the *A. mexicanus* genome (Astymex3_surface). Asterisks indicate peak marker position for QTL (black: normal model, grey: binary model). **E**, *bco2a* gene model showing the positions of two deletions (*a:* variable length deletion that removes 36bp exon 4 and extends into intron 4-5; *b:* 314bp deletion that removes 100bp Exon 6, Intron 6-7, and 70bp Exon 7) and one nonsense mutation (*c:* TAC to TAG in Exon 7) present in field-collected Pachón (n=19) and Tinaja (n=20) cavefish. Open boxes indicate 5’UTR and 3’UTR, red shaded boxes are coding regions of exons, and lines indicate introns.

Notably, genotype ratios at the peak deviated significantly from Mendelian expectations (χ^2^ = 66.22, p = 4.17 × 10^−15^), with underrepresentation of cavefish homozygotes (C/C), a pattern observed across much of chromosome 1 (Supplementary Data 1). This distortion may result from a large chromosomal inversion in Pachón cavefish that suppresses recombination in heterozygotes and reduces recovery of C/C genotypes, possibly through hybrid incompatibility or reduced viability. Nevertheless, the gene *beta-carotene oxygenase 2* (*bco2a*) lies within the 1.5 LOD support interval and encodes a mitochondrial enzyme that cleaves carotenoids, making it a strong mechanistic candidate for the accumulation phenotype observed in cavefish (Fig. 2D).

### Comparative genomics reveals predicted loss-of-function mutations in cavefish *bco2a*

We compared the *bco2a* locus across multiple *A. mexicanus* populations using whole-genome sequencing datasets that included 9–20 individuals per population, spanning both northern (Lineage 1) and southern (Lineage 2) localities (Supplementary Data 2). In Lineage 2 populations (nineteen Pachón, twenty Tinaja, and six of seven Yerbaniz), we identified a variable-length deletion (∼54 bp) affecting 36 bp at the end of exon 4 and 18 bp of intron 4–5, as well as a second 314 bp deletion that removes 100 bp of exon 6, intron 6–7, and 70 bp of exon 7 (Fig. 2E, Supplemental Figure 3). These deletions were absent in all river-dwelling surface fish (Río Choy, Río Mante, Rascón, *n* = 37), including Rascón that genetically cluster with Lineage 2 cavefish. In addition, the deletions were not found in any Lineage 1 cavefish (Molino, Jineo, Vasquez, Escondido, *n* = 37). Lineage 2 cavefish also possessed a single nucleotide polymorphism in exon 7 that introduces a premature stop codon (CHR 1:16203128). All nineteen Pachón individuals were homozygous for the nonsense mutation; eighteen of twenty Tinaja were homozygous and two were heterozygous; and six of seven Yerbaniz were homozygous mutant with one heterozygous (Supplementary Data 2). One out of eighteen Rascón surface fish was heterozygous for the nonsense mutation. In contrast, we found that northern lineage 1 cavefish are predicted to have two coding variants (W96R, V332L) and retain the full-length protein (Supplemental Figure 4).

### Pachón cavefish *bco2a* variants have reduced enzymatic activity

The deletions and nonsense mutation we discovered in southern lineage 2 cavefish are predicted to impact the function of Bco2a as the resulting protein would lack much of the carotenoid oxygenase domain (AA76-589) and all the predicted ferrous iron cation binding sites (AA239, 299, 370, 586, Fig. 3A). Due to the variable length of the deletions, in silico translation of the coding sequences results in most Pachón individuals (17/19) and all Tinaja individuals producing a truncated protein (173AA), with some Pachón cavefish producing a slightly longer version with a gap in the amino acid sequence (AA173-187, Fig. 3A, Supplemental Figure 4).

We compared enzymatic function of the full-length Río Choy surface fish Bco2a and both Pachón cavefish variants by expressing each protein in *E. coli* strains engineered to produce β-carotene or zeaxanthin and quantifying residual carotenoids in cell pellets via HPLC and spectrophotometry (Fig 3B-G). Non-transformed strains contained high carotenoid levels (β-carotene: 1.72 ± 0.46 μg/mL; zeaxanthin: 1.45 ± 0.36 μg/mL), whereas strains expressing mouse Bco2, used as a positive control, showed near complete depletion (β-carotene: 0.034 ± 0.015 μg/mL; zeaxanthin: 0.035 ± 0.021 μg/mL, Fig. 3D, E). Surface fish Bco2a transformants likewise displayed minimal carotenoids (β-carotene: 0.012 ± 0.011 μg/mL; zeaxanthin: 0.01 ± 0.007 μg/mL). In contrast, both Pachón variants accumulated levels comparable to the non-transformed controls (β-carotene: 0.55 ± 0.05 μg/mL; zeaxanthin: 0.68 ± 0.31 μg/mL, Fig. 3D, E).

**Figure 3.**
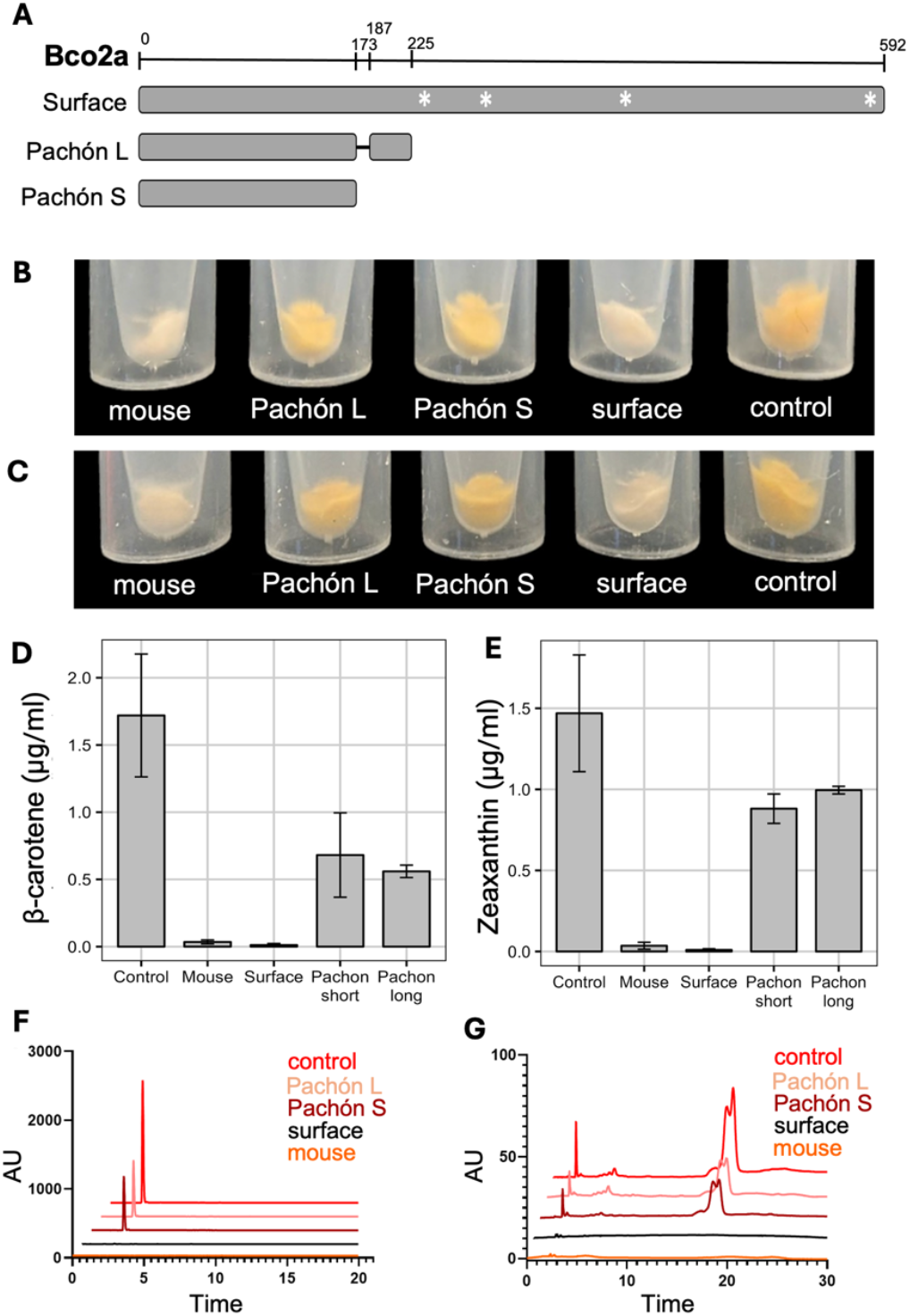
Pachón cavefish have mutations in Bco2a that lead to a truncated protein with reduced oxygenase activity. **A**, Representation of predicted consensus protein sequence alignment comparing Río Choy surface fish (n=9) with Pachón cavefish (n=19). A long and short version of the protein is observed in field collected Pachón, although the short version is most common (n=17/19). Asterisks indicate predicted Fe^2+^ cation binding sites (AA239, 299, 370, 586). **B**,**C**, Representative cell pellets of *E. coli* that produce β-carotene (B) or zeaxanthin (C) after expression of the indicated *bco2a* variant. Control is the non-transfected *E. coli* strains. **D**,**E**, Quantification by spectroscopy of carotenoid concentrations remaining in bacterial cell pellets of E. coli that produce β-carotene (D) or zeaxanthin (E) after expression of the indicated *bco2a* variant. Error bars represent standard deviation; n = 3 replicates per group. **F**,**G**, Quantification by HPLC of carotenoid concentrations remaining in bacterial cell pellets of E. coli that produce β-carotene (F) or zeaxanthin (G) after expression of the indicated *bco2a* variant. AU: area under the curve.

These results suggest that mutations in cavefish *bco2a* produce truncated enzymes lacking carotenoid-cleaving activity. Consistent with this, NCBI annotates the *bco2a* locus as a pseudogene in the Pachón cavefish assembly (AMEX_1.1, GCA_019721115), a designation applied when predicted gene models contain coding sequence disablements like frameshifts of premature stop codons. The *bco2a* paralog, *bco2b*, is unlikely to functionally compensate as it is not expressed in the gut (Supplemental Figure 5). Together, these data indicate that loss of Bco2a function underlies carotenoid accumulation in southern (lineage 2) cavefish.

### Evidence for positive selection on *bco2a*

To determine whether loss-of-function mutations in *bco2a* may be advantageous in the cave environment, we analyzed 5kb windows across the genome for each population using diploS/HIC with demographic models appropriate to cave fish or surface fish. We found that in all populations assayed (Surface: Río Choy, Rascón; Cave: Tinaja, Pachón, Molino) *bco2a* exhibits evidence of either a hard or soft sweep (Supplemental Data 3). This suggests the gene is under selection pressure, albeit potentially from different sources in the cave and surface environments. These patterns indicate environment-specific selection on *bco2a*, with functional alleles favored in surface populations and loss-of-function alleles favored in caves.

### Diet-dependent carotenoid and vitamin A accumulation in cavefish

To investigate how cavefish mutations in *bco2a* influence carotenoid metabolism dynamics, and may thereby provide an advantage in the cave environment, we fed laboratory-raised Pachón cavefish and surface fish a diet supplemented with either low (1,500 IU) or high (4,500 IU) levels of a carotenoid mixture (β-carotene, α-carotene, zeaxanthin, cryptoxanthin, and lutein), or a non-supplemented control diet that is deficient in carotenoids (Fig. 4A). We dissected the fish after one year and found that Pachón visceral adipose tissue (VAT) appeared redder than surface VAT across all supplementation levels consistent with carotenoid accumulation due to loss of Bco2a function (Fig. 4A).

**Figure 4.**
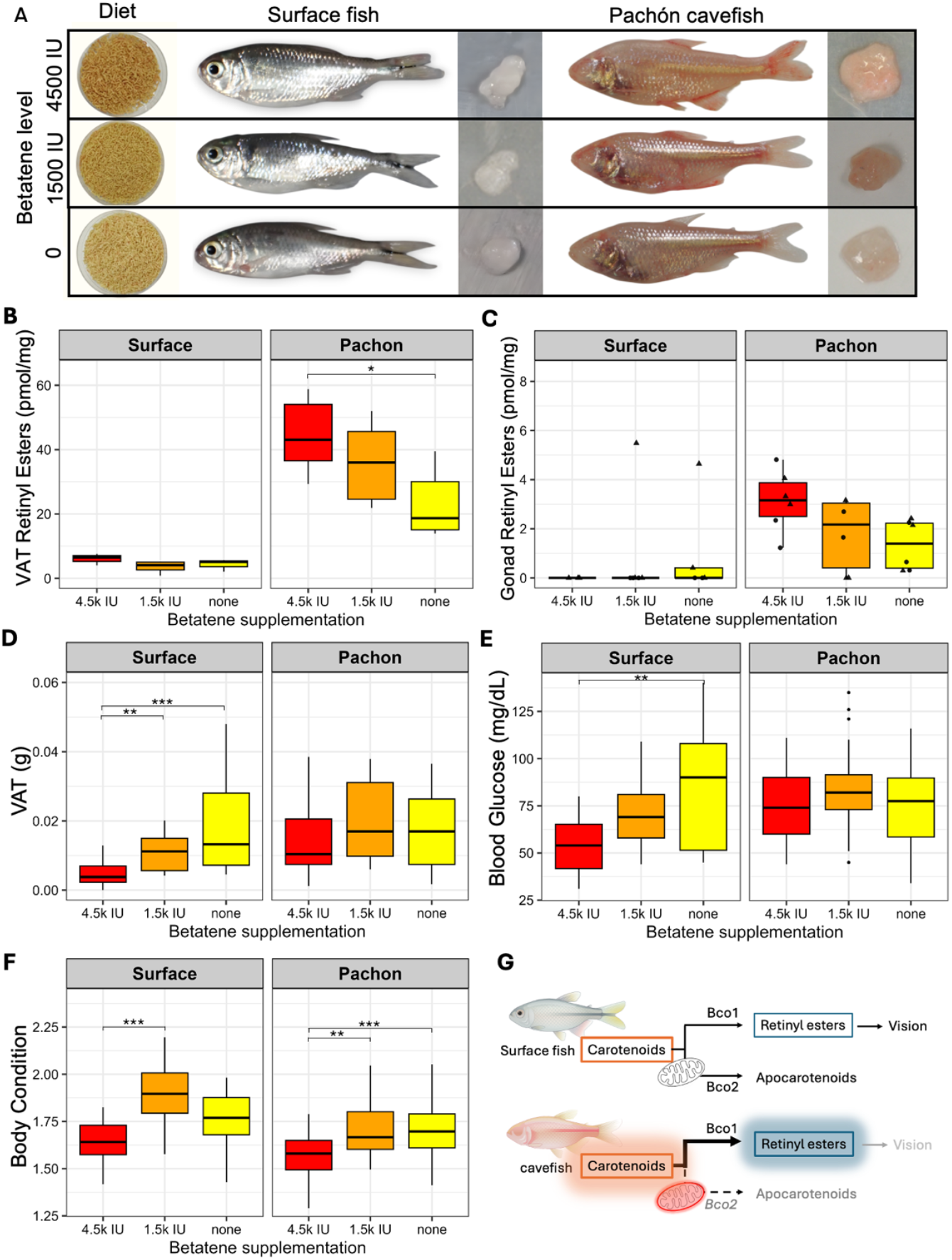
*A. mexicanus* cavefish show diet-responsive retinoid accumulation and metabolic trait stability compared to surface fish. **A**, Representative images of experimental diets supplemented with Betatene, a mixture of β-carotene, α-carotene, zeaxanthin, cryptoxanthin, and lutein. The high-carotenoid diet contained 4,500 IU, the low-carotenoid diet contained 1,500 IU, and the non-supplemented diet contained only base ingredients. Each row shows the diet and a representative surface fish and Pachón cavefish alongside an image of their visceral adipose tissue (VAT) after the one-year trial. **B-F**, Boxplots showing trait measurements after the diet trial. Boxes show the interquartile range (IQR) with the median as a horizontal line; whiskers extend to 1.5× IQR, and outliers are plotted as individual points. **B**, Retinyl ester (RE) levels in VAT. Pachón cavefish had higher RE than surface fish overall (p = 5.33×10^−6^) and accumulated more RE on the high-carotenoid diet compared to non-supplemented (p = 0.0242). RE did not differ significantly among surface fish groups. Sample sizes: Pachón, n = 6, 6, 6; surface, n = 3, 6, 4. **C**, Retinyl ester (RE) levels in gonads. Pachón cavefish had higher RE than surface fish overall (p = 0.0023). RE did not differ significantly among diet groups within either population (Pachón: p = 0.130; Surface: p = 0.488). Sample sizes: Pachón, n = 6, 6, 6; surface, n = 3, 6, 5. Points represent individual samples of the indicated gonad type (male testis: triangle, female ovary: circle). **D**, VAT mass. In surface fish, VAT mass was higher on the non-supplemented vs. high-carotenoid diet (p = 0.0001), and higher on the low-vs. high-carotenoid diet (p = 0.0078). VAT mass did not differ significantly among Pachón groups. Sample sizes: Pachón, n = 26, 23, 30; surface, n = 23, 16, 14. **E**, Fasting blood glucose. Surface fish had higher glucose on the non-supplemented diet compared to the high-carotenoid diet (p = 0.001). Glucose did not differ significantly among Pachón groups. Sample sizes: Pachón, n = 25, 23, 30; surface, n = 22, 16, 12. **F**, Body condition factor (BCF). In surface fish, BCF was greater on the low-vs. high-carotenoid diet (p = 0.0001). In Pachón fish, BCF was greater on the non-supplemented (p = 0.014) and low-carotenoid diets (p = 0.049) compared to the high-carotenoid diet. Sample sizes: Pachón, n = 26, 23, 30; surface, n = 23, 16, 14. Statistical comparisons were performed using Kruskal–Wallis tests with Dunn’s post-hoc tests. Significance is indicated as: *p < 0.05, **p < 0.01, ***p < 0.001. **G**, Comparative model of carotenoid metabolism. Loss of Bco2a activity in cavefish may increase substrate availability for Bco1, resulting in elevated carotenoid and retinyl ester levels. Combined with absence of vision this may lead to systemic vitamin A accumulation and associated metabolic changes. Created with BioRender.com.

We found that retinyl ester (RE), the primary storage form of vitamin A, was significantly elevated in Pachón cavefish across all diets compared to surface fish (Fig. 4B, 34.5 ± 3.2 vs. 4.71 ± 1.1 pmol/mg; Wilcoxon p = 5.33 × 10^−6^), with diet-dependent differences in RE accumulation (Kruskal-Wallis p = 0.0268). RE levels in Pachón cavefish were highest on the high-carotenoid diet (45.4 ± 4.1 pmol/mg), intermediate on the low-carotenoid diet (36.6 ± 2.9), and lowest on the non-supplemented diet (24.6 ± 3.7; Dunn’s p = 0.0242). Strikingly, even without carotenoid supplementation, Pachón cavefish stored over five times more RE in their VAT than surface fish (24.6 ± 3.7 vs. 4.71 ± 1.1 pmol/mg), underscoring their enhanced capacity for vitamin A storage (Fig. 4B). This trend extended to the gonads, where Pachón fish had significantly higher RE levels than surface fish (Fig. 4C, 2.58 ± 0.19 vs. 0.75 ± 0.14 pmol/mg; Wilcoxon p = 0.0023). These findings suggest that loss of Bco2a activity in Pachón cavefish may increase the pool of provitamin A carotenoids available for cleavage by Bco1, thereby promoting vitamin A synthesis (Fig. 4G). Coupled with the absence of visual demand for vitamin A, this shift may favor retinoid storage across multiple tissues (Fig. 4G).

### Cavefish exhibit resilience to carotenoid supplementation extremes

Pachón cavefish exhibited greater tolerance to dietary carotenoid variation than surface fish over the one-year feeding trial. Surface fish fed the non-supplemented diet, which is deficient in pro-vitamin A, frequently developed symptoms consistent with bacterial infection including raised scales, abdominal distension, gas bubbles in the gastrointestinal tract, impaired swimming, inflamed fins, and scale loss. Some of these phenotypes align with hallmark signs of immune dysfunction associated with vitamin A deficiency ^13–15^ and were never observed in cavefish under any diet. Reflecting this physiological vulnerability, more surface fish on the non-supplemented diet either died or had to be removed from the diet trial (27.5%), whereas no mortality or signs of distress occurred in cavefish on the same diet. Mortality remained low in the other treatment groups for both populations, though slightly elevated in surface fish fed the high-carotenoid diet (Surface: low = 2.5%, high = 15%; Cavefish: low = 7.5%, high = 5%).

Carotenoid intake influenced multiple metabolic traits in surface fish but had minimal effects in Pachón cavefish. In surface fish, visceral adipose tissue (VAT) mass varied significantly across diets (Kruskal-Wallis: χ^2^(2) = 17.161, p = 0.00019), with greater fat accumulation in fish fed the non-supplemented and low-carotenoid diets compared to the high-carotenoid group (Fig. 4D, Dunn’s test: p = 0.00043). In contrast, VAT mass in Pachón cavefish did not differ significantly across diets (χ^2^(2) = 2.4063, p = 0.3003). Fasting blood glucose levels were also diet-sensitive in surface fish, with significantly higher glucose observed in the non-supplemented group compared to the high-carotenoid group (Fig. 4E, 90.5 vs. 54.4 mg/dL; p = 0.00354). No such dietary differences were detected in Pachón cavefish (high: 76.1, low: 84.2, non-supplemented: 74.8 mg/dL). However, sex differences emerged in cavefish, with males exhibiting significantly higher glucose levels than females across all diets (two-way ANOVA: F(1,72) = 10.693, p = 0.002).

Body condition factor (BCF) was diet-dependent in both populations (Fig. 4F). Surface fish fed the low-carotenoid diet had higher BCF than those on the high-carotenoid diet (p = 0.00010), and Pachón cavefish fed either the low- or non-supplemented diets had higher BCF than those on the high-carotenoid diet (p = 0.049 and 0.014, respectively). Male surface fish also had lower BCF than females across all diets (p = 0.006), whereas sex had no significant effect on BCF in cavefish. Because tanks were provided equal rations that were fully consumed, we infer that these differences in metabolic traits reflect physiological responses to carotenoid supplementation rather than variation in food intake.

Together, these findings indicate that surface fish exhibit diet-dependent shifts in fat storage, glucose metabolism, and body condition, while Pachón cavefish maintain relative physiological stability across diets potentially reflecting an evolved resilience to variation in carotenoid availability.

## Discussion

Our findings reveal that some cave-adapted populations of *A. mexicanus* have evolved altered carotenoid metabolism via loss of the mitochondrial carotenoid oxygenase, Bco2a. This claim is supported by genetic, comparative, and biochemical evidence. QTL mapping identified a significant association between carotenoid-rich visceral adipose and the *bco2a* locus; comparative genomics across multiple wild populations revealed coding mutations predicted to truncate Bco2a and eliminate much of the carotenoid oxygenase domain; and functional assays confirmed that these variants lack carotenoid-cleaving activity. The *bco2a* nonsense mutation found in southern lineage cavefish was detected at low frequency in the sister surface fish population, Rascón. This allele distribution, combined with signatures of selection at the *bco2a* locus, suggests that impaired carotenoid degradation is advantageous in the cave environment and arose from standing genetic variation.

We found that cavefish have elevated vitamin A storage and display resilience to diets with excess and deficient carotenoids. Mechanistically, loss of Bco2a in cavefish could increase the pool of substrates available for retinoid production by Bco1 (Fig. 4G). In mammals, BCO1 and BCO2 operate in distinct cellular compartments, preventing competition: BCO1 cleaves provitamin A carotenoids in the cytosol, whereas BCO2 resides in the mitochondria and can cleave pro- and non-provitamin A carotenoids^16^. The relevance of BCO2 to vitamin A homeostasis is therefore debated. However, BCO2 knockout mice accumulate higher retinoid stores on diets containing only the provitamin A, β-carotene, suggesting that BCO2 activity may indirectly shape vitamin A balance by altering substrate availability for BCO1^4^. Fish are also known to use a wider range of carotenoids as sources of vitamin A compared to mammals, including xanthophylls like astaxanthin, zeaxanthin and lutein^17–19^. Thus, loss of mitochondrial Bco2a in cavefish could increase the Bco1 substrate pool. Combined with the lack of demand for retinal as a visual chromophore in eyeless cavefish, systemic elevation of vitamin A and carotenoids may result (Fig. 4G).

In mammals, BCO2 resides at the inner mitochondrial membrane^20^, where it clears carotenoids that would otherwise disrupt membrane potential and respiratory function due to their rigid, electron-accepting polyene backbones^6,21^. Accordingly, Bco2 knockout mice exhibit mitochondrial dysfunction, elevated ROS production, and changes in hepatic gene expression indicative of oxidative stress^5,6^. Bco2 knockout mice also display metabolic abnormalities like fatty liver, increased fasting blood glucose, and elevated circulating free fatty acids^5,6^. Interestingly, cavefish exhibit parallel traits like fatty liver^22^, elevated blood glucose^23^, and increased fat accumulation^24^, yet avoid mammalian like pathology^25^, consistent with a derived metabolic state that is advantageous in nutrient-scarce caves. Although mitochondrial function has not been directly measured in cavefish, loss of Bco2a is predicted to alter mitochondrial metabolism with organism-wide consequences for energy balance, and may contribute to some cavefish metabolic adaptations.

Carotenoids can act as antioxidants or pro-oxidants depending on redox context^26,27^. Therefore, carotenoid accumulation has potentially unfavorable costs. Notably, under high-oxygen laboratory conditions, cavefish exhibit elevated ROS in the brain and gut compared to surface fish^28^, indicating that carotenoid accumulation may impose oxidative costs in oxygen-rich settings. However, in the natural hypoxic cave habitat ^29,30^, these effects may be buffered or mitigated. In addition, cavefish may offset the risks of carotenoid accumulation through elevated levels of other antioxidants and changes in gene expression; they exhibit increased glutathione and vitamin C levels and upregulate ROS-scavenging enzymes compared to surface fish^31,32^. Rather than metabolizing carotenoids via mitochondrial catabolism, cavefish may retain them as functional components of a broader antioxidant network that protects against oxidative stress under variable oxygen conditions^30^.

Carotenoid accumulation in cavefish could also provide reproductive benefits. Cavefish spawn during the dry season, when oxygen levels in caves are lowest^33^. Maternal provisioning of carotenoids to eggs may help buffer developing embryos and larvae against oxidative stress, both under chronic hypoxia and during transient spikes in oxygen that occur with seasonal flooding. Supporting this idea, cavefish embryos and larvae show improved growth in the laboratory when reared under hypoxic conditions compared to oxygenated conditions^29^, suggesting their early development is physiologically tuned to low-oxygen environments. This strategy parallels observations in other fish species reproducing in low-oxygen habitats, where yolk-enriched carotenoids function as both nutritional reserves and redox modulators^34–36^.

Together, our results indicate that some *A. mexicanus* cavefish have adaptively rewired carotenoid metabolism through the loss of Bco2a, enabling them to retain and utilize carotenoids in ways that may support survival and reproduction in extreme environments. This rewiring is supported by genetic, physiological, and ecological evidence, and provides a striking example of how vertebrates can modify canonical nutrient pathways to accommodate novel ecological challenges.

## Methods

### Field-collection of *Astyanax mexicanus* VAT samples

Adult *Astyanax mexicanus* were collected from four populations in the Sierra de El Abra region in northeastern Mexico: Pachón (cave), Los Sabinos (cave), Río Choy (surface), and Río Subterráneo (hybrid cave-surface). Twenty adult individuals per population (n = 80 total) were sampled in early April 2023 using small mesh nets (25 cm × 20 cm) and, in the case of surface populations, cast nets. All sampling procedures were carried out under permits SPARN/DGVS/03371/23 and SPARN/DGVS/03307/23, issued by the General Directorate of Wildlife (Dirección General de Vida Silvestre, SEMARNAT). Collection of *Astyanax mexicanus* (referred to as *A. jordani* in the permit) was authorized from cave systems in the states of San Luis Potosí and Tamaulipas. Standard collection techniques were employed in accordance with Mexican wildlife scientific collection regulations (NOM-059-SEMARNAT-2010). Following capture, fish were temporarily housed in aerated buckets and transported to a nearby field station. On the same day, fish were euthanized using buffered MS-222 at a concentration of 200 mg/L, followed immediately by dissection. Standard length, body mass, and sex were recorded for each individual post-mortem, and the visceral adipose tissue (VAT) and gut contents were collected. Samples were flash-frozen in liquid nitrogen, transported on dry ice, and stored at –80°C until analysis. Samples were chosen for analysis based on adequate sample amount.

### Carotenoids extraction from fish tissue samples

Deep frozen fish samples were homogenized in 250 µL phosphate saline buffer (pH 7.5). Then 250 µL of methanol was added and the suspension was vortexed. After addition of 500 µL acetone, the suspension was again vortexed and kept for 5 minutes on ice. 200 µL of diethyl ether and 400 µL hexane were added to extract carotenoids. Phase separation was achieved by centrifugation at 3,000 x g for 1 minute. The upper organic layer was transferred to a new tube. The extraction was repeated once with 500 µL hexane. The combined organic phases were vacuum dried in a Speedvac (Eppendorf, Hauppauge, NY) and the debris were dissolved in 200 µL hexane: ethyl acetate (80:20 v/v) for HPLC analysis.

### High-performance liquid chromatography (HPLC) analysis

Extracted lipids were separated by HPLC performed on a 1200 Agilent HPLC series equipped with a diode array detector and normal-phase Restek silica column (4.6 mm ID x 150 mm with 5 µm packing, Bellefonte, PA, USA). Chromatographic separation was achieved with isocratic flow of hexane: ethyl acetate (80:20 v/v) with a flow rate of 1.4 mL/min. The system was scaled with known amounts of authentic standards.

### Quantitative trait loci mapping

Sequencing data from surface/Pachón F1 and F2 hybrids used in a previous QTL study^37^ were reanalyzed to generate an updated QTL linkage map aligned with the most recent chromosome-level *A. mexicanus* reference genome (Astymex3_surface). The mapping population (n = 219) consisted of individuals from three clutches produced by crossing F1 surface/Pachón hybrid siblings. After removing PCR duplicates, aligned reads were used to call SNPs in all F1 and F2 fish using freebayes v1.3.2 (Garrison and Marth. 2012). Variant calls were filtered with bcftools v1.14 (Danecek *et al*. 2021) DP>5 and GQ>20. We then called the genotypes of the F1 and F2 variants sites, using freebayes on 8 publicly available Río Choy surface fish and Pachón cavefish genomes (four individuals per population; SRR1575280, SRR1575281, SRR1575282, SRR1575283, SRR1575270, SRR1575271, SRR1575272, SRR1575273).

Using custom variant filtering scripts only markers showing fixed differences between the four Río Choy surface fish and four Pachón cavefish genomes were retained. Similarly, we further narrowed the markers to variants that exhibiting heterozygosity without evidence of homozygosity in surface/Pachón F1 hybrids. Then we extracted only the F2 individuals and their genotypes and we removed duplicate markers with rQTL ‘findDupMarkers()’ and markers with significant departures from the 1:2:1 expected ratios. To create a linkage map we ordered the markers based on their relative physical positions, according to their genome coordinates and transformed their physical distances into recombination distances by assuming a constant recombination rate across the genome resulting in a linear relationship between bp and cM distances. We used the *A. mexicanus* estimated size in cM from Carlson *et al*. (2014) at 2110.7 cM and in bp by Warren *et al*. (2021) at 1.335 Gb. Finally, our linkage map contained 3,667 high-confidence markers.

Phenotypic data for the F2 surface/Pachón hybrid mapping population were obtained from previously published studies^37^. Fish were reared at densities below one adult per 2 L until 14 months of age, after which they were transferred to individual 1.5 L tanks. Each fish was fed three pellets (approximately 6 mg) of New Life Spectrum TheraA+ small fish formula once daily for a period of 4 to 6 months. At the end of the feeding period, fish were euthanized using 1400 ppm Tricaine and dissected. Visceral adipose tissue (VAT) color was scored visually as either yellow or non-yellow at the time of dissection. In addition, VAT was photographed using a Canon Powershot D12 digital camera.

For quantitative color analysis, mean RGB values were extracted from a 16 × 16 pixel region of the VAT and a matching region of an 18% gray card present in the same image. Standardized RGB values were calculated by dividing each VAT color channel by the corresponding value from the gray card (e.g., Rstand = mean R_sample / mean R_graycard). Normalized red intensity (nIR) was then calculated as: nIR = Rstand / (Rstand + Gstand + Bstand). This metric has previously been used as an estimate of carotenoid-based coloration in stickleback pelvic spines^38^. VAT color was visually scored for F2 hybrids with clearly discernible VAT (n = 125/219), and normalized red intensity was calculated for those with usable images containing both VAT and a grayscale card (n = 62/125).

QTL analysis was conducted using the R/qtl package^39^. For visual VAT color (scored as a binary trait: 0 = non-yellow, 1 = yellow), we used Haley–Knott regression under a binary model. For quantitative measurements, we applied a normal model. Statistical significance was assessed using 1,000 genome-wide permutations to determine the 95th percentile of the maximum penalized LOD score (scanone function). Confidence intervals for the QTL were estimated using the 1.5-LOD support interval (lodint) extended to the nearest genotyped marker.

### Comparative genomics and population genetic metrics

*Bco2a* (ENSAMXG00000003177, APWO02000033.1: 978,225-1,011,843) genomic DNA and predicted cDNA sequence alignments were obtained from variant call format (VCF) files of surface fish (Río Choy, Mante, Rascón), lineage 1 cavefish (Molino, Jineo, Escondido, Vasquez), and lineage 2 cavefish (Tinaja, Pachón, Yerbaniz) genomes aligned to a surface fish reference genome^40^ (Supplementary Data 2). Transcript ENSAMXT00000049617.1 was used as a reference for intron-exon structure.

We aligned *bco2a* cDNA sequences of individuals from only one population (Surface n=9, Pachón n=19, Molino n=15, Tinaja n=20) to extract a consensus cDNA sequence for each population. The consensus sequences were translated, and the predicted peptide sequences were aligned using SnapGene Global alignment (Supplemental Fig. 4).

To evaluate evidence of positive selection, we realigned population genomic data from Moran 2023^9^ (https://doi.org/10.5061/dryad.3xsj3txmf) to GCF_023375975.1 following identical methods and filtering cut-offs as described therein. We simulated population genetic data from demographic data to represent a surface population and, separately, a cave population in discoal and used this simulated data to generate two sets of feature vectors in diploS/HIC from training and testing the convoluted neural network. We generated predictions of selection for 5kb windows across the genome, classifying each window as neutral, soft, hard, softLinked or hardLinked.

### Statistical analysis

All measurements were taken from independent individual fish or independent bacterial cultures, with no repeated measurements from the same sample. All statistical tests were two-sided. Normality of distributions was assessed using Shapiro–Wilk tests; when assumptions were not met, nonparametric alternatives (Wilcoxon rank-sum, Kruskal–Wallis) were applied.

### Cell-based assay to measure carotenoid oxygenase activity

#### Plasmid generation and bacterial transformation

*Astyanax mexicanus* surface fish Bco2a predicted peptide sequence (UniProt ID A0A3B1IL41, ENSAMXT00000049617, Astyanax_mexicanus-2.0) and Pachón cavefish peptide sequences deduced from field-collected DNA samples (Supplementary Fig. 4) were used to generate codon-optimized genes for *E. coli* expression using GenScript’s proprietary optimization algorithm. The optimized genes were synthesized and subcloned into the pMAL-c4x expression vector via EcoRI and BamHI restriction sites to generate an MBP-tagged fusion construct (Supplemental Data 4-6). Mouse *bco2a* cDNA cloned into pMAL-C5X vector was generated in a previous study^41^. Carotenoid-producing *E. coli* strains generated in a previous study^42^ were grown to early log phase (OD600 = 0.3) in LB medium containing 15 μg/mL chloramphenicol. Cells were harvested by centrifugation, treated with ice-cold 0.1 M CaCl2, and transformed with the plasmids described above using heat shock (42°C, 45 seconds).

#### Enzyme assay

Transformed bacteria were cultured in LB medium containing antibiotics to OD600 = 1.0. Carotenoid production was induced by temperature shift to 25°C for 1 hour, followed by induction of *bco2a* expression with 0.3 mM IPTG and supplementation with 25 mg/L ferrous sulfate. After 16 hours of expression at 25°C, cells were harvested by centrifugation at 4000 × g for analysis.

This process was repeated three times for each *bco2a* variant. From each replicate, cell pellets were processed for both spectrophotometric and HPLC analysis. For spectrophotometric analysis, cell pellets were resuspended in 300 μL cold methanol, followed by addition of 300 μL PBS (1:1 v/v) and 600 μL acetone. Carotenoids were extracted into 500 μL hexanes through phase separation. This extraction was repeated twice, and the carotenoids were reconstituted in acetone.

#### Spectrophotometric and HPLC Quantification

One sample from each replicate was analyzed by spectrophotometry, measuring absorbance across wavelengths from 400-500 nm. The averaged values from these measurements were used to calculate carotenoid concentrations using the Beer-Lambert law. Absorbance measurements were taken at 450 nm for beta-carotene and 478 nm for zeaxanthin using extinction coefficients of 139,500 M^−1^cm^−1^ and 144,000 M^−1^cm^−1^, respectively. The samples were diluted with acetone (1.5x dilution factor), and concentrations were calculated using the equation: c = (A/ε)*d, where c is the molar concentration, A is the absorbance, ε is the extinction coefficient, and d is the dilution factor. Final concentrations were expressed in µg/mL after conversion using the molecular weights of beta-carotene (536.87 g/mol) and zeaxanthin (568.88 g/mol). The same harvested cells used for spectrophotometric analysis were subsequently analyzed by HPLC as described above to provide detailed profiles of carotenoid content.

### Carotenoid-variable diet trial

#### Diet Preparation and Carotenoid Supplementation

Experimental diets were formulated to test the effects of carotenoid consumption on metabolism and physiology. Three diet treatments were prepared: The non-supplemented diet contained only carotenoids from the diet ingredients [Casein (Carolina) 48.6 g/100 g; Wheat starch (Red Lantern) 18 g/100 g; Rice bran (NOW Foods) 8 g/100 g; Methyl cellulose (Pure Organic Ingredients) 4 g/100 g; Cellulose (AllergyResearchGroup) 10 g/100 g; Vitamin mix, 1 g/100 g; Vitamin C, ascorbic acid (Carolina) 0.4 g/100 g; Mineral premix (Florida Aqua Farms) 1 g/100 g; Menhaden fish oil (Aquatic Nutrition) 3 g/100 g; Soybean oil (US Organic) 6 g/100 g], without additional supplementation. The low carotenoid diet was supplemented with 1500 IU Betatene (Swanson) per 100 g food, and the high carotenoid diet was supplemented with 4500 IU Betatene per 100 g food. Betatene capsules contained a mixture of β-carotene (14,288 μg), α-carotene (450 μg), zeaxanthin (90 μg), cryptoxanthin (105 μg), and lutein (67 μg) per capsule, derived from *Dunaliella salina* marine algae. The dry and wet ingredients of the diet were prepared separately before mixing. Betatene was incorporated into the wet (oil) portion before being combined with dry components to ensure even carotenoid distribution. The mixed diets were pressed through a gauge syringe, dried overnight at room temperature, and cut into thin pellets.

#### Fish Maintenance and Experimental Feeding Regimen

All procedures were approved by the institutional animal care and use committee (IACUC protocol 21-01-1124). Laboratory-spawned surface fish and Pachón cavefish were reared at 22– 24°C under a 14L:10D light cycle on a recirculating water system (conductivity ∼1200μS, pH ∼7.4). Prior to dietary manipulation, fish were fed live Artemia and New Life Spectrum Thera A+ pellets. Fish of the same clutches were assigned to their respective diets at 8 months of age and maintained on the diets for 12 months. Each diet group consisted of at least 40 individuals per population, housed in 30 L tanks at a stocking density of 40–60 fish per tank. Fish were fed twice daily, approximately one tablespoon of food per tank. Feeding was closely monitored to ensure that all food was consumed within 5–10 minutes and to verify that all fish were actively feeding. Fish that exhibited signs of distress were removed from the feeding trial.

#### Dissections and Tissue Collection

At the conclusion of the feeding trial, fish were euthanized in tricaine methanesulfonate (MS-222) at a concentration of 0.02% (200 mg/L), buffered with sodium bicarbonate, following ethical guidelines. Blood samples were obtained from the caudal vein for glucose analysis, and whole-body imaging was conducted before dissection. Internal organs were removed, imaged, weighed, and flash-frozen in liquid nitrogen for further biochemical analyses.

### Use of AI tools

Portions of the text were refined using ChatGPT (GPT-5, OpenAI, 2025) to improve clarity. All scientific content was generated by the authors.

## Acknowledgments

Suzanne McGaugh for providing sequence alignments and population metrics. Marc Bauhus, Ulla Pebesma, Michelle Borgers, and Ana Santacruz for their support during field collections, dissections and travel. Research equipment utilized in this publication was partly supported by the National Institute of General Medical Sciences of the National Institutes of Health under grant number P30 GM145646.

## Author Contributions

DPZ and MRR contributed to experimental design and execution, data analysis, figure preparation, and drafting of the manuscript; SB and JVL contributed HPLC analysis; FL contributed QTL mapping. All authors contributed to editing and approved the manuscript.

## Data availability

All data supporting the findings in the study are provided in the article and supplementary information. Raw sequencing data used for QTL mapping have been deposited in the NCBI Sequence Read Archive under accession number: SRP046999, Bioproject: PRJNA260715. Sequencing data used for population metrics and alignments is available under PRJNA558458. The surface fish *Astyanax mexicanus* genome assembly, Astyanax_mexicanus-2.0 (accession # GCF_000372685.2) is available on NCBI.

## References

1. Huang, Z., Liu, Y., Qi, G., Brand, D. & Zheng, S. Role of Vitamin A in the Immune System. J Clin Med 7, 258 (2018).

2. Penniston, K. L. & Tanumihardjo, S. A. The acute and chronic toxic effects of vitamin A. Am J Clin Nutr 83, 191–201 (2006).

3. Amengual, J. et al. A mitochondrial enzyme degrades carotenoids and protects against oxidative stress. The FASEB Journal 25, 948–959 (2011).

4. Amengual, J. et al. Two Carotenoid Oxygenases Contribute to Mammalian Provitamin A Metabolism. Journal of Biological Chemistry 288, 34081–34096 (2013).

5. Wu, L. et al. Lack of β, β-carotene-9’, 10’-oxygenase 2 leads to hepatic mitochondrial dysfunction and cellular oxidative stress in mice. Mol Nutr Food Res 61, (2017).

6. Amengual, J. et al. A mitochondrial enzyme degrades carotenoids and protects against oxidative stress. FASEB J 25, 948–59 (2011).

7. Turner, B.J. Biology and Evolution of the Mexican Cavefish. Edited by Alex C. Keene, Masato Yoshizawa, and Suzanne E. McGaugh. Academic Press. Amsterdam (The Netherlands) and Boston (Massachusehs): Elsevier. $99.95. xiv + 403 p.; ill.; index. ISBN: 978-0-12-802148-. Q Rev Biol 92, 485–486 (2017).

8. Herman, A. et al. The role of gene flow in rapid and repeated evolution of cave-related traits in Mexican tetra, Astyanax mexicanus. Mol Ecol 27, 4397–4416 (2018).

9. Moran, R. L. et al. Selection-driven trait loss in independently evolved cavefish populations. Nat Commun 14, 2557 (2023).

10. Riddle, M. R. et al. Genetic architecture underlying changes in carotenoid accumulation during the evolution of the blind Mexican cavefish, Astyanax mexicanus. J Exp Zool B Mol Dev Evol 334, 405–422 (2020).

11. Maoka, T. Carotenoids as natural functional pigments. J Nat Med 74, 1–16 (2020).

12. Nisar, N., Li, L., Lu, S., Khin, N. C. & Pogson, B. J. Carotenoid Metabolism in Plants. Mol Plant 8, 68–82 (2015).

13. Jiang, W.-D. et al. Inconsistently impairment of immune function and structural integrity of head kidney and spleen by vitamin A deficiency in grass carp (Ctenopharyngodon idella). Fish Shellfish Immunol 99, 243–256 (2020).

14. Jiang, W.-D. et al. Vitamin A deficiency impairs intestinal physical barrier function of fish. Fish Shellfish Immunol 87, 546–558 (2019).

15. Shastak, Y. & Pelletier, W. Captivating Colors, Crucial Roles: Astaxanthin’s Antioxidant Impact on Fish Oxidative Stress and Reproductive Performance. Animals 13, 3357 (2023).

16. Palczewski, G., Amengual, J., Hoppel, C. L. & Lintig, J. Evidence for compartmentalization of mammalian carotenoid metabolism. The FASEB Journal 28, 4457–4469 (2014).

17. Kaisuyama, M. & Matsuno, T. Carotenoid and vitamin A, and metabolism of carotenoids, β-carotene, canthaxanthin, astaxanthin, zeaxanthin, lutein and tunaxanthin in tilapia Zlapia nilotica. ComparaHve Biochemistry and Physiology Part B: ComparaHve Biochemistry 90, 131–139 (1988).

18. Gross, J. & Budowski, P. Conversion of carotenoids into vitamins A1 and A2 in two species of freshwater fish. Biochemical Journal 101, 747–754 (1966).

19. Moren, M., Næss, T. & Hamre, K. Conversion of β-carotene, canthaxanthin and astaxanthin to vitamin A in Atlantic halibut (Hippoglossus hippoglossus L.) juveniles. Fish Physiol Biochem 27, 71–80 (2002).

20. Palczewski, G., Amengual, J., Hoppel, C. L. & Lintig, J. Evidence for compartmentalization of mammalian carotenoid metabolism. The FASEB Journal 28, 4457–4469 (2014).

21. Demmig-Adams, B. & Adams, W. W. Antioxidants in Photosynthesis and Human Nutrition. Science (1979) 298, 2149–2153 (2002).

22. Aspiras, A. C., Rohner, N., Martineau, B., Borowsky, R. L. & Tabin, C. J. Melanocortin 4 receptor mutations contribute to the adaptation of cavefish to nutrient-poor conditions. Proceedings of the NaHonal Academy of Sciences 112, 9668–9673 (2015).

23. Riddle, M. R. et al. Insulin resistance in cavefish as an adaptation to a nutrient-limited environment. Nature 555, 647–651 (2018).

24. Xiong, S. et al. Enhanced lipogenesis through Pparγ helps cavefish adapt to food scarcity. Current Biology 32, 2272-2280.e6 (2022).

25. Riddle, M. R. et al. Insulin resistance in cavefish as an adaptation to a nutrient-limited environment. Nature 555, 647–651 (2018).

26. Young, A. J. & Lowe, G. M. Antioxidant and Prooxidant Properties of Carotenoids. Arch Biochem Biophys 385, 20–27 (2001).

27. Edge, R. & Truscoh, T. Singlet Oxygen and Free Radical Reactions of Retinoids and Carotenoids—A Review. AnHoxidants 7, 5 (2018).

28. Lloyd, E. et al. Elevated DNA Damage without signs of aging in the short-sleeping Mexican Cavefish. Preprint at 10.1101/2024.04.18.590174 (2024).

29. van der Weele, C. M. & Jeffery, W. R. Cavefish cope with environmental hypoxia by developing more erythrocytes and overexpression of hypoxia-inducible genes. Elife 11, (2022).

30. Boggs, T. & Gross, J. Reduced Oxygen as an Environmental Pressure in the Evolution of the Blind Mexican Cavefish. Diversity (Basel) 13, 26 (2021).

31. Terzi, A., Ngo, K. J. & Mourrain, P. Phylogenetic conservation of the interdependent homeostatic relationship of sleep regulation and redox metabolism. Journal of ComparaHve Physiology B 194, 241–252 (2024).

32. Medley, J. K. et al. The metabolome of Mexican cavefish shows a convergent signature highlighting sugar, antioxidant, and Ageing-Related metabolites. Elife 11, (2022).

33. Espinasa, L., Rohner, N. & Rétaux, S. Reproductive seasonality of Astyanax mexicanus cavefish. Zool Res 44, 698–700 (2023).

34. Pike, T. W., Blount, J. D., Lindström, J. & Metcalfe, N. B. Dietary carotenoid availability influences a male’s ability to provide parental care. Behavioral Ecology 18, 1100–1105 (2007).

35. Wilkins, L. G. E., Marques da Cunha, L., Glauser, G., Vallat, A. & Wedekind, C. Environmental stress linked to consumption of maternally derived carotenoids in brown trout embryos (Salmo truOa). Ecol Evol 7, 5082–5093 (2017).

36. Williams, B. L., Pintor, L. M., Toomey, M. B. & Gray, S. M. Intraspecific Variation in Color and Carotenoids across Environmental Extremes in an African Cichlid. Ecological and EvoluHonary Physiology 98, 57–69 (2025).

37. Riddle, M. R. et al. Genetic mapping of metabolic traits in the blind Mexican cavefish reveals sex-dependent quantitative trait loci associated with cave adaptation. BMC Ecol Evol 21, 94 (2021).

38. Amundsen, C., Nordeide, J., Gjøen, H., Larsen, B. & Egeland, E. Conspicuous carotenoid- based pelvic spine ornament in three-spined stickleback populations—occurrence and inheritance. PeerJ 3, e872 (2015).

39. Broman, K. W., Wu, H., Sen, S. & Churchill, G. A. R/qtl: QTL mapping in experimental crosses. BioinformaHcs 19, 889–890 (2003).

40. Warren, W. C. et al. A chromosome-level genome of Astyanax mexicanus surface fish for comparing population-specific genetic differences contributing to trait evolution. Nat Commun 12, 1447 (2021).

41. Bandara, S. et al. The Structural and Biochemical Basis of Apocarotenoid Processing by β- Carotene Oxygenase-2. ACS Chem Biol 16, 480–490 (2021).

42. Misawa, N. et al. Elucidation of the Erwinia uredovora carotenoid biosynthetic pathway by functional analysis of gene products expressed in Escherichia coli. J Bacteriol 172, 6704–12 (1990).

