## Supplemental Figures for "Adaptive Disruption of Carotenoid Metabolism in Cavefish via Loss of Beta-carotene oxygenase 2a (Bco2a)"

#### Supplemental Figure Legends

**Supplemental Figure 1 | Carotenoids in the gut contents of field collected *Astyanax mexicanus***. High-performance liquid chromatography (HPLC) chromatograms displaying the separation of carotenoid standards (A) and those extracted from the gut contents field-collected Río Choy surface fish (B-D) and Pachón cavefish (E-G). Peaks are identified based on retention time and spectral characteristics using standards shown in panel A. The sharp peak at ~2min corresponds to the solvent front (injection peak).

**Supplemental Figure 2 | Carotenoids in the visceral adipose tissue of field collected *Astyanax mexicanus***. High-performance liquid chromatography (HPLC) chromatograms displaying the separation of carotenoid standards (A) and those extracted from the visceral adipose tissue of field-collected Río Choy surface fish (B-D) and Pachón cavefish (E-G). Peaks are identified based on retention time and spectral characteristics using standards shown in panel A. The sharp peak at ~2min corresponds to the solvent front (injection peak).

**Supplemental Figure 3 | Alignment of Río Choy surface fish and Pachón cavefish *bco2a* cDNA.** Two sequences are shown per individual to illustrate heterozygosity. The alignment is presented as the reverse complement of the coding strand. Annotated regions highlight three major mutations found in Pachón cavefish: a variable-length deletion affecting 36 bp of exon 4 and extending into intron 4–5, a 314 bp deletion spanning exon 6 through exon 7, and a nonsense mutation in exon 7 (TAC→TAG).

**Supplemental Figure 4 | Alignment of consensus Bco2a peptide sequences from Río Choy surface fish and Pachón cavefish.** Annotations indicate the short and long Pachón variants (deduced from cDNA) that were codon-optimized and cloned into pMAL-c4x to generate MBP-tagged constructs for expression in carotenoid-producing *E. coli* (see Methods; Supplemental Data 4–6). Catalytic Ferrous iron binding sites are highlighted in green. Variants discovered in Lineage 1 cavefish are highlighted in orange.

### Supplemental Figure 1

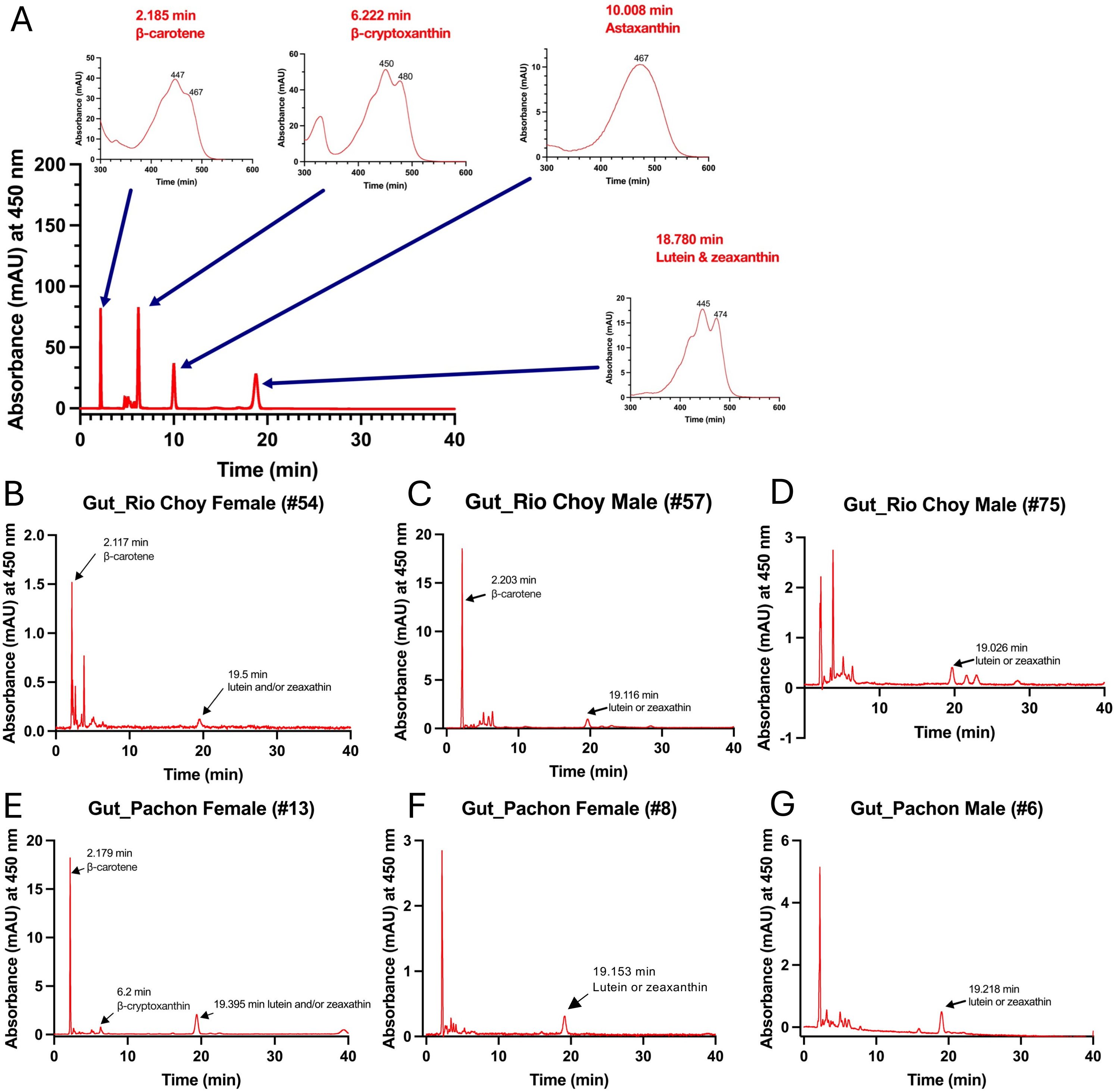

### Supplemental Figure 2

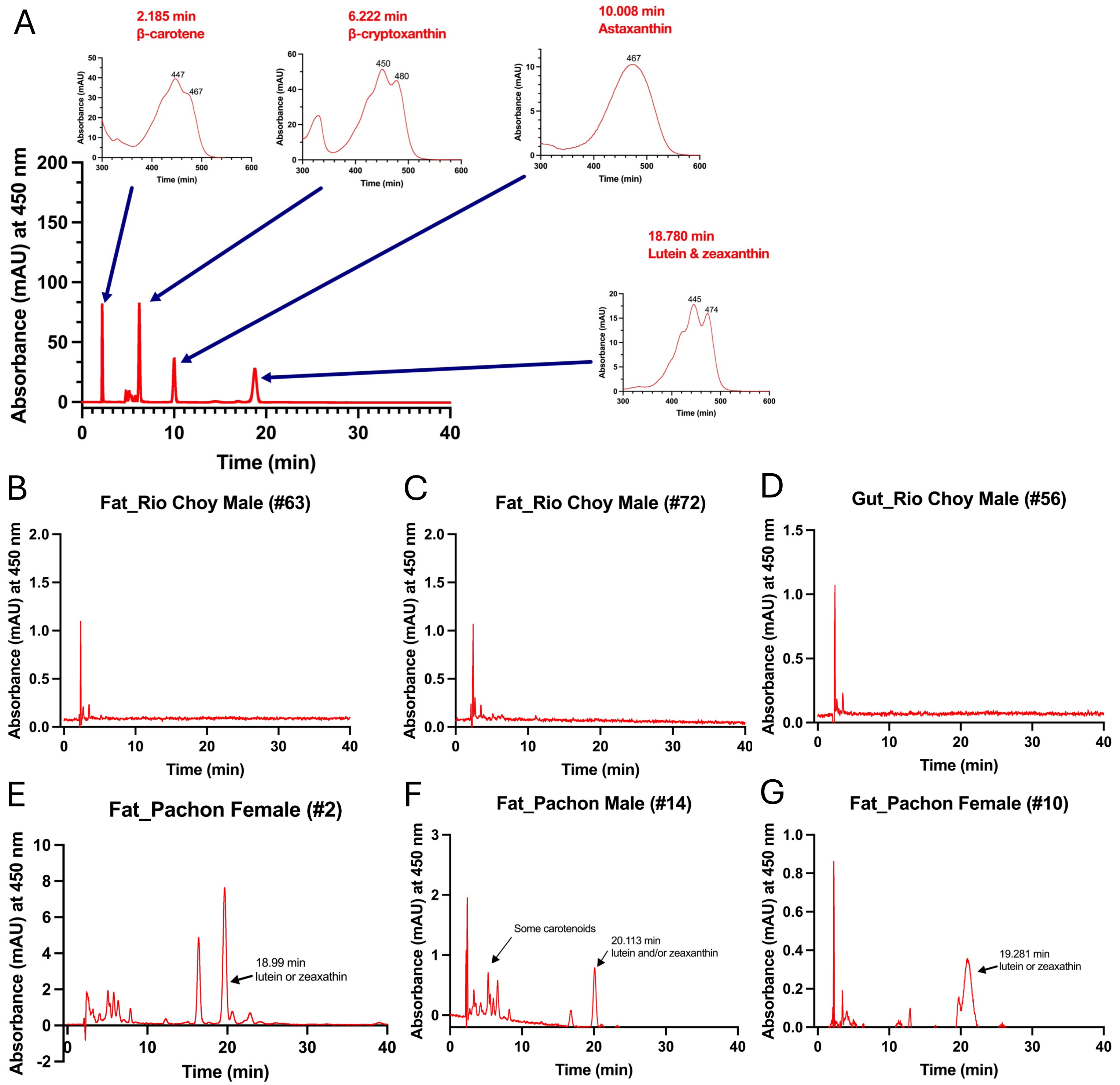

Aligned using MUSCLE

### Supplementary Figure 3

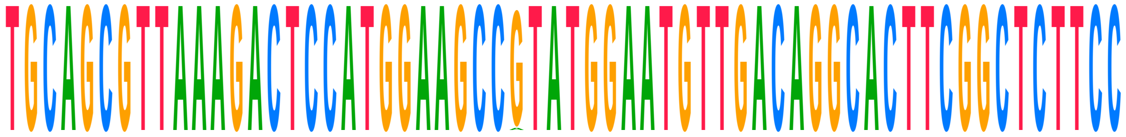

| Sequence | Alignment | Position |
|---|---|---|
| **Consensus** | **TGCAGCGTTAAAGACTCCATGGAAGCCGTATGGAATGTTGACAGGCACTTCGGCTCTTCC** | |
| Choy_1_ref | TGCAGCGTTAAAGACTCCATGGAAGCCGTATGGAATGTTGACAGGCACTTCGGCTCTTCC | 60 |
| Choy_1 | TGCAGCGTTAAAGACTCCATGGAAGCCGTATGGAATGTTGACAGGCACTTCGGCTCTTCC | 60 |
| Choy_10_ref | TGCAGCGTTAAAGACTCCATGGAAGCCGTATGGAATGTTGACAGGCACTTCGGCTCTTCC | 60 |
| Choy_10 | TGCAGCGTTAAAGACTCCATGGAAGCCGTATGGAATGTTGACAGGCACTTCGGCTCTTCC | 60 |
| Choy_11_ref | TGCAGCGTTAAAGACTCCATGGAAGCCGTATGGAATGTTGACAGGCACTTCGGCTCTTCC | 60 |
| Choy_11 | TGCAGCGTTAAAGACTCCATGGAAGCCGTATGGAATGTTGACAGGCACTTCGGCTCTTCC | 60 |
| Choy_12_ref | TGCAGCGTTAAAGACTCCATGGAAGCCGTATGGAATGTTGACAGGCACTTCGGCTCTTCC | 60 |
| Choy_12 | TGCAGCGTTAAAGACTCCATGGAAGCCATATGGAATGTTGACAGGCACTTCGGCTCTTCC | 60 |
| Choy_13_ref | TGCAGCGTTAAAGACTCCATGGAAGCCGTATGGAATGTTGACAGGCACTTCGGCTCTTCC | 60 |
| Choy_13 | TGCAGCGTTAAAGACTCCATGGAAGCCGTATGGAATGTTGACAGGCACTTCGGCTCTTCC | 60 |
| Choy_14_ref | TGCAGCGTTAAAGACTCCATGGAAGCCGTATGGAATGTTGACAGGCACTTCGGCTCTTCC | 60 |
| Choy_14 | TGCAGCGTTAAAGACTCCATGGAAGCCGTATGGAATGTTGACAGGCACTTCGGCTCTTCC | 60 |
| Choy_5_ref | TGCAGCGTTAAAGACTCCATGGAAGCCGTATGGAATGTTGACAGGCACTTCGGCTCTTCC | 60 |
| Choy_5 | TGCAGCGTTAAAGACTCCATGGAAGCCGTATGGAATGTTGACAGGCACTTCGGCTCTTCC | 60 |
| Choy_6_ref | TGCAGCGTTAAAGACTCCATGGAAGCCGTATGGAATGTTGACAGGCACTTCGGCTCTTCC | 60 |
| Choy_6 | TGCAGCGTTAAAGACTCCATGGAAGCCGTATGGAATGTTGACAGGCACTTCGGCTCTTCC | 60 |
| Choy_9_ref | TGCAGCGTTAAAGACTCCATGGAAGCCGTATGGAATGTTGACAGGCACTTCGGCTCTTCC | 60 |
| Choy_9 | TGCAGCGTTAAAGACTCCATGGAAGCCGTATGGAATGTTGACAGGCACTTCGGCTCTTCC | 60 |
| Pachon_wild_bco2a | | |
| Pachon_1Gross | TGCAGCGTTAAAGACTCCATGGAAGCCGTATGGAATGTTGACAGGCACTTCGGCTCTTCC | 60 |
| Pach_11 | TGCAGCGTTAAAGACTCCATGGAAGCCGTATGGAATGTTGACAGGCACTTCGGCTCTTCC | 60 |
| Pachon...ss_ref | TGCAGCGTTAAAGACTCCATGGAAGCCGTATGGAATGTTGACAGGCACTTCGGCTCTTCC | 60 |
| Pachon2B_ref | TGCAGCGTTAAAGACTCCATGGAAGCCGTATGGAATGTTGACAGGCACTTCGGCTCTTCC | 60 |
| Pachon...OG_ref | TGCAGCGTTAAAGACTCCATGGAAGCCGTATGGAATGTTGACAGGCACTTCGGCTCTTCC | 60 |
| Pach_12_ref | TGCAGCGTTAAAGACTCCATGGAAGCCGTATGGAATGTTGACAGGCACTTCGGCTCTTCC | 60 |
| Pach_12 | TGCAGCGTTAAAGACTCCATGGAAGCCGTATGGAATGTTGACAGGCACTTCGGCTCTTCC | 60 |
| Pach_17 | TGCAGCGTTAAAGACTCCATGGAAGCCGTATGGAATGTTGACAGGCACTTCGGCTCTTCC | 60 |
| Pachon_F2POG | TGCAGCGTTAAAGACTCCATGGAAGCCGTATGGAATGTTGACAGGCACTTCGGCTCTTCC | 60 |
| Pachon_3Gross | TGCAGCGTTAAAGACTCCATGGAAGCCGTATGGAATGTTGACAGGCACTTCGGCTCTTCC | 60 |
| Pach_15_ref | TGCAGCGTTAAAGACTCCATGGAAGCCGTATGGAATGTTGACAGGCACTTCGGCTCTTCC | 60 |
| Pachon...ro_ref | TGCAGCGTTAAAGACTCCATGGAAGCCGTATGGAATGTTGACAGGCACTTCGGCTCTTCC | 60 |
| Pachon...ss_ref | TGCAGCGTTAAAGACTCCATGGAAGCCGTATGGAATGTTGACAGGCACTTCGGCTCTTCC | 60 |
| Pachon_2Gross | TGCAGCGTTAAAGACTCCATGGAAGCCGTATGGAATGTTGACAGGCACTTCGGCTCTTCC | 60 |
| Pach_17_ref | TGCAGCGTTAAAGACTCCATGGAAGCCGTATGGAATGTTGACAGGCACTTCGGCTCTTCC | 60 |
| Pachon...ss_ref | TGCAGCGTTAAAGACTCCATGGAAGCCGTATGGAATGTTGACAGGCACTTCGGCTCTTCC | 60 |
| Pachon_ref_ref | TGCAGCGTTAAAGACTCCATGGAAGCCGTATGGAATGTTGACAGGCACTTCGGCTCTTCC | 60 |
| Pach_8 | TGCAGCGTTAAAGACTCCATGGAAGCCGTATGGAATGTTGACAGGCACTTCGGCTCTTCC | 60 |
| Pach_8_ref | TGCAGCGTTAAAGACTCCATGGAAGCCGTATGGAATGTTGACAGGCACTTCGGCTCTTCC | 60 |
| Pachon...OG_ref | TGCAGCGTTAAAGACTCCATGGAAGCCGTATGGAATGTTGACAGGCACTTCGGCTCTTCC | 60 |
| Pachon_6Boro | TGCAGCGTTAAAGACTCCATGGAAGCCGTATGGAATGTTGACAGGCACTTCGGCTCTTCC | 60 |
| Pachon_ref | TGCAGCGTTAAAGACTCCATGGAAGCCGTATGGAATGTTGACAGGCACTTCGGCTCTTCC | 60 |
| Pach_3_ref | TGCAGCGTTAAAGACTCCATGGAAGCCGTATGGAATGTTGACAGGCACTTCGGCTCTTCC | 60 |
| Pach_9 | TGCAGCGTTAAAGACTCCATGGAAGCCGTATGGAATGTTGACAGGCACTTCGGCTCTTCC | 60 |
| Pachon_G2POG | TGCAGCGTTAAAGACTCCATGGAAGCCGTATGGAATGTTGACAGGCACTTCGGCTCTTCC | 60 |
| Pachon...OG_ref | TGCAGCGTTAAAGACTCCATGGAAGCCGTATGGAATGTTGACAGGCACTTCGGCTCTTCC | 60 |
| Pach_9_ref | TGCAGCGTTAAAGACTCCATGGAAGCCGTATGGAATGTTGACAGGCACTTCGGCTCTTCC | 60 |
| Pachon2B | TGCAGCGTTAAAGACTCCATGGAAGCCGTATGGAATGTTGACAGGCACTTCGGCTCTTCC | 60 |
| Pachon...OG_ref | TGCAGCGTTAAAGACTCCATGGAAGCCGTATGGAATGTTGACAGGCACTTCGGCTCTTCC | 60 |
| Pachon_E2POG | TGCAGCGTTAAAGACTCCATGGAAGCCGTATGGAATGTTGACAGGCACTTCGGCTCTTCC | 60 |
| Pach_14 | TGCAGCGTTAAAGACTCCATGGAAGCCGTATGGAATGTTGACAGGCACTTCGGCTCTTCC | 60 |
| Pach_3 | TGCAGCGTTAAAGACTCCATGGAAGCCGTATGGAATGTTGACAGGCACTTCGGCTCTTCC | 60 |
| Pachon_H2POG | TGCAGCGTTAAAGACTCCATGGAAGCCGTATGGAATGTTGACAGGCACTTCGGCTCTTCC | 60 |
| Pach_7_ref | TGCAGCGTTAAAGACTCCATGGAAGCCGTATGGAATGTTGACAGGCACTTCGGCTCTTCC | 60 |
| Pach_15 | TGCAGCGTTAAAGACTCCATGGAAGCCGTATGGAATGTTGACAGGCACTTCGGCTCTTCC | 60 |
| Pach_14_ref | TGCAGCGTTAAAGACTCCATGGAAGCCGTATGGAATGTTGACAGGCACTTCGGCTCTTCC | 60 |

| | | |
|---|---|---|
| Pach_11_ref | TGCAGCGTTAAAGACTCCATGGAAGCCGTATGGAATGTTGACAGGCACTTCGGCTCTTCC | 60 |
| Pach_7 | TGCAGCGTTAAAGACTCCATGGAAGCCGTATGGAATGTTGACAGGCACTTCGGCTCTTCC | 60 |

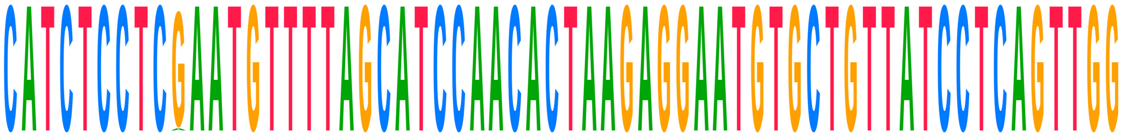

| | | |
|---|---|---|
| **Consensus** | **CATCTCCTCGAATGTTTTAGCATCCAACACTAAGAGGAATGTGCTGTTATCCTCAGTTGG** | |
| Choy_1_ref | CATCTCCTCGAATGTTTTAGCATCCAACACTAAGAGGAATGTGCTGTTATCCTCAGTTGG | 120 |
| Choy_1 | CATCTCCTCAAATGTTTTAGCATCCAACACTAAGAGGAATGTGCTGTTATCCTCAGTTGG | 120 |
| Choy_10_ref | CATCTCCTCGAATGTTTTAGCATCCAACACTAAGAGGAATGTGCTGTTATCCTCAGTTGG | 120 |
| Choy_10 | CATCTCCTCGAATGTTTTAGCATCCAACACTAAGAGGAATGTGCTGTTATCCTCAGTTGG | 120 |
| Choy_11_ref | CATCTCCTCGAATGTTTTAGCATCCAACACTAAGAGGAATGTGCTGTTATCCTCAGTTGG | 120 |
| Choy_11 | CATCTCCTCGAATGTTTTAGCATCCAACACTAAGAGGAATGTGCTGTTATCCTCAGTTGG | 120 |
| Choy_12_ref | CATCTCCTCGAATGTTTTAGCATCCAACACTAAGAGGAATGTGCTGTTATCCTCAGTTGG | 120 |
| Choy_12 | CATCTCCTCGAATGTTTTAGCATCCAACACTAAGAGGAATGTGCTGTTATCCTCAGTTGG | 120 |
| Choy_13_ref | CATCTCCTCGAATGTTTTAGCATCCAACACTAAGAGGAATGTGCTGTTATCCTCAGTTGG | 120 |
| Choy_13 | CATCTCCTCGAATGTTTTAGCATCCAACACTAAGAGGAATGTGCTGTTATCCTCAGTTGG | 120 |
| Choy_14_ref | CATCTCCTCGAATGTTTTAGCATCCAACACTAAGAGGAATGTGCTGTTATCCTCAGTTGG | 120 |
| Choy_14 | CATCTCCTCGAATGTTTTAGCATCCAACACTAAGAGGAATGTGCTGTTATCCTCAGTTGG | 120 |
| Choy_5_ref | CATCTCCTCGAATGTTTTAGCATCCAACACTAAGAGGAATGTGCTGTTATCCTCAGTTGG | 120 |
| Choy_5 | CATCTCCTCGAATGTTTTAGCATCCAACACTAAGAGGAATGTGCTGTTATCCTCAGTTGG | 120 |
| Choy_6_ref | CATCTCCTCGAATGTTTTAGCATCCAACACTAAGAGGAATGTGCTGTTATCCTCAGTTGG | 120 |
| Choy_6 | CATCTCCTCGAATGTTTTAGCATCCAACACTAAGAGGAATGTGCTGTTATCCTCAGTTGG | 120 |
| Choy_9_ref | CATCTCCTCGAATGTTTTAGCATCCAACACTAAGAGGAATGTGCTGTTATCCTCAGTTGG | 120 |
| Choy_9 | CATCTCCTCGAATGTTTTAGCATCCAACACTAAGAGGAATGTGCTGTTATCCTCAGTTGG | 120 |
| Pachon_wild_bco2a | | |
| Pachon_1Gross | CATCTCCTCGAATGTTTTAGCATCCAACACTAAGAGGAATGTGCTGTTATCCTCAGTTGG | 120 |
| Pach_11 | CATCTCCTCGAATGTTTTAGCATCCAACACTAAGAGGAATGTGCTGTTATCCTCAGTTGG | 120 |
| Pachon...ss_ref | CATCTCCTCGAATGTTTTAGCATCCAACACTAAGAGGAATGTGCTGTTATCCTCAGTTGG | 120 |
| Pachon2B_ref | CATCTCCTCGAATGTTTTAGCATCCAACACTAAGAGGAATGTGCTGTTATCCTCAGTTGG | 120 |
| Pachon...OG_ref | CATCTCCTCGAATGTTTTAGCATCCAACACTAAGAGGAATGTGCTGTTATCCTCAGTTGG | 120 |
| Pach_12_ref | CATCTCCTCGAATGTTTTAGCATCCAACACTAAGAGGAATGTGCTGTTATCCTCAGTTGG | 120 |
| Pach_12 | CATCTCCTCGAATGTTTTAGCATCCAACACTAAGAGGAATGTGCTGTTATCCTCAGTTGG | 120 |
| Pach_17 | CATCTCCTCGAATGTTTTAGCATCCAACACTAAGAGGAATGTGCTGTTATCCTCAGTTGG | 120 |
| Pachon_F2POG | CATCTCCTCGAATGTTTTAGCATCCAACACTAAGAGGAATGTGCTGTTATCCTCAGTTGG | 120 |
| Pachon_3Gross | CATCTCCTCGAATGTTTTAGCATCCAACACTAAGAGGAATGTGCTGTTATCCTCAGTTGG | 120 |
| Pach_15_ref | CATCTCCTCGAATGTTTTAGCATCCAACACTAAGAGGAATGTGCTGTTATCCTCAGTTGG | 120 |
| Pachon...ro_ref | CATCTCCTCGAATGTTTTAGCATCCAACACTAAGAGGAATGTGCTGTTATCCTCAGTTGG | 120 |
| Pachon...ss_ref | CATCTCCTCGAATGTTTTAGCATCCAACACTAAGAGGAATGTGCTGTTATCCTCAGTTGG | 120 |
| Pachon_2Gross | CATCTCCTCGAATGTTTTAGCATCCAACACTAAGAGGAATGTGCTGTTATCCTCAGTTGG | 120 |
| Pach_17_ref | CATCTCCTCGAATGTTTTAGCATCCAACACTAAGAGGAATGTGCTGTTATCCTCAGTTGG | 120 |
| Pachon...ss_ref | CATCTCCTCGAATGTTTTAGCATCCAACACTAAGAGGAATGTGCTGTTATCCTCAGTTGG | 120 |
| Pachon_ref_ref | CATCTCCTCGAATGTTTTAGCATCCAACACTAAGAGGAATGTGCTGTTATCCTCAGTTGG | 120 |
| Pach_8 | CATCTCCTCGAATGTTTTAGCATCCAACACTAAGAGGAATGTGCTGTTATCCTCAGTTGG | 120 |
| Pach_8_ref | CATCTCCTCGAATGTTTTAGCATCCAACACTAAGAGGAATGTGCTGTTATCCTCAGTTGG | 120 |
| Pachon...OG_ref | CATCTCCTCGAATGTTTTAGCATCCAACACTAAGAGGAATGTGCTGTTATCCTCAGTTGG | 120 |
| Pachon_6Boro | CATCTCCTCGAATGTTTTAGCATCCAACACTAAGAGGAATGTGCTGTTATCCTCAGTTGG | 120 |
| Pachon_ref | CATCTCCTCGAATGTTTTAGCATCCAACACTAAGAGGAATGTGCTGTTATCCTCAGTTGG | 120 |
| Pach_3_ref | CATCTCCTCGAATGTTTTAGCATCCAACACTAAGAGGAATGTGCTGTTATCCTCAGTTGG | 120 |
| Pach_9 | CATCTCCTCGAATGTTTTAGCATCCAACACTAAGAGGAATGTGCTGTTATCCTCAGTTGG | 120 |
| Pachon_G2POG | CATCTCCTCGAATGTTTTAGCATCCAACACTAAGAGGAATGTGCTGTTATCCTCAGTTGG | 120 |
| Pachon...OG_ref | CATCTCCTCGAATGTTTTAGCATCCAACACTAAGAGGAATGTGCTGTTATCCTCAGTTGG | 120 |
| Pach_9_ref | CATCTCCTCGAATGTTTTAGCATCCAACACTAAGAGGAATGTGCTGTTATCCTCAGTTGG | 120 |
| Pachon2B | CATCTCCTCGAATGTTTTAGCATCCAACACTAAGAGGAATGTGCTGTTATCCTCAGTTGG | 120 |
| Pachon...OG_ref | CATCTCCTCGAATGTTTTAGCATCCAACACTAAGAGGAATGTGCTGTTATCCTCAGTTGG | 120 |
| Pachon_E2POG | CATCTCCTCGAATGTTTTAGCATCCAACACTAAGAGGAATGTGCTGTTATCCTCAGTTGG | 120 |
| Pach_14 | CATCTCCTCGAATGTTTTAGCATCCAACACTAAGAGGAATGTGCTGTTATCCTCAGTTGG | 120 |
| Pach_3 | CATCTCCTCGAATGTTTTAGCATCCAACACTAAGAGGAATGTGCTGTTATCCTCAGTTGG | 120 |
| Pachon_H2POG | CATCTCCTCGAATGTTTTAGCATCCAACACTAAGAGGAATGTGCTGTTATCCTCAGTTGG | 120 |

| | | |
|---|---|---|
| Pach_7_ref | CATCTCCTCGAATGTTTTAGCATCCAACACTAAGAGGAATGTGCTGTTATCCTCAGTTGG | 120 |
| Pach_15 | CATCTCCTCGAATGTTTTAGCATCCAACACTAAGAGGAATGTGCTGTTATCCTCAGTTGG | 120 |
| Pach_14_ref | CATCTCCTCGAATGTTTTAGCATCCAACACTAAGAGGAATGTGCTGTTATCCTCAGTTGG | 120 |
| Pach_11_ref | CATCTCCTCGAATGTTTTAGCATCCAACACTAAGAGGAATGTGCTGTTATCCTCAGTTGG | 120 |
| Pach_7 | CATCTCCTCGAATGTTTTAGCATCCAACACTAAGAGGAATGTGCTGTTATCCTCAGTTGG | 120 |

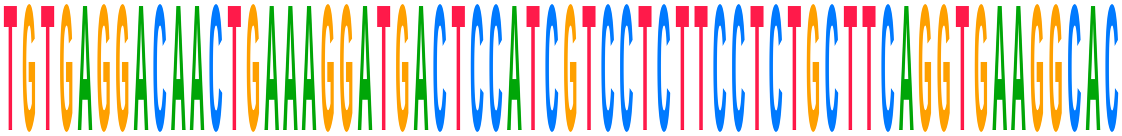


| | | |
|---|---|---|
| **Consensus** | **TGTGAGGACAACTGAAAGGATGACTCCATCGTCCTCTTCCTCTGCTTCAGGTGAAGGCAC** | |
| Choy_1_ref | TGTGAGGACAACTGAAAGGATGACTCCATCGTCCTCTTCCTCTGCTTCAGGTGAAGGCAC | 180 |
| Choy_1 | TGTGAGGACAACTGAAAGGATGACTCCATCGTCCTCTTCCTCTGCTTCAGGTGAAGGCAC | 180 |
| Choy_10_ref | TGTGAGGACAACTGAAAGGATGACTCCATCGTCCTCTTCCTCTGCTTCAGGTGAAGGCAC | 180 |
| Choy_10 | TGTGAGGACAACTGAAAGGATGACTCCATCGTCCTCTTCCTCTGCTTCAGGTGAAGGCAC | 180 |
| Choy_11_ref | TGTGAGGACAACTGAAAGGATGACTCCATCGTCCTCTTCCTCTGCTTCAGGTGAAGGCAC | 180 |
| Choy_11 | TGTGAGGACAACTGAAAGGATGACTCCATCGTCCTCTTCCTCTGCTTCAGGTGAAGGCAC | 180 |
| Choy_12_ref | TGTGAGGACAACTGAAAGGATGACTCCATCGTCCTCTTCCTCTGCTTCAGGTGAAGGCAC | 180 |
| Choy_12 | TGTGAGGACAACTGAAAGGATGACTCCATCGTCCTCTTCCTCTGCTTCAGGTGAAGGCAC | 180 |
| Choy_13_ref | TGTGAGGACAACTGAAAGGATGACTCCATCGTCCTCTTCCTCTGCTTCAGGTGAAGGCAC | 180 |
| Choy_13 | TGTGAGGACAACTGAAAGGATGACTCCATCGTCCTCTTCCTCTGCTTCAGGTGAAGGCAC | 180 |
| Choy_14_ref | TGTGAGGACAACTGAAAGGATGACTCCATCGTCCTCTTCCTCTGCTTCAGGTGAAGGCAC | 180 |
| Choy_14 | TGTGAGGACAACTGAAAGGATGACTCCATCGTCCTCTTCCTCTGCTTCAGGTGAAGGCAC | 180 |
| Choy_5_ref | TGTGAGGACAACTGAAAGGATGACTCCATCGTCCTCTTCCTCTGCTTCAGGTGAAGGCAC | 180 |
| Choy_5 | TGTGAGGACAACTGAAAGGATGACTCCATCGTCCTCTTCCTCTGCTTCAGGTGAAGGCAC | 180 |
| Choy_6_ref | TGTGAGGACAACTGAAAGGATGACTCCATCGTCCTCTTCCTCTGCTTCAGGTGAAGGCAC | 180 |
| Choy_6 | TGTGAGGACAACTGAAAGGATGACTCCATCGTCCTCTTCCTCTGCTTCAGGTGAAGGCAC | 180 |
| Choy_9_ref | TGTGAGGACAACTGAAAGGATGACTCCATCGTCCTCTTCCTCTGCTTCAGGTGAAGGCAC | 180 |
| Choy_9 | TGTGAGGACAACTGAAAGGATGACTCCATCGTCCTCTTCCTCTGCTTCAGGTGAAGGCAC | 180 |
| Pachon_wild_bco2a | | |
| Pachon_1Gross | TGTGAGGACAACTGAAAGGATGACTCCATCGTCCTCTTCCTCTGCTTCAGGTGAAGGCAC | 180 |
| Pach_11 | TGTGAGGACAACTGAAAGGATGACTCCATCGTCCTCTTCCTCTGCTTCAGGTGAAGGCAC | 180 |
| Pachon…ss_ref | TGTGAGGACAACTGAAAGGATGACTCCATCGTCCTCTTCCTCTGCTTCAGGTGAAGGCAC | 180 |
| Pachon2B_ref | TGTGAGGACAACTGAAAGGATGACTCCATCGTCCTCTTCCTCTGCTTCAGGTGAAGGCAC | 180 |
| Pachon…OG_ref | TGTGAGGACAACTGAAAGGATGACTCCATCGTCCTCTTCCTCTGCTTCAGGTGAAGGCAC | 180 |
| Pach_12_ref | TGTGAGGACAACTGAAAGGATGACTCCATCGTCCTCTTCCTCTGCTTCAGGTGAAGGCAC | 180 |
| Pach_12 | TGTGAGGACAACTGAAAGGATGACTCCATCGTCCTCTTCCTCTGCTTCAGGTGAAGGCAC | 180 |
| Pach_17 | TGTGAGGACAACTGAAAGGATGACTCCATCGTCCTCTTCCTCTGCTTCAGGTGAAGGCAC | 180 |
| Pachon_F2POG | TGTGAGGACAACTGAAAGGATGACTCCATCGTCCTCTTCCTCTGCTTCAGGTGAAGGCAC | 180 |
| Pachon_3Gross | TGTGAGGACAACTGAAAGGATGACTCCATCGTCCTCTTCCTCTGCTTCAGGTGAAGGCAC | 180 |
| Pach_15_ref | TGTGAGGACAACTGAAAGGATGACTCCATCGTCCTCTTCCTCTGCTTCAGGTGAAGGCAC | 180 |
| Pachon…ro_ref | TGTGAGGACAACTGAAAGGATGACTCCATCGTCCTCTTCCTCTGCTTCAGGTGAAGGCAC | 180 |
| Pachon…ss_ref | TGTGAGGACAACTGAAAGGATGACTCCATCGTCCTCTTCCTCTGCTTCAGGTGAAGGCAC | 180 |
| Pachon_2Gross | TGTGAGGACAACTGAAAGGATGACTCCATCGTCCTCTTCCTCTGCTTCAGGTGAAGGCAC | 180 |
| Pach_17_ref | TGTGAGGACAACTGAAAGGATGACTCCATCGTCCTCTTCCTCTGCTTCAGGTGAAGGCAC | 180 |
| Pachon…ss_ref | TGTGAGGACAACTGAAAGGATGACTCCATCGTCCTCTTCCTCTGCTTCAGGTGAAGGCAC | 180 |
| Pachon_ref_ref | TGTGAGGACAACTGAAAGGATGACTCCATCGTCCTCTTCCTCTGCTTCAGGTGAAGGCAC | 180 |
| Pach_8 | TGTGAGGACAACTGAAAGGATGACTCCATCGTCCTCTTCCTCTGCTTCAGGTGAAGGCAC | 180 |
| Pach_8_ref | TGTGAGGACAACTGAAAGGATGACTCCATCGTCCTCTTCCTCTGCTTCAGGTGAAGGCAC | 180 |
| Pachon…OG_ref | TGTGAGGACAACTGAAAGGATGACTCCATCGTCCTCTTCCTCTGCTTCAGGTGAAGGCAC | 180 |
| Pachon_6Boro | TGTGAGGACAACTGAAAGGATGACTCCATCGTCCTCTTCCTCTGCTTCAGGTGAAGGCAC | 180 |
| Pachon_ref | TGTGAGGACAACTGAAAGGATGACTCCATCGTCCTCTTCCTCTGCTTCAGGTGAAGGCAC | 180 |
| Pach_3_ref | TGTGAGGACAACTGAAAGGATGACTCCATCGTCCTCTTCCTCTGCTTCAGGTGAAGGCAC | 180 |
| Pach_9 | TGTGAGGACAACTGAAAGGATGACTCCATCGTCCTCTTCCTCTGCTTCAGGTGAAGGCAC | 180 |
| Pachon_G2POG | TGTGAGGACAACTGAAAGGATGACTCCATCGTCCTCTTCCTCTGCTTCAGGTGAAGGCAC | 180 |
| Pachon…OG_ref | TGTGAGGACAACTGAAAGGATGACTCCATCGTCCTCTTCCTCTGCTTCAGGTGAAGGCAC | 180 |
| Pach_9_ref | TGTGAGGACAACTGAAAGGATGACTCCATCGTCCTCTTCCTCTGCTTCAGGTGAAGGCAC | 180 |
| Pachon2B | TGTGAGGACAACTGAAAGGATGACTCCATCGTCCTCTTCCTCTGCTTCAGGTGAAGGCAC | 180 |
| Pachon…OG_ref | TGTGAGGACAACTGAAAGGATGACTCCATCGTCCTCTTCCTCTGCTTCAGGTGAAGGCAC | 180 |
| Pachon_E2POG | TGTGAGGACAACTGAAAGGATGACTCCATCGTCCTCTTCCTCTGCTTCAGGTGAAGGCAC | 180 |

| | | |
|---|---|---|
| Pach_14 | TGTGAGGACAACTGAAAGGATGACTCCATCGTCCTCTTCCTCTGCTTCAGGTGAAGGCAC | 180 |
| Pach_3 | TGTGAGGACAACTGAAAGGATGACTCCATCGTCCTCTTCCTCTGCTTCAGGTGAAGGCAC | 180 |
| Pachon_H2POG | TGTGAGGACAACTGAAAGGATGACTCCATCGTCCTCTTCCTCTGCTTCAGGTGAAGGCAC | 180 |
| Pach_7_ref | TGTGAGGACAACTGAAAGGATGACTCCATCGTCCTCTTCCTCTGCTTCAGGTGAAGGCAC | 180 |
| Pach_15 | TGTGAGGACAACTGAAAGGATGACTCCATCGTCCTCTTCCTCTGCTTCAGGTGAAGGCAC | 180 |
| Pach_14_ref | TGTGAGGACAACTGAAAGGATGACTCCATCGTCCTCTTCCTCTGCTTCAGGTGAAGGCAC | 180 |
| Pach_11_ref | TGTGAGGACAACTGAAAGGATGACTCCATCGTCCTCTTCCTCTGCTTCAGGTGAAGGCAC | 180 |
| Pach_7 | TGTGAGGACAACTGAAAGGATGACTCCATCGTCCTCTTCCTCTGCTTCAGGTGAAGGCAC | 180 |

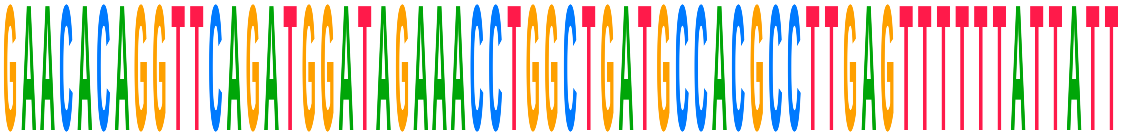

| | | |
|---|---|---|
| **Consensus** | **GAACACAGGTTCAGATGGATAGAAACCTGGCTGATGCCACGCCTTGAGTTTTTTATTATT** | |
| Choy_1_ref | GAACACAGGTTCAGATGGATAGAAACCTGGCTGATGCCACGCCTTGAGTTTTTTATTATT | 240 |
| Choy_1 | GAACACAGGTTCAGATGGATAGAAACCTGGCTGATGCCACGCCTTGAGTTTTTTATTATT | 240 |
| Choy_10_ref | GAACACAGGTTCAGATGGATAGAAACCTGGCTGATGCCACGCCTTGAGTTTTTTATTATT | 240 |
| Choy_10 | GAACACAGGTTCAGATGGATAGAAACCTGGCTGATGCCACGCCTTGAGTTTTTTATTATT | 240 |
| Choy_11_ref | GAACACAGGTTCAGATGGATAGAAACCTGGCTGATGCCACGCCTTGAGTTTTTTATTATT | 240 |
| Choy_11 | GAACACAGGTTCAGATGGATAGAAACCTGGCTGATGCCACGCCTTGAGTTTTTTATTATT | 240 |
| Choy_12_ref | GAACACAGGTTCAGATGGATAGAAACCTGGCTGATGCCACGCCTTGAGTTTTTTATTATT | 240 |
| Choy_12 | GAACACAGGTTCAGATGGATAGAAACCTGGCTGATGCCACGCCTTGAGTTTTTTATTATT | 240 |
| Choy_13_ref | GAACACAGGTTCAGATGGATAGAAACCTGGCTGATGCCACGCCTTGAGTTTTTTATTATT | 240 |
| Choy_13 | GAACACAGGTTCAGATGGATAGAAACCTGGCTGATGCCACGCCTTGAGTTTTTTATTATT | 240 |
| Choy_14_ref | GAACACAGGTTCAGATGGATAGAAACCTGGCTGATGCCACGCCTTGAGTTTTTTATTATT | 240 |
| Choy_14 | GAACACAGGTTCAGATGGATAGAAACCTGGCTGATGCCACGCCTTGAGTTTTTTATTATT | 240 |
| Choy_5_ref | GAACACAGGTTCAGATGGATAGAAACCTGGCTGATGCCACGCCTTGAGTTTTTTATTATT | 240 |
| Choy_5 | GAACACAGGTTCAGATGGATAGAAACCTGGCTGATGCCACGCCTTGAGTTTTTTATTATT | 240 |
| Choy_6_ref | GAACACAGGTTCAGATGGATAGAAACCTGGCTGATGCCACGCCTTGAGTTTTTTATTATT | 240 |
| Choy_6 | GAACACAGGTTCAGATGGATAGAAACCTGGCTGATGCCACGCCTTGAGTTTTTTATTATT | 240 |
| Choy_9_ref | GAACACAGGTTCAGATGGATAGAAACCTGGCTGATGCCACGCCTTGAGTTTTTTATTATT | 240 |
| Choy_9 | GAACACAGGTTCAGATGGATAGAAACCTGGCTGATGCCACGCCTTGAGTTTTTTATTATT | 240 |
| Pachon_wild_bco2a | | |
| Pachon_1Gross | GAACACAGGTTCAGATGGATAGAAACCTGGCTGATGCCACGCCTTGAGTTTTTTATTATT | 240 |
| Pach_11 | GAACACAGGTTCAGATGGATAGAAACCTGGCTGATGCCACGCCTTGAGTTTTTTATTATT | 240 |
| Pachon...ss_ref | GAACACAGGTTCAGATGGATAGAAACCTGGCTGATGCCACGCCTTGAGTTTTTTATTATT | 240 |
| Pachon2B_ref | GAACACAGGTTCAGATGGATAGAAACCTGGCTGATGCCACGCCTTGAGTTTTTTATTATT | 240 |
| Pachon...OG_ref | GAACACAGGTTCAGATGGATAGAAACCTGGCTGATGCCACGCCTTGAGTTTTTTATTATT | 240 |
| Pach_12_ref | GAACACAGGTTCAGATGGATAGAAACCTGGCTGATGCCACGCCTTGAGTTTTTTATTATT | 240 |
| Pach_12 | GAACACAGGTTCAGATGGATAGAAACCTGGCTGATGCCACGCCTTGAGTTTTTTATTATT | 240 |
| Pach_17 | GAACACAGGTTCAGATGGATAGAAACCTGGCTGATGCCACGCCTTGAGTTTTTTATTATT | 240 |
| Pachon_F2POG | GAACACAGGTTCAGATGGATAGAAACCTGGCTGATGCCACGCCTTGAGTTTTTTATTATT | 240 |
| Pachon_3Gross | GAACACAGGTTCAGATGGATAGAAACCTGGCTGATGCCACGCCTTGAGTTTTTTATTATT | 240 |
| Pach_15_ref | GAACACAGGTTCAGATGGATAGAAACCTGGCTGATGCCACGCCTTGAGTTTTTTATTATT | 240 |
| Pachon...ro_ref | GAACACAGGTTCAGATGGATAGAAACCTGGCTGATGCCACGCCTTGAGTTTTTTATTATT | 240 |
| Pachon...ss_ref | GAACACAGGTTCAGATGGATAGAAACCTGGCTGATGCCACGCCTTGAGTTTTTTATTATT | 240 |
| Pachon_2Gross | GAACACAGGTTCAGATGGATAGAAACCTGGCTGATGCCACGCCTTGAGTTTTTTATTATT | 240 |
| Pach_17_ref | GAACACAGGTTCAGATGGATAGAAACCTGGCTGATGCCACGCCTTGAGTTTTTTATTATT | 240 |
| Pachon...ss_ref | GAACACAGGTTCAGATGGATAGAAACCTGGCTGATGCCACGCCTTGAGTTTTTTATTATT | 240 |
| Pachon_ref_ref | GAACACAGGTTCAGATGGATAGAAACCTGGCTGATGCCACGCCTTGAGTTTTTTATTATT | 240 |
| Pach_8 | GAACACAGGTTCAGATGGATAGAAACCTGGCTGATGCCACGCCTTGAGTTTTTTATTATT | 240 |
| Pach_8_ref | GAACACAGGTTCAGATGGATAGAAACCTGGCTGATGCCACGCCTTGAGTTTTTTATTATT | 240 |
| Pachon...OG_ref | GAACACAGGTTCAGATGGATAGAAACCTGGCTGATGCCACGCCTTGAGTTTTTTATTATT | 240 |
| Pachon_6Boro | GAACACAGGTTCAGATGGATAGAAACCTGGCTGATGCCACGCCTTGAGTTTTTTATTATT | 240 |
| Pachon_ref | GAACACAGGTTCAGATGGATAGAAACCTGGCTGATGCCACGCCTTGAGTTTTTTATTATT | 240 |
| Pach_3_ref | GAACACAGGTTCAGATGGATAGAAACCTGGCTGATGCCACGCCTTGAGTTTTTTATTATT | 240 |
| Pach_9 | GAACACAGGTTCAGATGGATAGAAACCTGGCTGATGCCACGCCTTGAGTTTTTTATTATT | 240 |
| Pachon_G2POG | GAACACAGGTTCAGATGGATAGAAACCTGGCTGATGCCACGCCTTGAGTTTTTTATTATT | 240 |
| Pachon...OG_ref | GAACACAGGTTCAGATGGATAGAAACCTGGCTGATGCCACGCCTTGAGTTTTTTATTATT | 240 |
| Pach_9_ref | GAACACAGGTTCAGATGGATAGAAACCTGGCTGATGCCACGCCTTGAGTTTTTTATTATT | 240 |

| | | |
|---|---|---|
| Pachon2B | GAACACAGGTTCAGATGGATAGAAACCTGGCTGATGCCACGCCTTGAGTTTTTTATTATT | 240 |
| Pachon...OG_ref | GAACACAGGTTCAGATGGATAGAAACCTGGCTGATGCCACGCCTTGAGTTTTTTATTATT | 240 |
| Pachon_E2POG | GAACACAGGTTCAGATGGATAGAAACCTGGCTGATGCCACGCCTTGAGTTTTTTATTATT | 240 |
| Pach_14 | GAACACAGGTTCAGATGGATAGAAACCTGGCTGATGCCACGCCTTGAGTTTTTTATTATT | 240 |
| Pach_3 | GAACACAGGTTCAGATGGATAGAAACCTGGCTGATGCCACGCCTTGAGTTTTTTATTATT | 240 |
| Pachon_H2POG | GAACACAGGTTCAGATGGATAGAAACCTGGCTGATGCCACGCCTTGAGTTTTTTATTATT | 240 |
| Pach_7_ref | GAACACAGGTTCAGATGGATAGAAACCTGGCTGATGCCACGCCTTGAGTTTTTTATTATT | 240 |
| Pach_15 | GAACACAGGTTCAGATGGATAGAAACCTGGCTGATGCCACGCCTTGAGTTTTTTATTATT | 240 |
| Pach_14_ref | GAACACAGGTTCAGATGGATAGAAACCTGGCTGATGCCACGCCTTGAGTTTTTTATTATT | 240 |
| Pach_11_ref | GAACACAGGTTCAGATGGATAGAAACCTGGCTGATGCCACGCCTTGAGTTTTTTATTATT | 240 |
| Pach_7 | GAACACAGGTTCAGATGGATAGAAACCTGGCTGATGCCACGCCTTGAGTTTTTTATTATT | 240 |

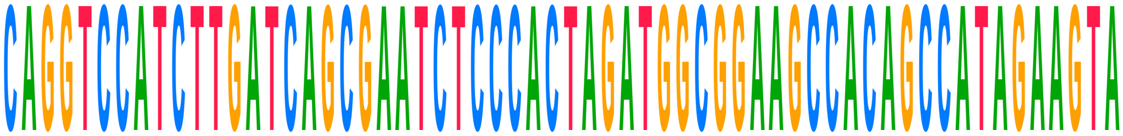

| | | |
|---|---|---|
| **Consensus** | **CAGGTCCATCTTGATCAGCGAATCTCCCACTAGATGGCGGAAGCCACAGCCATAGAAGTA** | |
| Choy_1_ref | CAGGTCCATCTTGATCAGCGAATCTCCCACTAGATGGCGGAAGCCACAGCCATAGAAGTA | 300 |
| Choy_1 | CAGGTCCATCTTGATCAGCGAATCTCCCACTAGATGGCGGAAGCCACAGCCATAGAAGTA | 300 |
| Choy_10_ref | CAGGTCCATCTTGATCAGCGAATCTCCCACTAGATGGCGGAAGCCACAGCCATAGAAGTA | 300 |
| Choy_10 | CAGGTCCATCTTGATCAGCGAATCTCCCACTAGATGGCGGAAGCCACAGCCATAGAAGTA | 300 |
| Choy_11_ref | CAGGTCCATCTTG------------------AGATGGCGGAAGCCACAGCCATAGAAGTA | 282 |
| Choy_11 | CAGGTCCATCTTG------------------AGATGGCGGAAGCCACAGCCATAGAAGTA | 282 |
| Choy_12_ref | CAGGTCCATCTTGATCAGCGAATCTCCCACTAGATGGCGGAAGCCACAGCCATAGAAGTA | 300 |
| Choy_12 | CAGGTCCATCTTGATCAGCGAATCTCCCACTAGATGGCGGAAGCCACAGCCATAGAAGTA | 300 |
| Choy_13_ref | CAGGTCCATCTTGATCAGCGAATCTCCCACTAGATGGCGGAAGCCACAGCCATAGAAGTA | 300 |
| Choy_13 | CAGGTCCATCTTGATCAGCGAATCTCCCACTAGATGGCGGAAGCCACAGCCATAGAAGTA | 300 |
| Choy_14_ref | CAGGTCCATCTTGATCAGCGAATCTCCCACTAGATGGCGGAAGCCACAGCCATAGAAGTA | 300 |
| Choy_14 | CAGGTCCATCTTGATCAGCGAATCTCCCACTAGATGGCGGAAGCCACAGCCATAGAAGTA | 300 |
| Choy_5_ref | CAGGTCCATCTTGATCAGCGAATCTCCCACTAGATGGCGGAAGCCACAGCCATAGAAGTA | 300 |
| Choy_5 | CAGGTCCATCTTGATCAGCGAATCTCCCACTAGATGGCGGAAGCCACAGCCATAGAAGTA | 300 |
| Choy_6_ref | CAGGTCCATCTTGATCAGCGAATCTCCCACTAGATGGCGGAAGCCACAGCCATAGAAGTA | 300 |
| Choy_6 | CAGGTCCATCTTGATCAGCGAATCTCCCACTAGATGGCGGAAGCCACAGCCATAGAAGTA | 300 |
| Choy_9_ref | CAGGTCCATCTTGATCAGCGAATCTCCCACTAGATGGCGGAAGCCACAGCCATAGAAGTA | 300 |
| Choy_9 | CAGGTCCATCTTGATCAGCGAATCTCCCACTAGATGGCGGAAGCCACAGCCATAGAAGTA | 300 |
| Pachon_wild_bco2a | | |
| Pachon_1Gross | CAGGTCCATCTTGATCAGCGAATCTCCCACTAGATGGCGGAAGCCACAGCCATAGAAGTA | 300 |
| Pach_11 | CAGGTCCATCTTGATCAGCGAATCTCCCACTAGATGGCGGAAGCCACAGCCATAGAAGTA | 300 |
| Pachon...ss_ref | CAGGTCCATCTTGATCAGCGAATCTCCCACTAGATGGCGGAAGCCACAGCCATAGAAGTA | 300 |
| Pachon2B_ref | CAGGTCCATCTTGATCAGCGAATCTCCCACTAGATGGCGGAAGCCACAGCCATAGAAGTA | 300 |
| Pachon...OG_ref | CAGGTCCATCTTGATCAGCGAATCTCCCACTAGATGGCGGAAGCCACAGCCATAGAAGTA | 300 |
| Pach_12_ref | CAGGTCCATCTTGATCAGCGAATCTCCCACTAGATGGCGGAAGCCACAGCCATAGAAGTA | 300 |
| Pach_12 | CAGGTCCATCTTGATCAGCGAATCTCCCACTAGATGGCGGAAGCCACAGCCATAGAAGTA | 300 |
| Pach_17 | CAGGTCCATCTTGATCAGCGAATCTCCCACTAGATGGCGGAAGCCACAGCCATAGAAGTA | 300 |
| Pachon_F2POG | CAGGTCCATCTTGATCAGCGAATCTCCCACTAGATGGCGGAAGCCACAGCCATAGAAGTA | 300 |
| Pachon_3Gross | CAGGTCCATCTTGATCAGCGAATCTCCCACTAGATGGCGGAAGCCACAGCCATAGAAGTA | 300 |
| Pach_15_ref | CAGGTCCATCTTGATCAGCGAATCTCCCACTAGATGGCGGAAGCCACAGCCATAGAAGTA | 300 |
| Pachon...ro_ref | CAGGTCCATCTTGATCAGCGAATCTCCCACTAGATGGCGGAAGCCACAGCCATAGAAGTA | 300 |
| Pachon...ss_ref | CAGGTCCATCTTGATCAGCGAATCTCCCACTAGATGGCGGAAGCCACAGCCATAGAAGTA | 300 |
| Pachon_2Gross | CAGGTCCATCTTGATCAGCGAATCTCCCACTAGATGGCGGAAGCCACAGCCATAGAAGTA | 300 |
| Pach_17_ref | CAGGTCCATCTTGATCAGCGAATCTCCCACTAGATGGCGGAAGCCACAGCCATAGAAGTA | 300 |
| Pachon...ss_ref | CAGGTCCATCTTGATCAGCGAATCTCCCACTAGATGGCGGAAGCCACAGCCATAGAAGTA | 300 |
| Pachon_ref_ref | CAGGTCCATCTTGATCAGCGAATCTCCCACTAGATGGCGGAAGCCACAGCCATAGAAGTA | 300 |
| Pach_8 | CAGGTCCATCTTGATCAGCGAATCTCCCACTAGATGGCGGAAGCCACAGCCATAGAAGTA | 300 |
| Pach_8_ref | CAGGTCCATCTTGATCAGCGAATCTCCCACTAGATGGCGGAAGCCACAGCCATAGAAGTA | 300 |
| Pachon...OG_ref | CAGGTCCATCTTGATCAGCGAATCTCCCACTAGATGGCGGAAGCCACAGCCATAGAAGTA | 300 |
| Pachon_6Boro | CAGGTCCATCTTGATCAGCGAATCTCCCACTAGATGGCGGAAGCCACAGCCATAGAAGTA | 300 |
| Pachon_ref | CAGGTCCATCTTGATCAGCGAATCTCCCACTAGATGGCGGAAGCCACAGCCATAGAAGTA | 300 |
| Pach_3_ref | CAGGTCCATCTTGATCAGCGAATCTCCCACTAGATGGCGGAAGCCACAGCCATAGAAGTA | 300 |
| Pach_9 | CAGGTCCATCTTGATCAGCGAATCTCCCACTAGATGGCGGAAGCCACAGCCATAGAAGTA | 300 |

| | | |
|---|---|---|
| Pachon_G2POG | CAGGTCCATCTTGATCAGCGAATCTCCCACTAGATGGCGGAAGCCACAGCCATAGAAGTA | 300 |
| Pachon...OG_ref | CAGGTCCATCTTGATCAGCGAATCTCCCACTAGATGGCGGAAGCCACAGCCATAGAAGTA | 300 |
| Pach_9_ref | CAGGTCCATCTTGATCAGCGAATCTCCCACTAGATGGCGGAAGCCACAGCCATAGAAGTA | 300 |
| Pachon2B | CAGGTCCATCTTGATCAGCGAATCTCCCACTAGATGGCGGAAGCCACAGCCATAGAAGTA | 300 |
| Pachon...OG_ref | CAGGTCCATCTTGATCAGCGAATCTCCCACTAGATGGCGGAAGCCACAGCCATAGAAGTA | 300 |
| Pachon_E2POG | CAGGTCCATCTTGATCAGCGAATCTCCCACTAGATGGCGGAAGCCACAGCCATAGAAGTA | 300 |
| Pach_14 | CAGGTCCATCTTGATCAGCGAATCTCCCACTAGATGGCGGAAGCCACAGCCATAGAAGTA | 300 |
| Pach_3 | CAGGTCCATCTTGATCAGCGAATCTCCCACTAGATGGCGGAAGCCACAGCCATAGAAGTA | 300 |
| Pachon_H2POG | CAGGTCCATCTTGATCAGCGAATCTCCCACTAGATGGCGGAAGCCACAGCCATAGAAGTA | 300 |
| Pach_7_ref | CAGGTCCATCTTGATCAGCGAATCTCCCACTAGATGGCGGAAGCCACAGCCATAGAAGTA | 300 |
| Pach_15 | CAGGTCCATCTTGATCAGCGAATCTCCCACTAGATGGCGGAAGCCACAGCCATAGAAGTA | 300 |
| Pach_14_ref | CAGGTCCATCTTGATCAGCGAATCTCCCACTAGATGGCGGAAGCCACAGCCATAGAAGTA | 300 |
| Pach_11_ref | CAGGTCCATCTTGATCAGCGAATCTCCCACTAGATGGCGGAAGCCACAGCCATAGAAGTA | 300 |
| Pach_7 | CAGGTCCATCTTGATCAGCGAATCTCCCACTAGATGGCGGAAGCCACAGCCATAGAAGTA | 300 |

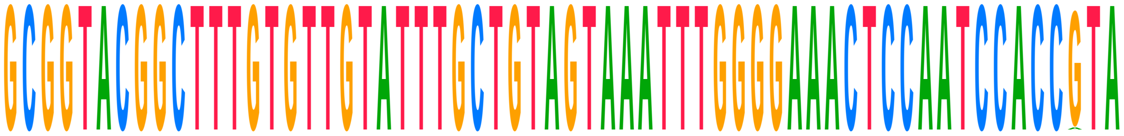

| | | |
|---|---|---|
| **Consensus** | **GCGGTACGGCTTTGTGTTGTATTTGCTGTAGTAAATTTGGGGAAACTCCAATCCACCGTA** | |
| Choy_1_ref | GCGGTACGGCTTTGTGTTGTATTTGCTGTAGTAAATTTGGGGAAACTCCAATCCACCGTA | 360 |
| Choy_1 | GCGGTACGGCTTTGTGTTGTATTTGCTGTAGTAAATTTGGGGAAACTCCAATCCACCGTA | 360 |
| Choy_10_ref | GCGGTACGGCTTTGTGTTGTATTTGCTGTAGTAAATTTGGGGAAACTCCAATCCACCGTA | 360 |
| Choy_10 | GCGGTACGGCTTTGTGTTGTATTTGCTGTAGTAAATTTGGGGAAACTCCAATCCACCGTA | 360 |
| Choy_11_ref | GCGGTACGGCTTTGTGTTGTATTTGCTGTAGTAAATTTGGGGAAACTCCAATCCACCGTA | 342 |
| Choy_11 | GCGGTACGGCTTTGTGTTGTATTTGCTGTAGTAAATTTGGGGAAACTCCAATCCACCGTA | 342 |
| Choy_12_ref | GCGGTACGGCTTTGTGTTGTATTTGCTGTAGTAAATTTGGGGAAACTCCAATCCACCGTA | 360 |
| Choy_12 | GCGGTACGGCTTTGTGTTGTATTTGCTGTAGTAAATTTGGGGAAACTCCAATCCACCGTA | 360 |
| Choy_13_ref | GCGGTACGGCTTTGTGTTGTATTTGCTGTAGTAAATTTGGGGAAACTCCAATCCACCGTA | 360 |
| Choy_13 | GCGGTACGGCTTTGTGTTGTATTTGCTGTAGTAAATTTGGGGAAACTCCAATCCACCGTA | 360 |
| Choy_14_ref | GCGGTACGGCTTTGTGTTGTATTTGCTGTAGTAAATTTGGGGAAACTCCAATCCACCGTA | 360 |
| Choy_14 | GCGGTACGGCTTTGTGTTGTATTTGCTGTAGTAAATTTGGGGAAACTCCAATCCACCGTA | 360 |
| Choy_5_ref | GCGGTACGGCTTTGTGTTGTATTTGCTGTAGTAAATTTGGGGAAACTCCAATCCACCGTA | 360 |
| Choy_5 | GCGGTACGGCTTTGTGTTGTATTTGCTGTAGTAAATTTGGGGAAACTCCAATCCACCGTA | 360 |
| Choy_6_ref | GCGGTACGGCTTTGTGTTGTATTTGCTGTAGTAAATTTGGGGAAACTCCAATCCACCGTA | 360 |
| Choy_6 | GCGGTACGGCTTTGTGTTGTATTTGCTGTAGTAAATTTGGGGAAACTCCAATCCACCGTA | 360 |
| Choy_9_ref | GCGGTACGGCTTTGTGTTGTATTTGCTGTAGTAAATTTGGGGAAACTCCAATCCACCGTA | 360 |
| Choy_9 | GCGGTACGGCTTTGTGTTGTATTTGCTGTAGTAAATTTGGGGAAACTCCAATCCACCATA | 360 |
| Pachon_wild_bco2a | | |
| Pachon_1Gross | GCGGTACGGCTTTGTGTTGTATTTGCTGTAGTAAATTTGGGGAAACTCCAATCCACCGTA | 360 |
| Pach_11 | GCGGTACGGCTTTGTGTTGTATTTGCTGTAGTAAATTTGGGGAAACTCCAATCCACCGTA | 360 |
| Pachon...ss_ref | GCGGTACGGCTTTGTGTTGTATTTGCTGTAGTAAATTTGGGGAAACTCCAATCCACCGTA | 360 |
| Pachon2B_ref | GCGGTACGGCTTTGTGTTGTATTTGCTGTAGTAAATTTGGGGAAACTCCAATCCACCGTA | 360 |
| Pachon...OG_ref | GCGGTACGGCTTTGTGTTGTATTTGCTGTAGTAAATTTGGGGAAACTCCAATCCACCGTA | 360 |
| Pach_12_ref | GCGGTACGGCTTTGTGTTGTATTTGCTGTAGTAAATTTGGGGAAACTCCAATCCACCGTA | 360 |
| Pach_12 | GCGGTACGGCTTTGTGTTGTATTTGCTGTAGTAAATTTGGGGAAACTCCAATCCACCGTA | 360 |
| Pach_17 | GCGGTACGGCTTTGTGTTGTATTTGCTGTAGTAAATTTGGGGAAACTCCAATCCACCGTA | 360 |
| Pachon_F2POG | GCGGTACGGCTTTGTGTTGTATTTGCTGTAGTAAATTTGGGGAAACTCCAATCCACCGTA | 360 |
| Pachon_3Gross | GCGGTACGGCTTTGTGTTGTATTTGCTGTAGTAAATTTGGGGAAACTCCAATCCACCGTA | 360 |
| Pach_15_ref | GCGGTACGGCTTTGTGTTGTATTTGCTGTAGTAAATTTGGGGAAACTCCAATCCACCGTA | 360 |
| Pachon...ro_ref | GCGGTACGGCTTTGTGTTGTATTTGCTGTAGTAAATTTGGGGAAACTCCAATCCACCGTA | 360 |
| Pachon...ss_ref | GCGGTACGGCTTTGTGTTGTATTTGCTGTAGTAAATTTGGGGAAACTCCAATCCACCGTA | 360 |
| Pachon_2Gross | GCGGTACGGCTTTGTGTTGTATTTGCTGTAGTAAATTTGGGGAAACTCCAATCCACCGTA | 360 |
| Pach_17_ref | GCGGTACGGCTTTGTGTTGTATTTGCTGTAGTAAATTTGGGGAAACTCCAATCCACCGTA | 360 |
| Pachon...ss_ref | GCGGTACGGCTTTGTGTTGTATTTGCTGTAGTAAATTTGGGGAAACTCCAATCCACCGTA | 360 |
| Pachon_ref_ref | GCGGTACGGCTTTGTGTTGTATTTGCTGTAGTAAATTTGGGGAAACTCCAATCCACCGTA | 360 |
| Pach_8 | GCGGTACGGCTTTGTGTTGTATTTGCTGTAGTAAATTTGGGGAAACTCCAATCCACCGTA | 360 |
| Pach_8_ref | GCGGTACGGCTTTGTGTTGTATTTGCTGTAGTAAATTTGGGGAAACTCCAATCCACCGTA | 360 |
| Pachon...OG_ref | GCGGTACGGCTTTGTGTTGTATTTGCTGTAGTAAATTTGGGGAAACTCCAATCCACCGTA | 360 |
| Pachon_6Boro | GCGGTACGGCTTTGTGTTGTATTTGCTGTAGTAAATTTGGGGAAACTCCAATCCACCGTA | 360 |

| | | |
|---|---|---|
| Pachon_ref | GCGGTACGGCTTTGTGTTGTATTTGCTGTAGTAAATTTGGGGAAACTCCAATCCACCGTA | 360 |
| Pach_3_ref | GCGGTACGGCTTTGTGTTGTATTTGCTGTAGTAAATTTGGGGAAACTCCAATCCACCGTA | 360 |
| Pach_9 | GCGGTACGGCTTTGTGTTGTATTTGCTGTAGTAAATTTGGGGAAACTCCAATCCACCGTA | 360 |
| Pachon_G2POG | GCGGTACGGCTTTGTGTTGTATTTGCTGTAGTAAATTTGGGGAAACTCCAATCCACCGTA | 360 |
| Pachon...OG_ref | GCGGTACGGCTTTGTGTTGTATTTGCTGTAGTAAATTTGGGGAAACTCCAATCCACCGTA | 360 |
| Pach_9_ref | GCGGTACGGCTTTGTGTTGTATTTGCTGTAGTAAATTTGGGGAAACTCCAATCCACCGTA | 360 |
| Pachon2B | GCGGTACGGCTTTGTGTTGTATTTGCTGTAGTAAATTTGGGGAAACTCCAATCCACCGTA | 360 |
| Pachon...OG_ref | GCGGTACGGCTTTGTGTTGTATTTGCTGTAGTAAATTTGGGGAAACTCCAATCCACCGTA | 360 |
| Pachon_E2POG | GCGGTACGGCTTTGTGTTGTATTTGCTGTAGTAAATTTGGGGAAACTCCAATCCACCGTA | 360 |
| Pach_14 | GCGGTACGGCTTTGTGTTGTATTTGCTGTAGTAAATTTGGGGAAACTCCAATCCACCGTA | 360 |
| Pach_3 | GCGGTACGGCTTTGTGTTGTATTTGCTGTAGTAAATTTGGGGAAACTCCAATCCACCGTA | 360 |
| Pachon_H2POG | GCGGTACGGCTTTGTGTTGTATTTGCTGTAGTAAATTTGGGGAAACTCCAATCCACCGTA | 360 |
| Pach_7_ref | GCGGTACGGCTTTGTGTTGTATTTGCTGTAGTAAATTTGGGGAAACTCCAATCCACCGTA | 360 |
| Pach_15 | GCGGTACGGCTTTGTGTTGTATTTGCTGTAGTAAATTTGGGGAAACTCCAATCCACCGTA | 360 |
| Pach_14_ref | GCGGTACGGCTTTGTGTTGTATTTGCTGTAGTAAATTTGGGGAAACTCCAATCCACCGTA | 360 |
| Pach_11_ref | GCGGTACGGCTTTGTGTTGTATTTGCTGTAGTAAATTTGGGGAAACTCCAATCCACCGTA | 360 |
| Pach_7 | GCGGTACGGCTTTGTGTTGTATTTGCTGTAGTAAATTTGGGGAAACTCCAATCCACCGTA | 360 |

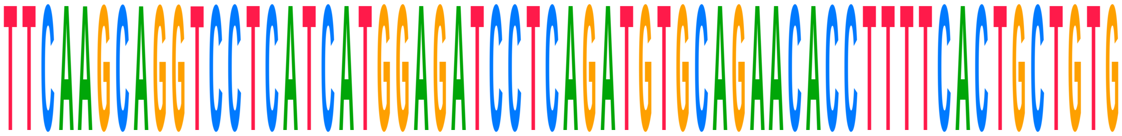

| | | |
|---|---|---|
| **Consensus** | **TTCAAGCAGGTCCTCATCATGGAGATCCTCAGATGTGCAGAACACCTTTTCACTGCTGTG** | |
| Choy_1_ref | TTCAAGCAGGTCCTCATCATGGAGATCCTCAGATGTGCAGAACACCTTTTCACTGCTGTG | 420 |
| Choy_1 | TTCAAGCAGGTCCTCATCATGGAGATCCTCAGATGTGCAGAACACCTTTTCACTGCTGTG | 420 |
| Choy_10_ref | TTCA--------CTCATCATGGAGATCCTCAGATGTGCAGAACACCTTTTCACTGCTGTG | 412 |
| Choy_10 | TTCA--------CTCATCATGGAGATCCTCAGATGTGCAGAACACCTTTTCACTGCTGTG | 412 |
| Choy_11_ref | TTCAAGCAGGTCCTCATCATGGAGATCCTCAGATGTGCAGAACACCTTTTCACTGCTGTG | 402 |
| Choy_11 | TTCAAGCAGGTCCTCATCATGGAGATCCTCAGATGTGCAGAACACCTTTTCACTGCTGTG | 402 |
| Choy_12_ref | TTCAAGCAGGTCCTCATCATGGAGATCCTCAGATGTGCAGAACACCTTTTCACTGCTGTG | 420 |
| Choy_12 | TTCAAGCAGGTCCTCATCATGGAGATCCTCAGATGTGCAGAACACCTTTTCACTGCTGTG | 420 |
| Choy_13_ref | TTCAAGCAGGTCCTCATCATGGAGATCCTCAGATGTGCAGAACACCTTTTCACTGCTGTG | 420 |
| Choy_13 | TTCAAGCAGGTCCTCATCATGGAGATCCTCAGATGTGCAGAACACCTTTTCACTGCTGTG | 420 |
| Choy_14_ref | TTCAAGCAGGTCCTCATCATGGAGATCCTCAGATGTGCAGAACA-CTTTTCACTGCTGTG | 419 |
| Choy_14 | TTCAAGCAGGTCCTCATCATGGAGATCCTCAGATGTGCAGAACA-CTTTTCACTGCTGTG | 419 |
| Choy_5_ref | TTCAAGCAGGTCCTCATCATGGAGATCCTCAGATGTGCAGAACACCTTTTCACTGCTGTG | 420 |
| Choy_5 | TTCAAGCAGGTCCTCATCATGGAGATCCTCAGATGTGCAGAACACCTTTTCACTGCTGTG | 420 |
| Choy_6_ref | TTCAAGCAGGTCCTCATCATGGAGATCCTCAGATGTGCAGAACACCTTTTCACTGCTGTG | 420 |
| Choy_6 | TTCAAGCAGGTCCTCATCATGGAGATCCTCAGATGTGCAGAACACCTTTTCACTGCTGTG | 420 |
| Choy_9_ref | TTCAAGCAGGTCCTCATCATGGAGATCCTCAGATGTGCAGAACACCTTTTCACTGCTGTG | 420 |
| Choy_9 | TTCAAGCAGGTCCTCATCATGGAGATCCTCAGATGTGCAGAACACCTTTTCACTGCTGTG | 420 |
| Pachon_wild_bco2a | | |
| Pachon_1Gross | TTCAAGCAGGTCCTCATCATGGAGATCCTCAGATGTGCAGAACACCTTTTCACTGCTGTG | 420 |
| Pach_11 | TTCAAGCAGGTCCTCATCATGGAGATCCTCAGATGTGCAGAACACCTTTTCACTGCTGTG | 420 |
| Pachon...ss_ref | TTCAAGCAGGTCCTCATCATGGAGATCCTCAGATGTGCAGAACACCTTTTCACTGCTGTG | 420 |
| Pachon2B_ref | TTCAAGCAGGTCCTCATCATGGAGATCCTCAGATGTGCAGAACACCTTTTCACTGCTGTG | 420 |
| Pachon...OG_ref | TTCAAGCAGGTCCTCATCATGGAGATCCTCAGATGTGCAGAACACCTTTTCACTGCTGTG | 420 |
| Pach_12_ref | TTCAAGCAGGTCCTCATCATGGAGATCCTCAGATGTGCAGAACACCTTTTCACTGCTGTG | 420 |
| Pach_12 | TTCAAGCAGGTCCTCATCATGGAGATCCTCAGATGTGCAGAACACCTTTTCACTGCTGTG | 420 |
| Pach_17 | TTCAAGCAGGTCCTCATCATGGAGATCCTCAGATGTGCAGAACACCTTTTCACTGCTGTG | 420 |
| Pachon_F2POG | TTCAAGCAGGTCCTCATCATGGAGATCCTCAGATGTGCAGAACACCTTTTCACTGCTGTG | 420 |
| Pachon_3Gross | TTCAAGCAGGTCCTCATCATGGAGATCCTCAGATGTGCAGAACACCTTTTCACTGCTGTG | 420 |
| Pach_15_ref | TTCAAGCAGGTCCTCATCATGGAGATCCTCAGATGTGCAGAACACCTTTTCACTGCTGTG | 420 |
| Pachon...ro_ref | TTCAAGCAGGTCCTCATCATGGAGATCCTCAGATGTGCAGAACACCTTTTCACTGCTGTG | 420 |
| Pachon...ss_ref | TTCAAGCAGGTCCTCATCATGGAGATCCTCAGATGTGCAGAACACCTTTTCACTGCTGTG | 420 |
| Pachon_2Gross | TTCAAGCAGGTCCTCATCATGGAGATCCTCAGATGTGCAGAACACCTTTTCACTGCTGTG | 420 |
| Pach_17_ref | TTCAAGCAGGTCCTCATCATGGAGATCCTCAGATGTGCAGAACACCTTTTCACTGCTGTG | 420 |
| Pachon...ss_ref | TTCAAGCAGGTCCTCATCATGGAGATCCTCAGATGTGCAGAACACCTTTTCACTGCTGTG | 420 |
| Pachon_ref_ref | TTCAAGCAGGTCCTCATCATGGAGATCCTCAGATGTGCAGAACACCTTTTCACTGCTGTG | 420 |
| Pach_8 | TTCAAGCAGGTCCTCATCATGGAGATCCTCAGATGTGCAGAACACCTTTTCACTGCTGTG | 420 |

| | | |
|---|---|---|
| Pach_8_ref | TTCAAGCAGGTCCTCATCATGGAGATCCTCAGATGTGCAGAACACCTTTTCACTGCTGTG | 420 |
| Pachon...OG_ref | TTCAAGCAGGTCCTCATCATGGAGATCCTCAGATGTGCAGAACACCTTTTCACTGCTGTG | 420 |
| Pachon_6Boro | TTCAAGCAGGTCCTCATCATGGAGATCCTCAGATGTGCAGAACACCTTTTCACTGCTGTG | 420 |
| Pachon_ref | TTCAAGCAGGTCCTCATCATGGAGATCCTCAGATGTGCAGAACACCTTTTCACTGCTGTG | 420 |
| Pach_3_ref | TTCAAGCAGGTCCTCATCATGGAGATCCTCAGATGTGCAGAACACCTTTTCACTGCTGTG | 420 |
| Pach_9 | TTCAAGCAGGTCCTCATCATGGAGATCCTCAGATGTGCAGAACACCTTTTCACTGCTGTG | 420 |
| Pachon_G2POG | TTCAAGCAGGTCCTCATCATGGAGATCCTCAGATGTGCAGAACACCTTTTCACTGCTGTG | 420 |
| Pachon...OG_ref | TTCAAGCAGGTCCTCATCATGGAGATCCTCAGATGTGCAGAACACCTTTTCACTGCTGTG | 420 |
| Pach_9_ref | TTCAAGCAGGTCCTCATCATGGAGATCCTCAGATGTGCAGAACACCTTTTCACTGCTGTG | 420 |
| Pachon2B | TTCAAGCAGGTCCTCATCATGGAGATCCTCAGATGTGCAGAACACCTTTTCACTGCTGTG | 420 |
| Pachon...OG_ref | TTCAAGCAGGTCCTCATCATGGAGATCCTCAGATGTGCAGAACACCTTTTCACTGCTGTG | 420 |
| Pachon_E2POG | TTCAAGCAGGTCCTCATCATGGAGATCCTCAGATGTGCAGAACACCTTTTCACTGCTGTG | 420 |
| Pach_14 | TTCAAGCAGGTCCTCATCATGGAGATCCTCAGATGTGCAGAACACCTTTTCACTGCTGTG | 420 |
| Pach_3 | TTCAAGCAGGTCCTCATCATGGAGATCCTCAGATGTGCAGAACACCTTTTCACTGCTGTG | 420 |
| Pachon_H2POG | TTCAAGCAGGTCCTCATCATGGAGATCCTCAGATGTGCAGAACACCTTTTCACTGCTGTG | 420 |
| Pach_7_ref | TTCAAGCAGGTCCTCATCATGGAGATCCTCAGATGTGCAGAACACCTTTTCACTGCTGTG | 420 |
| Pach_15 | TTCAAGCAGGTCCTCATCATGGAGATCCTCAGATGTGCAGAACACCTTTTCACTGCTGTG | 420 |
| Pach_14_ref | TTCAAGCAGGTCCTCATCATGGAGATCCTCAGATGTGCAGAACACCTTTTCACTGCTGTG | 420 |
| Pach_11_ref | TTCAAGCAGGTCCTCATCATGGAGATCCTCAGATGTGCAGAACACCTTTTCACTGCTGTG | 420 |
| Pach_7 | TTCAAGCAGGTCCTCATCATGGAGATCCTCAGATGTGCAGAACACCTTTTCACTGCTGTG | 420 |

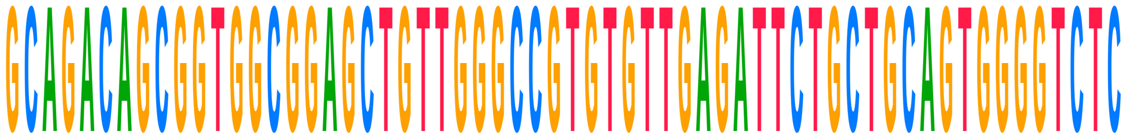

| | | |
|---|---|---|
| **Consensus** | **GCAGACAGCGGTGGCGGAGCTGTTGGGCCGTGTGTTGAGATTCTGCTGCAGTGGGGTCTC** | |
| Choy_1_ref | GCAGACAGCGGTGGCGGAGCTGTTGGGCCGTGTGTTGAGATTCTGCTGCAGTGGGGTCTC | 480 |
| Choy_1 | GCAGACAGCGGTGGCGGAGCTGTTGGGCCGTGTGTTGAGATTCTGCTGCAGTGGGGTCTC | 480 |
| Choy_10_ref | GCAGACAGCGGTGGCGGAGCTGTTGGGCCGTGTGTTGAGATTCTGCTGCAGTGGGGTCTC | 472 |
| Choy_10 | GCAGACAGCGGTGGCGGAGCTGTTGGGCCGTGTGTTGAGATTCTGCTGCAGTGGGGTCTC | 472 |
| Choy_11_ref | GCAGACAGCGGTGGCGGAGCTGTTGGGCCGTGTGTTGAGATTCTGCTGCAGTGGGGTCTC | 462 |
| Choy_11 | GCAGACAGCGGTGGCGGAGCTGTTGGGCCGTGTGTTGAGATTCTGCTGCAGTGGGGTCTC | 462 |
| Choy_12_ref | GCAGACAGCGGTGGCGGAGCTGTTGGGCCGTGTGTTGAGATTCTGCTGCAGTGGGGTCTC | 480 |
| Choy_12 | GCAGACAGCGGTGGCGGAGCTGTTGGGCCGTGTGTTGAGATTCTGCTGCAGTGGGGTCTC | 480 |
| Choy_13_ref | GCAGACAGCGGTGGCGGAGCTGTTGGGCCGTGTGTTGAGATTCTGCTGCAGTGGGGTCTC | 480 |
| Choy_13 | GCAGACAGCGGTGGCGGAGCTGTTGGGCCGTGTGTTGAGATTCTGCTGCAGTGGGGTCTC | 480 |
| Choy_14_ref | GCAGACAGCGGTGGCGGAGCTGTTGGGCCGTGTGTTGAGATTCTGCTGCAGTGGGGTCTC | 479 |
| Choy_14 | GCAGACAGCGGTGGCGGAGCTGTTGGGCCGTGTGTTGAGATTCTGCTGCAGTGGGGTCTC | 479 |
| Choy_5_ref | GCAGACAGCGGTGGCGGAGCTGTTGGGCCGTGTGTTGAGATTCTGCTGCAGTGGGGTCTC | 480 |
| Choy_5 | GCAGACAGCGGTGGCGGAGCTGTTGGGCCGTGTGTTGAGATTCTGCTGCAGTGGGGTCTC | 480 |
| Choy_6_ref | GCAGACAGCGGTGGCGGAGCTGTTGGGCCGTGTGTTGAGATTCTGCTGCAGTGGGGTCTC | 480 |
| Choy_6 | GCAGACAGCGGTGGCGGAGCTGTTGGGCCGTGTGTTGAGATTCTGCTGCAGTGGGGTCTC | 480 |
| Choy_9_ref | GCAGACAGCGGTGGCGGAGCTGTTGGGCCGTGTGTTGAGATTCTGCTGCAGTGGGGTCTC | 480 |
| Choy_9 | GCAGACAGCGGTGGCGGAGCTGTTGGGCCGTGTGTTGAGATTCTGCTGCAGTGGGGTCTC | 480 |
| Pachon_wild_bco2a | | |
| Pachon_1Gross | GCAGACAGCGGTGGCGGAGCTGTTGGGCCGTGTGTTGAGATTCTGCTGCAGTGGGGTCTC | 480 |
| Pach_11 | GCAGACAGCGGTGGCGGAGCTGTTGGGCCGTGTGTTGAGATTCTGCTGCAGTGGGGTCTC | 480 |
| Pachon...ss_ref | GCAGACAGCGGTGGCGGAGCTGTTGGGCCGTGTGTTGAGATTCTGCTGCAGTGGGGTCTC | 480 |
| Pachon2B_ref | GCAGACAGCGGTGGCGGAGCTGTTGGGCCGTGTGTTGAGATTCTGCTGCAGTGGGGTCTC | 480 |
| Pachon...OG_ref | GCAGACAGCGGTGGCGGAGCTGTTGGGCCGTGTGTTGAGATTCTGCTGCAGTGGGGTCTC | 480 |
| Pach_12_ref | GCAGACAGCGGTGGCGGAGCTGTTGGGCCGTGTGTTGAGATTCTGCTGCAGTGGGGTCTC | 480 |
| Pach_12 | GCAGACAGCGGTGGCGGAGCTGTTGGGCCGTGTGTTGAGATTCTGCTGCAGTGGGGTCTC | 480 |
| Pach_17 | GCAGACAGCGGTGGCGGAGCTGTTGGGCCGTGTGTTGAGATTCTGCTGCAGTGGGGTCTC | 480 |
| Pachon_F2POG | GCAGACAGCGGTGGCGGAGCTGTTGGGCCGTGTGTTGAGATTCTGCTGCAGTGGGGTCTC | 480 |
| Pachon_3Gross | GCAGACAGCGGTGGCGGAGCTGTTGGGCCGTGTGTTGAGATTCTGCTGCAGTGGGGTCTC | 480 |
| Pach_15_ref | GCAGACAGCGGTGGCGGAGCTGTTGGGCCGTGTGTTGAGATTCTGCTGCAGTGGGGTCTC | 480 |
| Pachon...ro_ref | GCAGACAGCGGTGGCGGAGCTGTTGGGCCGTGTGTTGAGATTCTGCTGCAGTGGGGTCTC | 480 |
| Pachon...ss_ref | GCAGACAGCGGTGGCGGAGCTGTTGGGCCGTGTGTTGAGATTCTGCTGCAGTGGGGTCTC | 480 |
| Pachon_2Gross | GCAGACAGCGGTGGCGGAGCTGTTGGGCCGTGTGTTGAGATTCTGCTGCAGTGGGGTCTC | 480 |
| Pach_17_ref | GCAGACAGCGGTGGCGGAGCTGTTGGGCCGTGTGTTGAGATTCTGCTGCAGTGGGGTCTC | 480 |

| | | |
|---|---|---|
| Pachon...ss_ref | GCAGACAGCGGTGGCGGAGCTGTTGGGCCGTGTGTTGAGATTCTGCTGCAGTGGGGTCTC | 480 |
| Pachon_ref_ref | GCAGACAGCGGTGGCGGAGCTGTTGGGCCGTGTGTTGAGATTCTGCTGCAGTGGGGTCTC | 480 |
| Pach_8 | GCAGACAGCGGTGGCGGAGCTGTTGGGCCGTGTGTTGAGATTCTGCTGCAGTGGGGTCTC | 480 |
| Pach_8_ref | GCAGACAGCGGTGGCGGAGCTGTTGGGCCGTGTGTTGAGATTCTGCTGCAGTGGGGTCTC | 480 |
| Pachon...OG_ref | GCAGACAGCGGTGGCGGAGCTGTTGGGCCGTGTGTTGAGATTCTGCTGCAGTGGGGTCTC | 480 |
| Pachon_6Boro | GCAGACAGCGGTGGCGGAGCTGTTGGGCCGTGTGTTGAGATTCTGCTGCAGTGGGGTCTC | 480 |
| Pachon_ref | GCAGACAGCGGTGGCGGAGCTGTTGGGCCGTGTGTTGAGATTCTGCTGCAGTGGGGTCTC | 480 |
| Pach_3_ref | GCAGACAGCGGTGGCGGAGCTGTTGGGCCGTGTGTTGAGATTCTGCTGCAGTGGGGTCTC | 480 |
| Pach_9 | GCAGACAGCGGTGGCGGAGCTGTTGGGCCGTGTGTTGAGATTCTGCTGCAGTGGGGTCTC | 480 |
| Pachon_G2POG | GCAGACAGCGGTGGCGGAGCTGTTGGGCCGTGTGTTGAGATTCTGCTGCAGTGGGGTCTC | 480 |
| Pachon...OG_ref | GCAGACAGCGGTGGCGGAGCTGTTGGGCCGTGTGTTGAGATTCTGCTGCAGTGGGGTCTC | 480 |
| Pach_9_ref | GCAGACAGCGGTGGCGGAGCTGTTGGGCCGTGTGTTGAGATTCTGCTGCAGTGGGGTCTC | 480 |
| Pachon2B | GCAGACAGCGGTGGCGGAGCTGTTGGGCCGTGTGTTGAGATTCTGCTGCAGTGGGGTCTC | 480 |
| Pachon...OG_ref | GCAGACAGCGGTGGCGGAGCTGTTGGGCCGTGTGTTGAGATTCTGCTGCAGTGGGGTCTC | 480 |
| Pachon_E2POG | GCAGACAGCGGTGGCGGAGCTGTTGGGCCGTGTGTTGAGATTCTGCTGCAGTGGGGTCTC | 480 |
| Pach_14 | GCAGACAGCGGTGGCGGAGCTGTTGGGCCGTGTGTTGAGATTCTGCTGCAGTGGGGTCTC | 480 |
| Pach_3 | GCAGACAGCGGTGGCGGAGCTGTTGGGCCGTGTGTTGAGATTCTGCTGCAGTGGGGTCTC | 480 |
| Pachon_H2POG | GCAGACAGCGGTGGCGGAGCTGTTGGGCCGTGTGTTGAGATTCTGCTGCAGTGGGGTCTC | 480 |
| Pach_7_ref | GCAGACAGCGGTGGCGGAGCTGTTGGGCCGTGTGTTGAGATTCTGCTGCAGTGGGGTCTC | 480 |
| Pach_15 | GCAGACAGCGGTGGCGGAGCTGTTGGGCCGTGTGTTGAGATTCTGCTGCAGTGGGGTCTC | 480 |
| Pach_14_ref | GCAGACAGCGGTGGCGGAGCTGTTGGGCCGTGTGTTGAGATTCTGCTGCAGTGGGGTCTC | 480 |
| Pach_11_ref | GCAGACAGCGGTGGCGGAGCTGTTGGGCCGTGTGTTGAGATTCTGCTGCAGTGGGGTCTC | 480 |
| Pach_7 | GCAGACAGCGGTGGCGGAGCTGTTGGGCCGTGTGTTGAGATTCTGCTGCAGTGGGGTCTC | 480 |

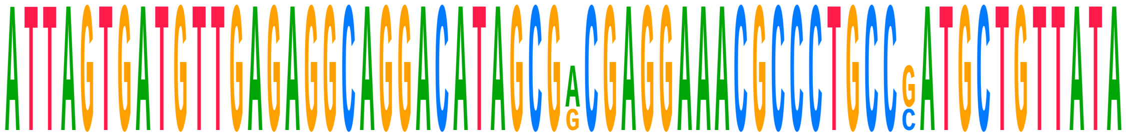

| | | |
|---|---|---|
| **Consensus** | **ATTAGTGATGTTGAGAGGCAGGACATAGCGACGAGGAAACGCCCTGCCGATGCTGTTATA** | |
| Choy_1_ref | ATTAGTGATGTTGAGAGGCAGGACATAGCGGCGAGGAAACGCCCTGCCCATGCTGTTATA | 540 |
| Choy_1 | ATTAGTGATGTTGAGAGGCAGGACATAGCGGCGAGGAAACGCCCTGCCCATGCTGTTATA | 540 |
| Choy_10_ref | ATTAGTGATGTTGAGAGGCAGGACATAGCGGCGAGGAAACGCCCTGCCCATGCTGTTATA | 532 |
| Choy_10 | ATTAGTGATGTTGAGAGGCAGGACATAGCGGCGAGGAAACGCCCTGCCCATGCTGTTATA | 532 |
| Choy_11_ref | ATTAGTGATGTTGAGAGGCAGGACATAGCGGCGAGGAAACGCCCTGCCCATGCTGTTATA | 522 |
| Choy_11 | ATTAGTGATGTTGAGAGGCAGGACATAGCGGCGAGGAAACGCCCTGCCCATGCTGTTATA | 522 |
| Choy_12_ref | ATTAGTGATGTTGAGAGGCAGGACATAGCGGCGAGGAAACGCCCTGCCCATGCTGTTATA | 540 |
| Choy_12 | ATTAGTGATGTTGAGAGGCAGGACATAGCGGCGAGGAAACGCCCTGCCCATGCTGTTATA | 540 |
| Choy_13_ref | ATTAGTGATGTTGAGAGGCAGGACATAGCGGCGAGGAAACGCCCTGCCCATGCTGTTATA | 540 |
| Choy_13 | ATTAGTGATGTTGAGAGGCAGGACATAGCGGCGAGGAAACGCCCTGCCCATGCTGTTATA | 540 |
| Choy_14_ref | ---AGTGATGTTGAGAGGCAGGACATAGCGGCGAGGAAACGCCCTGCCCATGCTGTTATA | 536 |
| Choy_14 | ---AGTGATGTTGAGAGGCAGGACATAGCGGCGAGGAAACGCCCTGCCCATGCTGTTATA | 536 |
| Choy_5_ref | ATTAGTGATGTTGAGAGGCAGGACATAGCGGCGAGGAAACGCCCTGCCCATGCTGTTATA | 540 |
| Choy_5 | ATTAGTGATGTTGAGAGGCAGGACATAGCGGCGAGGAAACGCCCTGCCCATGCTGTTATA | 540 |
| Choy_6_ref | ATTAGTGATGTTGAGAGGCAGGACATAGCGGCGAGGAAACGCCCTGCCCATGCTGTTATA | 540 |
| Choy_6 | ATTAGTGATGTTGAGAGGCAGGACATAGCGGCGAGGAAACGCCCTGCCCATGCTGTTATA | 540 |
| Choy_9_ref | ATTAGTGATGTTGAGAGGCAGGACATAGCGGCGAGGAAACGCCCTGCCCATGCTGTTATA | 540 |
| Choy_9 | ATTAGTGATGTTGAGAGGCAGGACATAGCGGCGAGGAAACGCCCTGCCCATGCTGTTATA | 540 |
| Pachon_wild_bco2a | | |
| Pachon_1Gross | ATTAGTGATGTTGAGAGGCAGGACATAGCGACGAGGAAACGCCCTGCCGATGCTGTTATA | 540 |
| Pach_11 | ATTAGTGATGTTGAGAGGCAGGACATAGCGACGAGGAAACGCCCTGCCGATGCTGTTATA | 540 |
| Pachon...ss_ref | ATTAGTGATGTTGAGAGGCAGGACATAGCGACGAGGAAACGCCCTGCCGATGCTGTTATA | 540 |
| Pachon2B_ref | ATTAGTGATGTTGAGAGGCAGGACATAGCGACGAGGAAACGCCCTGCCGATGCTGTTATA | 540 |
| Pachon...OG_ref | ATTAGTGATGTTGAGAGGCAGGACATAGCGACGAGGAAACGCCCTGCCGATGCTGTTATA | 540 |
| Pach_12_ref | ATTAGTGATGTTGAGAGGCAGGACATAGCGACGAGGAAACGCCCTGCCGATGCTGTTATA | 540 |
| Pach_12 | ATTAGTGATGTTGAGAGGCAGGACATAGCGACGAGGAAACGCCCTGCCGATGCTGTTATA | 540 |
| Pach_17 | ATTAGTGATGTTGAGAGGCAGGACATAGCGACGAGGAAACGCCCTGCCGATGCTGTTATA | 540 |
| Pachon_F2POG | ATTAGTGATGTTGAGAGGCAGGACATAGCGACGAGGAAACGCCCTGCCGATGCTGTTATA | 540 |
| Pachon_3Gross | ATTAGTGATGTTGAGAGGCAGGACATAGCGACGAGGAAACGCCCTGCCGATGCTGTTATA | 540 |
| Pach_15_ref | ATTAGTGATGTTGAGAGGCAGGACATAGCGACGAGGAAACGCCCTGCCGATGCTGTTATA | 540 |
| Pachon...ro_ref | ATTAGTGATGTTGAGAGGCAGGACATAGCGACGAGGAAACGCCCTGCCGATGCTGTTATA | 540 |

| | | |
|---|---|---|
| Pachon...ss_ref | ATTAGTGATGTTGAGAGGCAGGACATAGCGGCGAGGAAACGCCCTGCCCATGCTGTTATA | 540 |
| Pachon_2Gross | ATTAGTGATGTTGAGAGGCAGGACATAGCGACGAGGAAACGCCCTGCCGATGCTGTTATA | 540 |
| Pach_17_ref | ATTAGTGATGTTGAGAGGCAGGACATAGCGACGAGGAAACGCCCTGCCGATGCTGTTATA | 540 |
| Pachon...ss_ref | ATTAGTGATGTTGAGAGGCAGGACATAGCGACGAGGAAACGCCCTGCCGATGCTGTTATA | 540 |
| Pachon_ref_ref | ATTAGTGATGTTGAGAGGCAGGACATAGCGACGAGGAAACGCCCTGCCGATGCTGTTATA | 540 |
| Pach_8 | ATTAGTGATGTTGAGAGGCAGGACATAGCGACGAGGAAACGCCCTGCCGATGCTGTTATA | 540 |
| Pach_8_ref | ATTAGTGATGTTGAGAGGCAGGACATAGCGACGAGGAAACGCCCTGCCGATGCTGTTATA | 540 |
| Pachon...OG_ref | ATTAGTGATGTTGAGAGGCAGGACATAGCGACGAGGAAACGCCCTGCCGATGCTGTTATA | 540 |
| Pachon_6Boro | ATTAGTGATGTTGAGAGGCAGGACATAGCGACGAGGAAACGCCCTGCCGATGCTGTTATA | 540 |
| Pachon_ref | ATTAGTGATGTTGAGAGGCAGGACATAGCGACGAGGAAACGCCCTGCCGATGCTGTTATA | 540 |
| Pach_3_ref | ATTAGTGATGTTGAGAGGCAGGACATAGCGACGAGGAAACGCCCTGCCGATGCTGTTATA | 540 |
| Pach_9 | ATTAGTGATGTTGAGAGGCAGGACATAGCGACGAGGAAACGCCCTGCCGATGCTGTTATA | 540 |
| Pachon_G2POG | ATTAGTGATGTTGAGAGGCAGGACATAGCGACGAGGAAACGCCCTGCCGATGCTGTTATA | 540 |
| Pachon...OG_ref | ATTAGTGATGTTGAGAGGCAGGACATAGCGACGAGGAAACGCCCTGCCGATGCTGTTATA | 540 |
| Pach_9_ref | ATTAGTGATGTTGAGAGGCAGGACATAGCGACGAGGAAACGCCCTGCCGATGCTGTTATA | 540 |
| Pachon2B | ATTAGTGATGTTGAGAGGCAGGACATAGCGACGAGGAAACGCCCTGCCGATGCTGTTATA | 540 |
| Pachon...OG_ref | ATTAGTGATGTTGAGAGGCAGGACATAGCGACGAGGAAACGCCCTGCCGATGCTGTTATA | 540 |
| Pachon_E2POG | ATTAGTGATGTTGAGAGGCAGGACATAGCGACGAGGAAACGCCCTGCCGATGCTGTTATA | 540 |
| Pach_14 | ATTAGTGATGTTGAGAGGCAGGACATAGCGACGAGGAAACGCCCTGCCGATGCTGTTATA | 540 |
| Pach_3 | ATTAGTGATGTTGAGAGGCAGGACATAGCGACGAGGAAACGCCCTGCCGATGCTGTTATA | 540 |
| Pachon_H2POG | ATTAGTGATGTTGAGAGGCAGGACATAGCGACGAGGAAACGCCCTGCCGATGCTGTTATA | 540 |
| Pach_7_ref | ATTAGTGATGTTGAGAGGCAGGACATAGCGACGAGGAAACGCCCTGCCGATGCTGTTATA | 540 |
| Pach_15 | ATTAGTGATGTTGAGAGGCAGGACATAGCGACGAGGAAACGCCCTGCCGATGCTGTTATA | 540 |
| Pach_14_ref | ATTAGTGATGTTGAGAGGCAGGACATAGCGACGAGGAAACGCCCTGCCGATGCTGTTATA | 540 |
| Pach_11_ref | ATTAGTGATGTTGAGAGGCAGGACATAGCGACGAGGAAACGCCCTGCCGATGCTGTTATA | 540 |
| Pach_7 | ATTAGTGATGTTGAGAGGCAGGACATAGCGACGAGGAAACGCCCTGCCGATGCTGTTATA | 540 |

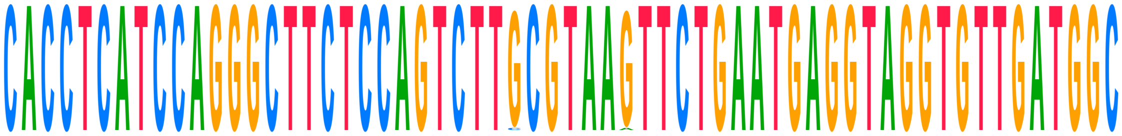

| | | |
|---|---|---|
| **Consensus** | **CACCTCATCCAGGGCTTCTCCAGTCTTGCGTAAGTTCTGAATGAGGTAGGTGTTGATGGC** | |
| Choy_1_ref | CACCTCATCCAGGGCTTCTCCAGTCTTGCGTAAGTTCTGAATGAGGTAGGTGTTGATGGC | 600 |
| Choy_1 | CACCTCATCCAGGGCTTCTCCAGTCTTGCGTAAGTTCTGAATGAGGTAGGTGTTGATGGC | 600 |
| Choy_10_ref | CACCTCATCCAGGGCTTCTCCAGTCTTGCGTAAGTTCTGAATGAGGTAGGTGTTGATGGC | 592 |
| Choy_10 | CACCTCATCCAGGGCTTCTCCAGTCTTCCGTAAATTCTGAATGAGGTAGGTGTTGATGGC | 592 |
| Choy_11_ref | CACCTCATCCAGGGCTTCTCCAGTCTTGCGTAAGTTCTGAATGAGGTAGGTGTTGATGGC | 582 |
| Choy_11 | CACCTCATCCAGGGCTTCTCCAGTCTTGCGTAAGTTCTGAATGAGGTAGGTGTTGATGGC | 582 |
| Choy_12_ref | CACCTCATCCAGGGCTTCTCCAGTCTTGCGTAAGTTCTGAATGAGGTAGGTGTTGATGGC | 600 |
| Choy_12 | CACCTCATCCAGGGCTTCTCCAGTCTTGCGTAAGTTCTGAATGAGGTAGGTGTTGATGGC | 600 |
| Choy_13_ref | CACCTCATCCAGGGCTTCTCCAGTCTTGCGTAAGTTCTGAATGAGGTAGGTGTTGATGGC | 600 |
| Choy_13 | CACCTCATCCAGGGCTTCTCCAGTCTTGCGTAAGTTCTGAATGAGGTAGGTGTTGATGGC | 600 |
| Choy_14_ref | CACCTCATCCAGGGCTTCTCCAGTCTTGCGTAAGTTCTGAATGAGGTAGGTGTTGATGGC | 596 |
| Choy_14 | CACCTCATCCAGGGCTTCTCCAGTCTTGCGTAAGTTCTGAATGAGGTAGGTGTTGATGGC | 596 |
| Choy_5_ref | CACCTCATCCAGGGCTTCTCCAGTCTTGCGTAAGTTCTGAATGAGGTAGGTGTTGATGGC | 600 |
| Choy_5 | CACCTCATCCAGGGCTTCTCCAGTCTTGCGTAAGTTCTGAATGAGGTAGGTGTTGATGGC | 600 |
| Choy_6_ref | CACCTCATCCAGGGCTTCTCCAGTCTTGCGTAAGTTCTGAATGAGGTAGGTGTTGATGGC | 600 |
| Choy_6 | CACCTCATCCAGGGCTTCTCCAGTCTTGCGTAAGTTCTGAATGAGGTAGGTGTTGATGGC | 600 |
| Choy_9_ref | CACCTCATCCAGGGCTTCTCCAGTCTTGCGTAAGTTCTGAATGAGGTAGGTGTTGATGGC | 600 |
| Choy_9 | CACCTCATCCAGGGCTTCTCCAGTCTTGCGTAAGTTCTGAATGAGGTAGGTGTTGATGGC | 600 |
| Pachon_wild_bco2a | | |
| Pachon_1Gross | CACCTCATCCAGGGCTTCTCCAGTCTTGCGTAAGTTCTGAATGAGGTAGGTGTTGATGGC | 600 |
| Pach_11 | CACCTCATCCAGGGCTTCTCCAGTCTTGCGTAAGTTCTGAATGAGGTAGGTGTTGATGGC | 600 |
| Pachon...ss_ref | CACCTCATCCAGGGCTTCTCCAGTCTTGCGTAAGTTCTGAATGAGGTAGGTGTTGATGGC | 600 |
| Pachon2B_ref | CACCTCATCCAGGGCTTCTCCAGTCTTGCGTAAGTTCTGAATGAGGTAGGTGTTGATGGC | 600 |
| Pachon...OG_ref | CACCTCATCCAGGGCTTCTCCAGTCTTGCGTAAGTTCTGAATGAGGTAGGTGTTGATGGC | 600 |
| Pach_12_ref | CACCTCATCCAGGGCTTCTCCAGTCTTGCGTAAGTTCTGAATGAGGTAGGTGTTGATGGC | 600 |
| Pach_12 | CACCTCATCCAGGGCTTCTCCAGTCTTGCGTAAGTTCTGAATGAGGTAGGTGTTGATGGC | 600 |
| Pach_17 | CACCTCATCCAGGGCTTCTCCAGTCTTGCGTAAGTTCTGAATGAGGTAGGTGTTGATGGC | 600 |
| Pachon_F2POG | CACCTCATCCAGGGCTTCTCCAGTCTTGCGTAAGTTCTGAATGAGGTAGGTGTTGATGGC | 600 |

| | | |
|---|---|---|
| Pachon_3Gross | CACCTCATCCAGGGCTTCTCCAGTCTTGCGTAAGTTCTGAATGAGGTAGGTGTTGATGGC | 600 |
| Pach_15_ref | CACCTCATCCAGGGCTTCTCCAGTCTTGCGTAAGTTCTGAATGAGGTAGGTGTTGATGGC | 600 |
| Pachon...ro_ref | CACCTCATCCAGGGCTTCTCCAGTCTTGCGTAAGTTCTGAATGAGGTAGGTGTTGATGGC | 600 |
| Pachon...ss_ref | CACCTCATCCAGGGCTTCTCCAGTCTTGCGTAAGTTCTGAATGAGGTAGGTGTTGATGGC | 600 |
| Pachon_2Gross | CACCTCATCCAGGGCTTCTCCAGTCTTGCGTAAGTTCTGAATGAGGTAGGTGTTGATGGC | 600 |
| Pach_17_ref | CACCTCATCCAGGGCTTCTCCAGTCTTGCGTAAGTTCTGAATGAGGTAGGTGTTGATGGC | 600 |
| Pachon...ss_ref | CACCTCATCCAGGGCTTCTCCAGTCTTGCGTAAGTTCTGAATGAGGTAGGTGTTGATGGC | 600 |
| Pachon_ref_ref | CACCTCATCCAGGGCTTCTCCAGTCTTGCGTAAGTTCTGAATGAGGTAGGTGTTGATGGC | 600 |
| Pach_8 | CACCTCATCCAGGGCTTCTCCAGTCTTGCGTAAGTTCTGAATGAGGTAGGTGTTGATGGC | 600 |
| Pach_8_ref | CACCTCATCCAGGGCTTCTCCAGTCTTGCGTAAGTTCTGAATGAGGTAGGTGTTGATGGC | 600 |
| Pachon...OG_ref | CACCTCATCCAGGGCTTCTCCAGTCTTGCGTAAGTTCTGAATGAGGTAGGTGTTGATGGC | 600 |
| Pachon_6Boro | CACCTCATCCAGGGCTTCTCCAGTCTTGCGTAAGTTCTGAATGAGGTAGGTGTTGATGGC | 600 |
| Pachon_ref | CACCTCATCCAGGGCTTCTCCAGTCTTGCGTAAGTTCTGAATGAGGTAGGTGTTGATGGC | 600 |
| Pach_3_ref | CACCTCATCCAGGGCTTCTCCAGTCTTGCGTAAGTTCTGAATGAGGTAGGTGTTGATGGC | 600 |
| Pach_9 | CACCTCATCCAGGGCTTCTCCAGTCTTGCGTAAGTTCTGAATGAGGTAGGTGTTGATGGC | 600 |
| Pachon_G2POG | CACCTCATCCAGGGCTTCTCCAGTCTTGCGTAAGTTCTGAATGAGGTAGGTGTTGATGGC | 600 |
| Pachon...OG_ref | CACCTCATCCAGGGCTTCTCCAGTCTTGCGTAAGTTCTGAATGAGGTAGGTGTTGATGGC | 600 |
| Pach_9_ref | CACCTCATCCAGGGCTTCTCCAGTCTTGCGTAAGTTCTGAATGAGGTAGGTGTTGATGGC | 600 |
| Pachon2B | CACCTCATCCAGGGCTTCTCCAGTCTTGCGTAAGTTCTGAATGAGGTAGGTGTTGATGGC | 600 |
| Pachon...OG_ref | CACCTCATCCAGGGCTTCTCCAGTCTTGCGTAAGTTCTGAATGAGGTAGGTGTTGATGGC | 600 |
| Pachon_E2POG | CACCTCATCCAGGGCTTCTCCAGTCTTGCGTAAGTTCTGAATGAGGTAGGTGTTGATGGC | 600 |
| Pach_14 | CACCTCATCCAGGGCTTCTCCAGTCTTGCGTAAGTTCTGAATGAGGTAGGTGTTGATGGC | 600 |
| Pach_3 | CACCTCATCCAGGGCTTCTCCAGTCTTGCGTAAGTTCTGAATGAGGTAGGTGTTGATGGC | 600 |
| Pachon_H2POG | CACCTCATCCAGGGCTTCTCCAGTCTTGCGTAAGTTCTGAATGAGGTAGGTGTTGATGGC | 600 |
| Pach_7_ref | CACCTCATCCAGGGCTTCTCCAGTCTTGCGTAAGTTCTGAATGAGGTAGGTGTTGATGGC | 600 |
| Pach_15 | CACCTCATCCAGGGCTTCTCCAGTCTTGCGTAAGTTCTGAATGAGGTAGGTGTTGATGGC | 600 |
| Pach_14_ref | CACCTCATCCAGGGCTTCTCCAGTCTTGCGTAAGTTCTGAATGAGGTAGGTGTTGATGGC | 600 |
| Pach_11_ref | CACCTCATCCAGGGCTTCTCCAGTCTTGCGTAAGTTCTGAATGAGGTAGGTGTTGATGGC | 600 |
| Pach_7 | CACCTCATCCAGGGCTTCTCCAGTCTTGCGTAAGTTCTGAATGAGGTAGGTGTTGATGGC | 600 |

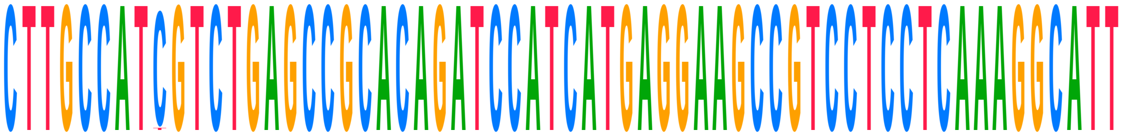

| | | |
|---|---|---|
| **Consensus** | **CTTGCCATCGTCTGAGCCGCACAGATCCATCATGAGGAAGCCGTCCTCCTCAAAGGCATT** | |
| Choy_1_ref | CTTGCCAT--TCTGAGCCGCACAGATCCATCATGAGGAAGCCGTCCTCCTCAAAGGCATT | 658 |
| Choy_1 | CTTGCCAT--TCTGAGCCGCACAGATCCATCATGAGGAAGCCGTCCTCCTCAAAGGCATT | 658 |
| Choy_10_ref | CTTGCCATCGTCTGAGCCGCACAGATCCATCATGAGGAAGCCGTCCTCCTCAAAGGCATT | 652 |
| Choy_10 | CTTGCCATCGTCTGAGCCGCACAGATCCATCATGAGGAAGCCGTCCTCCTCAAAGGCATT | 652 |
| Choy_11_ref | CTTGCCATCGTCTGAGCCGCACAGATCCATCATGAGGAAGCCGTCCTCCTCAAAGGCATT | 642 |
| Choy_11 | CTTGCCATCGTCTGAGCCGCACAGATCCATCATGAGGAAGCCGTCCTCCTCAAAGGCATT | 642 |
| Choy_12_ref | CTTGCCATCGTCTGAGCCGCACAGATCCATCATGAGGAAGCCGTCCTCCTCAAAGGCATT | 660 |
| Choy_12 | CTTGCCATCGTCTGAGCCGCACAGATCCATCATGAGGAAGCCGTCCTCCTCAAAGGCATT | 660 |
| Choy_13_ref | CTTGCCATCGTCTGAGCCGCACAGATCCATCATGAGGAAGCCGTCCTCCTCAAAGGCATT | 660 |
| Choy_13 | CTTGCCATCGTCTGAGCCGCACAGATCCATCATGAGGAAGCCGTCCTCCTCAAAGGCATT | 660 |
| Choy_14_ref | CTTGCCATCGTCTGAGCCGCACAGATCCATCATGAGGAAGCCGTCCTCCTCAAAGGCATT | 656 |
| Choy_14 | CTTGCCATCGTCTGAGCCGCACAGATCCATCATGAGGAAGCCGTCCTCCTCAAAGGCATT | 656 |
| Choy_5_ref | CTTGCCATCGTCTGAGCCGCACAGATCCATCATGAGGAAGCCGTCCTCCTCAAAGGCATT | 660 |
| Choy_5 | CTTGCCATCGTCTGAGCCGCACAGATCCATCATGAGGAAGCCGTCCTCCTCAAAGGCATT | 660 |
| Choy_6_ref | CTTGCCATCGTCTGAGCCGCACAGATCCATCATGAGGAAGCCGTCCTCCTCAAAGGCATT | 660 |
| Choy_6 | CTTGCCATCGTCTGAGCCGCACAGATCCATCATGAGGAAGCCGTCCTCCTCAAAGGCATT | 660 |
| Choy_9_ref | CTTGCCATCGTCTGAGCCGCACAGATCCATCATGAGGAAGCCGTCCTCCTCAAAGGCATT | 660 |
| Choy_9 | CTTGCCATTGTCTGAGCCGCACAGATCCATCATGAGGAAGCCGTCCTCCTCAAAGGCATT | 660 |
| Pachon_wild_bco2a | | |
| Pachon_1Gross | CTTGCCATCGTCTGAGCCGCACAGATCCATCATGAGGAAGCCGTCCTCCTCAAAGGCATT | 660 |
| Pach_11 | CTTGCCATCGTCTGAGCCGCACAGATCCATCATGAGGAAGCCGTCCTCCTCAAAGGCATT | 660 |
| Pachon...ss_ref | CTTGCCATCGTCTGAGCCGCACAGATCCATCATGAGGAAGCCGTCCTCCTCAAAGGCATT | 660 |
| Pachon2B_ref | CTTGCCATCGTCTGAGCCGCACAGATCCATCATGAGGAAGCCGTCCTCCTCAAAGGCATT | 660 |
| Pachon...OG_ref | CTTGCCATCGTCTGAGCCGCACAGATCCATCATGAGGAAGCCGTCCTCCTCAAAGGCATT | 660 |
| Pach_12_ref | CTTGCCATCGTCTGAGCCGCACAGATCCATCATGAGGAAGCCGTCCTCCTCAAAGGCATT | 660 |

| | | |
|---|---|---|
| Pach_12 | CTTGCCATCGTCTGAGCCGCACAGATCCATCATGAGGAAGCCGTCCTCCTCAAAGGCATT | 660 |
| Pach_17 | CTTGCCATCGTCTGAGCCGCACAGATCCATCATGAGGAAGCCGTCCTCCTCAAAGGCATT | 660 |
| Pachon_F2POG | CTTGCCATCGTCTGAGCCGCACAGATCCATCATGAGGAAGCCGTCCTCCTCAAAGGCATT | 660 |
| Pachon_3Gross | CTTGCCATCGTCTGAGCCGCACAGATCCATCATGAGGAAGCCGTCCTCCTCAAAGGCATT | 660 |
| Pach_15_ref | CTTGCCATCGTCTGAGCCGCACAGATCCATCATGAGGAAGCCGTCCTCCTCAAAGGCATT | 660 |
| Pachon…ro_ref | CTTGCCATCGTCTGAGCCGCACAGATCCATCATGAGGAAGCCGTCCTCCTCAAAGGCATT | 660 |
| Pachon…ss_ref | CTTGCCATCGTCTGAGCCGCACAGATCCATCATGAGGAAGCCGTCCTCCTCAAAGGCATT | 660 |
| Pachon_2Gross | CTTGCCATCGTCTGAGCCGCACAGATCCATCATGAGGAAGCCGTCCTCCTCAAAGGCATT | 660 |
| Pach_17_ref | CTTGCCATCGTCTGAGCCGCACAGATCCATCATGAGGAAGCCGTCCTCCTCAAAGGCATT | 660 |
| Pachon…ss_ref | CTTGCCATCGTCTGAGCCGCACAGATCCATCATGAGGAAGCCGTCCTCCTCAAAGGCATT | 660 |
| Pachon_ref_ref | CTTGCCATCGTCTGAGCCGCACAGATCCATCATGAGGAAGCCGTCCTCCTCAAAGGCATT | 660 |
| Pach_8 | CTTGCCATCGTCTGAGCCGCACAGATCCATCATGAGGAAGCCGTCCTCCTCAAAGGCATT | 660 |
| Pach_8_ref | CTTGCCATCGTCTGAGCCGCACAGATCCATCATGAGGAAGCCGTCCTCCTCAAAGGCATT | 660 |
| Pachon…OG_ref | CTTGCCATCGTCTGAGCCGCACAGATCCATCATGAGGAAGCCGTCCTCCTCAAAGGCATT | 660 |
| Pachon_6Boro | CTTGCCATCGTCTGAGCCGCACAGATCCATCATGAGGAAGCCGTCCTCCTCAAAGGCATT | 660 |
| Pachon_ref | CTTGCCATCGTCTGAGCCGCACAGATCCATCATGAGGAAGCCGTCCTCCTCAAAGGCATT | 660 |
| Pach_3_ref | CTTGCCATCGTCTGAGCCGCACAGATCCATCATGAGGAAGCCGTCCTCCTCAAAGGCATT | 660 |
| Pach_9 | CTTGCCATCGTCTGAGCCGCACAGATCCATCATGAGGAAGCCGTCCTCCTCAAAGGCATT | 660 |
| Pachon_G2POG | CTTGCCATCGTCTGAGCCGCACAGATCCATCATGAGGAAGCCGTCCTCCTCAAAGGCATT | 660 |
| Pachon…OG_ref | CTTGCCATCGTCTGAGCCGCACAGATCCATCATGAGGAAGCCGTCCTCCTCAAAGGCATT | 660 |
| Pach_9_ref | CTTGCCATCGTCTGAGCCGCACAGATCCATCATGAGGAAGCCGTCCTCCTCAAAGGCATT | 660 |
| Pachon2B | CTTGCCATCGTCTGAGCCGCACAGATCCATCATGAGGAAGCCGTCCTCCTCAAAGGCATT | 660 |
| Pachon…OG_ref | CTTGCCATCGTCTGAGCCGCACAGATCCATCATGAGGAAGCCGTCCTCCTCAAAGGCATT | 660 |
| Pachon_E2POG | CTTGCCATCGTCTGAGCCGCACAGATCCATCATGAGGAAGCCGTCCTCCTCAAAGGCATT | 660 |
| Pach_14 | CTTGCCATCGTCTGAGCCGCACAGATCCATCATGAGGAAGCCGTCCTCCTCAAAGGCATT | 660 |
| Pach_3 | CTTGCCATCGTCTGAGCCGCACAGATCCATCATGAGGAAGCCGTCCTCCTCAAAGGCATT | 660 |
| Pachon_H2POG | CTTGCCATCGTCTGAGCCGCACAGATCCATCATGAGGAAGCCGTCCTCCTCAAAGGCATT | 660 |
| Pach_7_ref | CTTGCCATCGTCTGAGCCGCACAGATCCATCATGAGGAAGCCGTCCTCCTCAAAGGCATT | 660 |
| Pach_15 | CTTGCCATCGTCTGAGCCGCACAGATCCATCATGAGGAAGCCGTCCTCCTCAAAGGCATT | 660 |
| Pach_14_ref | CTTGCCATCGTCTGAGCCGCACAGATCCATCATGAGGAAGCCGTCCTCCTCAAAGGCATT | 660 |
| Pach_11_ref | CTTGCCATCGTCTGAGCCGCACAGATCCATCATGAGGAAGCCGTCCTCCTCAAAGGCATT | 660 |
| Pach_7 | CTTGCCATCGTCTGAGCCGCACAGATCCATCATGAGGAAGCCGTCCTCCTCAAAGGCATT | 660 |

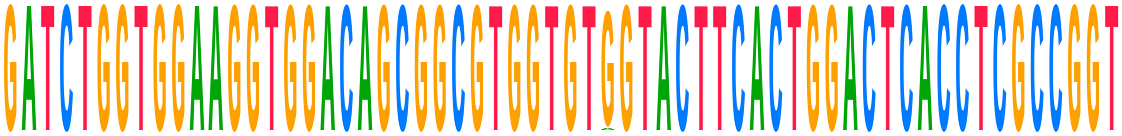

| | | |
|---|---|---|
| **Consensus** | **GATCTGGTGGAAGGTGGACAGCGGCGTGGTGTGGTACTTCACTGGACTCACCTCGCCGGT** | |
| Choy_1_ref | GATCTGGTGGAAGGTGGACAGCGGCGTGGTGTGGTACTTCACTGGACTCACCTCGCCGGT | 718 |
| Choy_1 | GATCTGGTGGAAGGTGGACAGCGGCGTGGTGTGGTACTTCACTGGACTCACCTCGCCGGT | 718 |
| Choy_10_ref | GATCTGGTGGAAGGTGGACAGCGGCGTGGTGTGGTACTTCACTGGACTCACCTCGCCGGT | 712 |
| Choy_10 | GATCTGGTGGAAGGTGGACAGCGGCGTGGTGTGGTACTTCACTGGACTCACCTCGCCGGT | 712 |
| Choy_11_ref | GATCTGGTGGAAGGTGGACAGCGGCGTGGTGTGGTACTTCACTGGACTCACCTCGCCGGT | 702 |
| Choy_11 | GATCTGGTGGAAGGTGGACAGCGGCGTGGTGTGGTACTTCACTGGACTCACCTCGCCGGT | 702 |
| Choy_12_ref | GATCTGGTGGA--------------------------------------ACCTCGCCGGT | 682 |
| Choy_12 | GATCTGGTGGA--------------------------------------ACCTCGCCGGT | 682 |
| Choy_13_ref | GATCTGGTGGAAGGTGGACAGCGGCGTGGTGTGGTACTTCACTGGACTCACCTCGCCGGT | 720 |
| Choy_13 | GATCTGGTGGAAGGTGGACAGCGGCGTGGTGTAGTACTTCACTGGACTCACCTCGCCGGT | 720 |
| Choy_14_ref | GATCTGGTGGAAGGTGGACAGCGGCGTGGTGTGGTACTTCACTGGACTCACCTCGCCGGT | 716 |
| Choy_14 | GATCTGGTGGAAGGTGGACAGCGGCGTGGTGTGGTACTTCACTGGACTCACCTCGCCGGT | 716 |
| Choy_5_ref | GATCTGGTGGAAGGTGGACAGCGGCGTGGTGTGGTACTTCACTGGACTCACCTCGCCGGT | 720 |
| Choy_5 | GATCTGGTGGAAGGTGGACAGCGGCGTGGTGTGGTACTTCACTGGACTCACCTCGCCGGT | 720 |
| Choy_6_ref | GATCTGGTGGAAGGTGGACAGCGGCGTGGTGTGGTACTTCACTGGACTCACCTCGCCGGT | 720 |
| Choy_6 | GATCTGGTGGAAGGTGGACAGCGGCGTGGTGTGGTACTTCACTGGACTCACCTCGCCGGT | 720 |
| Choy_9_ref | GATCTGGTGGAAGGTGGACAGCGGCGTGGTGTGGTACTTCACTGGACTCACCTCGCCGGT | 720 |
| Choy_9 | GATCTGGTGGAAGGTGGACAGCGGCGTGGTGTGGTACTTCACTGGACTCACCTCGCCGGT | 720 |
| Pachon_wild_bco2a | | |
| Pachon_1Gross | GATCTGGTGGAAGGTGGACAGCGGCGTGGTGTGGTACTTCACTGGACTCACCTCGCCGGT | 720 |
| Pach_11 | GATCTGGTGGAAGGTGGACAGCGGCGTGGTGTGGTACTTCACTGGACTCACCTCGCCGGT | 720 |
| Pachon…ss_ref | GATCTGGTGGAAGGTGGACAGCGGCGTGGTGTGGTACTTCACTGGACTCACCTCGCCGGT | 720 |

| | | |
|---|---|---|
| Pachon2B_ref | GATCTGGTGGAAGGTGGACAGCGGCGTGGTGTGGTACTTCACTGGACTCACCTCGCCGGT | 720 |
| Pachon...OG_ref | GATCTGGTGGAAGGTGGACAGCGGCGTGGTGTGGTACTTCACTGGACTCACCTCGCCGGT | 720 |
| Pach_12_ref | GATCTGGTGGAAGGTGGACAGCGGCGTGGTGTGGTACTTCACTGGACTCACCTCGCCGGT | 720 |
| Pach_12 | GATCTGGTGGAAGGTGGACAGCGGCGTGGTGTGGTACTTCACTGGACTCACCTCGCCGGT | 720 |
| Pach_17 | GATCTGGTGGAAGGTGGACAGCGGCGTGGTGTGGTACTTCACTGGACTCACCTCGCCGGT | 720 |
| Pachon_F2POG | GATCTGGTGGAAGGTGGACAGCGGCGTGGTGTGGTACTTCACTGGACTCACCTCGCCGGT | 720 |
| Pachon_3Gross | GATCTGGTGGAAGGTGGACAGCGGCGTGGTGTGGTACTTCACTGGACTCACCTCGCCGGT | 720 |
| Pach_15_ref | GATCTGGTGGAAGGTGGACAGCGGCGTGGTGTGGTACTTCACTGGACTCACCTCGCCGGT | 720 |
| Pachon...ro_ref | GATCTGGTGGAAGGTGGACAGCGGCGTGGTGTGGTACTTCACTGGACTCACCTCGCCGGT | 720 |
| Pachon...ss_ref | GATCTGGTGGAAGGTGGACAGCGGCGTGGTGTGGTACTTCACTGGACTCACCTCGCCGGT | 720 |
| Pachon_2Gross | GATCTGGTGGAAGGTGGACAGCGGCGTGGTGTGGTACTTCACTGGACTCACCTCGCCGGT | 720 |
| Pach_17_ref | GATCTGGTGGAAGGTGGACAGCGGCGTGGTGTGGTACTTCACTGGACTCACCTCGCCGGT | 720 |
| Pachon...ss_ref | GATCTGGTGGAAGGTGGACAGCGGCGTGGTGTGGTACTTCACTGGACTCACCTCGCCGGT | 720 |
| Pachon_ref_ref | GATCTGGTGGAAGGTGGACAGCGGCGTGGTGTGGTACTTCACTGGACTCACCTCGCCGGT | 720 |
| Pach_8 | GATCTGGTGGAAGGTGGACAGCGGCGTGGTGTGGTACTTCACTGGACTCACCTCGCCGGT | 720 |
| Pach_8_ref | GATCTGGTGGAAGGTGGACAGCGGCGTGGTGTGGTACTTCACTGGACTCACCTCGCCGGT | 720 |
| Pachon...OG_ref | GATCTGGTGGAAGGTGGACAGCGGCGTGGTGTGGTACTTCACTGGACTCACCTCGCCGGT | 720 |
| Pachon_6Boro | GATCTGGTGGAAGGTGGACAGCGGCGTGGTGTGGTACTTCACTGGACTCACCTCGCCGGT | 720 |
| Pachon_ref | GATCTGGTGGAAGGTGGACAGCGGCGTGGTGTGGTACTTCACTGGACTCACCTCGCCGGT | 720 |
| Pach_3_ref | GATCTGGTGGAAGGTGGACAGCGGCGTGGTGTGGTACTTCACTGGACTCACCTCGCCGGT | 720 |
| Pach_9 | GATCTGGTGGAAGGTGGACAGCGGCGTGGTGTGGTACTTCACTGGACTCACCTCGCCGGT | 720 |
| Pachon_G2POG | GATCTGGTGGAAGGTGGACAGCGGCGTGGTGTGGTACTTCACTGGACTCACCTCGCCGGT | 720 |
| Pachon...OG_ref | GATCTGGTGGAAGGTGGACAGCGGCGTGGTGTGGTACTTCACTGGACTCACCTCGCCGGT | 720 |
| Pach_9_ref | GATCTGGTGGAAGGTGGACAGCGGCGTGGTGTGGTACTTCACTGGACTCACCTCGCCGGT | 720 |
| Pachon2B | GATCTGGTGGAAGGTGGACAGCGGCGTGGTGTGGTACTTCACTGGACTCACCTCGCCGGT | 720 |
| Pachon...OG_ref | GATCTGGTGGAAGGTGGACAGCGGCGTGGTGTGGTACTTCACTGGACTCACCTCGCCGGT | 720 |
| Pachon_E2POG | GATCTGGTGGAAGGTGGACAGCGGCGTGGTGTGGTACTTCACTGGACTCACCTCGCCGGT | 720 |
| Pach_14 | GATCTGGTGGAAGGTGGACAGCGGCGTGGTGTGGTACTTCACTGGACTCACCTCGCCGGT | 720 |
| Pach_3 | GATCTGGTGGAAGGTGGACAGCGGCGTGGTGTGGTACTTCACTGGACTCACCTCGCCGGT | 720 |
| Pachon_H2POG | GATCTGGTGGAAGGTGGACAGCGGCGTGGTGTGGTACTTCACTGGACTCACCTCGCCGGT | 720 |
| Pach_7_ref | GATCTGGTGGAAGGTGGACAGCGGCGTGGTGTGGTACTTCACTGGACTCACCTCGCCGGT | 720 |
| Pach_15 | GATCTGGTGGAAGGTGGACAGCGGCGTGGTGTGGTACTTCACTGGACTCACCTCGCCGGT | 720 |
| Pach_14_ref | GATCTGGTGGAAGGTGGACAGCGGCGTGGTGTGGTACTTCACTGGACTCACCTCGCCGGT | 720 |
| Pach_11_ref | GATCTGGTGGAAGGTGGACAGCGGCGTGGTGTGGTACTTCACTGGACTCACCTCGCCGGT | 720 |
| Pach_7 | GATCTGGTGGAAGGTGGACAGCGGCGTGGTGTGGTACTTCACTGGACTCACCTCGCCGGT | 720 |

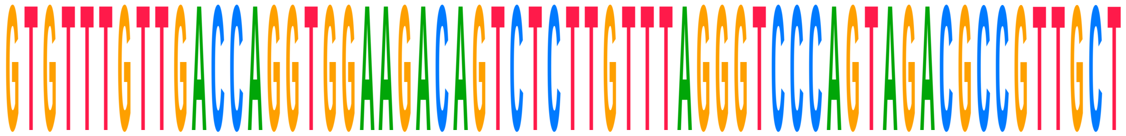

| | | |
|---|---|---|
| **Consensus** | **GTGTTTGTTGACCAGGTGGAAGACAGTCTCTTGTTTAGGGTCCCAGTAGACGCCGTTGCT** | |
| Choy_1_ref | GTGTTTGTTGACCAGGTGGAAGACAGTCTCTTGTTTAGGGTCCCAGTAGACGCCGTTGCT | 778 |
| Choy_1 | GTGTTTGTTGACCAGGTGGAAGACAGTCTCTTGTTTAGGGTCCCAGTAGACGCCGTTGCT | 778 |
| Choy_10_ref | GTGTTTGTTGACCAGGTGGAAGACAGTCTCTTGTTTAGGGTCCCAGTAGACGCCGTTGCT | 772 |
| Choy_10 | GTGTTTGTTGACCAGGTGGAAGACAGTCTCTTGTTTAGGGTCCCAGTAGACGCCGTTGCT | 772 |
| Choy_11_ref | GTGTTTGTTGACCAGGTGGAAGACAGTCTCTTGTTTAGGGTCCCAGTAGACGCCGTTGCT | 762 |
| Choy_11 | GTGTTTGTTGACCAGGTGGAAGACAGTCTCTTGTTTAGGGTCCCAGTAGACGCCGTTGCT | 762 |
| Choy_12_ref | GTGTTTGTTGACCAGGTGGAAGACAGTCTCTTGTTTAGGGTCCCAGTAGACGCCGTTGCT | 742 |
| Choy_12 | GTGTTTGTTGACCAGGTGGAAGACAGTCTCTTGTTTAGGGTCCCAGTAGACGCCGTTGCT | 742 |
| Choy_13_ref | GTGTTTGTTGACCAGGTGGAAGACAGTCTCTTGTTTAGGGTCCCAGTAGACGCCGTTGCT | 780 |
| Choy_13 | GTGTTTGTTGACCAGGTGGAAGACAGTCTCTTGTTTAGGGTCCCAGTAGACGCCGTTGCT | 780 |
| Choy_14_ref | GTGTTTGTTGACCAGGTGGAAGACAGTCTCTTGTTTAGGGTCCCAGTAGACGCCGTTGCT | 776 |
| Choy_14 | GTGTTTGTTGACCAGGTGGAAGACAGTCTCTTGTTTAGGGTCCCAGTAGACGCCGTTGCT | 776 |
| Choy_5_ref | GTGTTTGTTGACCAGGTGGAAGACAGTCTCTTGTTTAGGGTCCCAGTAGACGCCGTTGCT | 780 |
| Choy_5 | GTGTTTGTTGACCAGGTGGAAGACAGTCTCTTGTTTAGGGTCCCAGTAGACGCCGTTGCT | 780 |
| Choy_6_ref | GTGTTTGTTGACCAGGTGGAAGACAGTCTCTTGTTTAGGGTCCCAGTAGACGCCGTTGCT | 780 |
| Choy_6 | GTGTTTGTTGACCAGGTGGAAGACAGTCTCTTGTTTAGGGTCCCAGTAGACGCCGTTGCT | 780 |
| Choy_9_ref | GTGTTTGTTGACCAGGTGGAAGACAGTCTCTTGTTTAGGGTCCCAGTAGACGCCGTTGCT | 780 |
| Choy_9 | GTGTTTGTTGACCAGGTGGAAGACAGTCTCTTGTTTAGGGTCCCAGTAGACGCCGTTGCT | 780 |
| Pachon_wild_bco2a | | |

| | | |
|---|---|---|
| Pachon_1Gross | GTGTTTGTTGACCAGGTGGAAGACAGTCTCTTGTTTAGGGTCCCAGTAGACGCCGTTGCT | 780 |
| Pach_11 | GTGTTTGTTGACCAGGTGGAAGACAGTCTCTTGTTTAGGGTCCCAGTAGACGCCGTTGCT | 780 |
| Pachon...ss_ref | GTGTTTGTTGACCAGGTGGAAGACAGTCTCTTGTTTAGGGTCCCAGTAGACGCCGTTGCT | 780 |
| Pachon2B_ref | GTGTTTGTTGACCAGGTGGAAGACAGTCTCTTGTTTAGGGTCCCAGTAGACGCCGTTGCT | 780 |
| Pachon...OG_ref | GTGTTTGTTGACCAGGTGGAAGACAGTCTCTTGTTTAGGGTCCCAGTAGACGCCGTTGCT | 780 |
| Pach_12_ref | GTGTTTGTTGACCAGGTGGAAGACAGTCTCTTGTTTAGGGTCCCAGTAGACGCCGTTGCT | 780 |
| Pach_12 | GTGTTTGTTGACCAGGTGGAAGACAGTCTCTTGTTTAGGGTCCCAGTAGACGCCGTTGCT | 780 |
| Pach_17 | GTGTTTGTTGACCAGGTGGAAGACAGTCTCTTGTTTAGGGTCCCAGTAGACGCCGTTGCT | 780 |
| Pachon_F2POG | GTGTTTGTTGACCAGGTGGAAGACAGTCTCTTGTTTAGGGTCCCAGTAGACGCCGTTGCT | 780 |
| Pachon_3Gross | GTGTTTGTTGACCAGGTGGAAGACAGTCTCTTGTTTAGGGTCCCAGTAGACGCCGTTGCT | 780 |
| Pach_15_ref | GTGTTTGTTGACCAGGTGGAAGACAGTCTCTTGTTTAGGGTCCCAGTAGACGCCGTTGCT | 780 |
| Pachon...ro_ref | GTGTTTGTTGACCAGGTGGAAGACAGTCTCTTGTTTAGGGTCCCAGTAGACGCCGTTGCT | 780 |
| Pachon...ss_ref | GTGTTTGTTGACCAGGTGGAAGACAGTCTCTTGTTTAGGGTCCCAGTAGACGCCGTTGCT | 780 |
| Pachon_2Gross | GTGTTTGTTGACCAGGTGGAAGACAGTCTCTTGTTTAGGGTCCCAGTAGACGCCGTTGCT | 780 |
| Pach_17_ref | GTGTTTGTTGACCAGGTGGAAGACAGTCTCTTGTTTAGGGTCCCAGTAGACGCCGTTGCT | 780 |
| Pachon...ss_ref | GTGTTTGTTGACCAGGTGGAAGACAGTCTCTTGTTTAGGGTCCCAGTAGACGCCGTTGCT | 780 |
| Pachon_ref_ref | GTGTTTGTTGACCAGGTGGAAGACAGTCTCTTGTTTAGGGTCCCAGTAGACGCCGTTGCT | 780 |
| Pach_8 | GTGTTTGTTGACCAGGTGGAAGACAGTCTCTTGTTTAGGGTCCCAGTAGACGCCGTTGCT | 780 |
| Pach_8_ref | GTGTTTGTTGACCAGGTGGAAGACAGTCTCTTGTTTAGGGTCCCAGTAGACGCCGTTGCT | 780 |
| Pachon...OG_ref | GTGTTTGTTGACCAGGTGGAAGACAGTCTCTTGTTTAGGGTCCCAGTAGACGCCGTTGCT | 780 |
| Pachon_6Boro | GTGTTTGTTGACCAGGTGGAAGACAGTCTCTTGTTTAGGGTCCCAGTAGACGCCGTTGCT | 780 |
| Pachon_ref | GTGTTTGTTGACCAGGTGGAAGACAGTCTCTTGTTTAGGGTCCCAGTAGACGCCGTTGCT | 780 |
| Pach_3_ref | GTGTTTGTTGACCAGGTGGAAGACAGTCTCTTGTTTAGGGTCCCAGTAGACGCCGTTGCT | 780 |
| Pach_9 | GTGTTTGTTGACCAGGTGGAAGACAGTCTCTTGTTTAGGGTCCCAGTAGACGCCGTTGCT | 780 |
| Pachon_G2POG | GTGTTTGTTGACCAGGTGGAAGACAGTCTCTTGTTTAGGGTCCCAGTAGACGCCGTTGCT | 780 |
| Pachon...OG_ref | GTGTTTGTTGACCAGGTGGAAGACAGTCTCTTGTTTAGGGTCCCAGTAGACGCCGTTGCT | 780 |
| Pach_9_ref | GTGTTTGTTGACCAGGTGGAAGACAGTCTCTTGTTTAGGGTCCCAGTAGACGCCGTTGCT | 780 |
| Pachon2B | GTGTTTGTTGACCAGGTGGAAGACAGTCTCTTGTTTAGGGTCCCAGTAGACGCCGTTGCT | 780 |
| Pachon...OG_ref | GTGTTTGTTGACCAGGTGGAAGACAGTCTCTTGTTTAGGGTCCCAGTAGACGCCGTTGCT | 780 |
| Pachon_E2POG | GTGTTTGTTGACCAGGTGGAAGACAGTCTCTTGTTTAGGGTCCCAGTAGACGCCGTTGCT | 780 |
| Pach_14 | GTGTTTGTTGACCAGGTGGAAGACAGTCTCTTGTTTAGGGTCCCAGTAGACGCCGTTGCT | 780 |
| Pach_3 | GTGTTTGTTGACCAGGTGGAAGACAGTCTCTTGTTTAGGGTCCCAGTAGACGCCGTTGCT | 780 |
| Pachon_H2POG | GTGTTTGTTGACCAGGTGGAAGACAGTCTCTTGTTTAGGGTCCCAGTAGACGCCGTTGCT | 780 |
| Pach_7_ref | GTGTTTGTTGACCAGGTGGAAGACAGTCTCTTGTTTAGGGTCCCAGTAGACG-------- | 772 |
| Pach_15 | GTGTTTGTTGACCAGGTGGAAGACAGTCTCTTGTTTAGGGTCCCAGTAGACGCCGTTGCT | 780 |
| Pach_14_ref | GTGTTTGTTGACCAGGTGGAAGACAGTCTCTTGTTTAGGGTCCCAGTAGACGCCGTTGCT | 780 |
| Pach_11_ref | GTGTTTGTTGACCAGGTGGAAGACAGTCTCTTGTTTAGGGTCCCAGTAGACGCCGTTGCT | 780 |
| Pach_7 | GTGTTTGTTGACCAGGTGGAAGACAGTCTCTTGTTTAGGGTCCCAGTAGACG-------- | 772 |

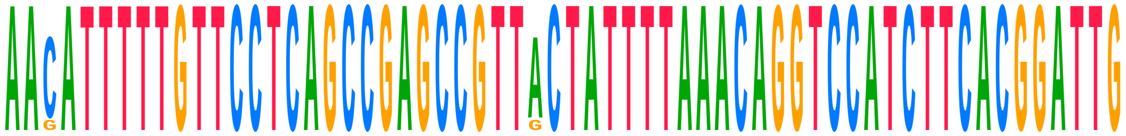

| | | |
|---|---|---|
| **Consensus** | **AACATTTTTGTTCCTCAGCCGAGCCGTTACTATTTTAAACAGGTCCATCTTCACGGATTG** | |
| Choy_1_ref | AACATTTTTGTTCCTCAGCCGAGCCGTTGCTATTTTAAACAGGTCCATCTTCACGGATTG | 838 |
| Choy_1 | AACATTTTTGTTCCTCAGCCGAGCCGTTGCTATTTTAAACAGGTCCATCTTCACGGATTG | 838 |
| Choy_10_ref | AAGATTTTTGTTCCTCAGCCGAGCCGTTACTATTTTAAACAGGTCCATCTTCACGGATTG | 832 |
| Choy_10 | AACATTTTTGTTCCTCAGCCGAGCCGTTGCTATTTTAAACAGGTCCATCTTCACGGATTG | 832 |
| Choy_11_ref | AAGATTTTTGTTCCTCAGCCGAGCCGTTACTATTTTAAACAGGTCCATCTTCACGGATTG | 822 |
| Choy_11 | AAGATTTTTGTTCCTCAGCCGAGCCGTTACTATTTTAAACAGGTCCATCTTCACGGATTG | 822 |
| Choy_12_ref | AAGATTTTTGTTCCTCAGCCGAGCCGTTACTATTTTAAACAGGTCCATCTTCACGGATTG | 802 |
| Choy_12 | AACATTTTTGTTCCTCAGCCGAGCCGTTGCTATTTTAAACAGGTCCATCTTCACGGATTG | 802 |
| Choy_13_ref | AAGATTTTTGTTCCTCAGCCGAGCCGTTACTATTTTAAACAGGTCCATCTTCACGGATTG | 840 |
| Choy_13 | AACATTTTTGTTCCTCAGCCGAGCCGTTACTATTTTAAACAGGTCCATCTTCACGGATTG | 840 |
| Choy_14_ref | AACATTTTTGTTCCTCAGCCGAGCCGTT-CTATTTTAAA---GTCCATCTTCACGGATTG | 832 |
| Choy_14 | AACATTTTTGTTCCTCAGCCGAGCCGTT-CTATTTTAAA---GTCCATCTTCACGGATTG | 832 |
| Choy_5_ref | AACATTTTTGTTCCTCAGCCGAGCCGTTACTATTTTAAACAGGTCCATCTTCACGGATTG | 840 |
| Choy_5 | AACATTTTTGTTCCTCAGCCGAGCCGTTGCTATTTTAAACAGGTCCATCTTCACGGATTG | 840 |
| Choy_6_ref | AACATTTTTGTTCCTCAGCCGAGCCGTTACTATTTTAAACAGGTCCATCTTCACGGATTG | 840 |
| Choy_6 | AACATTTTTGTTCCTCAGCCGAGCCGTTGCTATTTTAAACAGGTCCATCTTCACGGATTG | 840 |

| | | |
|---|---|---|
| Choy_9_ref | AAGATTTTTGTTCCTCAGCCGAGCCGTTACTATTTTAAACAGGTCCATCTTCACGGATTG | 840 |
| Choy_9 | AACATTTTTGTTCCTCAGCCGAGCCGTTACTATTTTAAACAGGTCCATCTTCACGGATTG | 840 |
| Pachon_wild_bco2a | | |
| Pachon_1Gross | AACATTTTTGTTCCTCAGCCGAGCCGTTACTATTTTAAACAGGTCCATCTTCACGGATTG | 840 |
| Pach_11 | AACATTTTTGTTCCTCAGCCGAGCCGTTACTATTTTAAACAGGTCCATCTTCACGGATTG | 840 |
| Pachon...ss_ref | AACATTTTTGTTCCTCAGCCGAGCCGTTACTATTTTAAACAGGTCCATCTTCACGGATTG | 840 |
| Pachon2B_ref | AACATTTTTGTTCCTCAGCCGAGCCGTTACTATTTTAAACAGGTCCATCTTCACGGATTG | 840 |
| Pachon...OG_ref | AACATTTTTGTTCCTCAGCCGAGCCGTTACTATTTTAAACAGGTCCATCTTCACGGATTG | 840 |
| Pach_12_ref | AACA--------CCTCAGCCGAGCCGTTACTATTTTAAACAGGTCCATCTTCACGGATTG | 832 |
| Pach_12 | AACA--------CCTCAGCCGAGCCGTTACTATTTTAAACAGGTCCATCTTCACGGATTG | 832 |
| Pach_17 | AACATTTTTGTTCCTCAGCCGAGCCGTTACTATTTTAAACAGGTCCATCTTCACGGATTG | 840 |
| Pachon_F2POG | AACATTTTTGTTCCTCAGCCGAGCCGTTACTATTTTAAACAGGTCCATCTTCACGGATTG | 840 |
| Pachon_3Gross | AACATTTTTGTTCCTCAGCCGAGCCGTTACTATTTTAAACAGGTCCATCTTCACGGATTG | 840 |
| Pach_15_ref | AACATTTTTGTTCCTCAGCCGAGCCGTTACTATTTTAAACAGGTCCATCTTCACGGATTG | 840 |
| Pachon...ro_ref | AACATTTTTGTTCCTCAGCCGAGCCGTTACTATTTTAAACAGGTCCATCTTCACGGATTG | 840 |
| Pachon...ss_ref | AACATTTTTGTTCCTCAGCCGAGCCGTTACTATTTTAAACAGGTCCATCTTCACGGATTG | 840 |
| Pachon_2Gross | AACATTTTTGTTCCTCAGCCGAGCCGTTACTATTTTAAACAGGTCCATCTTCACGGATTG | 840 |
| Pach_17_ref | AACATTTTTGTTCCTCAGCCGAGCCGTTACTATTTTAAACAGGTCCATCTTCACGGATTG | 840 |
| Pachon...ss_ref | AACATTTTTGTTCCTCAGCCGAGCCGTTACTATTTTAAACAGGTCCATCTTCACGGATTG | 840 |
| Pachon_ref_ref | AACATTTTTGTTCCTCAGCCGAGCCGTTACTATTTTAAACAGGTCCATCTTCACGGATTG | 840 |
| Pach_8 | AACATTTTTGTTCCTCAGCCGAGCCGTTACTATTTTAAACAGGTCCATCTTCACGGATTG | 840 |
| Pach_8_ref | AACATTTTTGTTCCTCAGCCGAGCCGTTACTATTTTAAACAGGTCCATCTTCACGGATTG | 840 |
| Pachon...OG_ref | AACATTTTTGTTCCTCAGCCGAGCCGTTACTATTTTAAACAGGTCCATCTTCACGGATTG | 840 |
| Pachon_6Boro | AACATTTTTGTTCCTCAGCCGAGCCGTTACTATTTTAAACAGGTCCATCTTCACGGATTG | 840 |
| Pachon_ref | AACATTTTTGTTCCTCAGCCGAGCCGTTACTATTTTAAACAGGTCCATCTTCACGGATTG | 840 |
| Pach_3_ref | AACATTTTTGTTCCTCAGCCGAGCCGTTACTATTTTAAACAGGTCCATCTTCACGGATTG | 840 |
| Pach_9 | AACATTTTTGTTCCTCAGCCGAGCCGTTACTATTTTAAACAGGTCCATCTTCACGGATTG | 840 |
| Pachon_G2POG | AACATTTTTGTTCCTCAGCCGAGCCGTTACTATTTTAAACAGGTCCATCTTCACGGATTG | 840 |
| Pachon...OG_ref | AACATTTTTGTTCCTCAGCCGAGCCGTTACTATTTTAAACAGGTCCATCTTCACGGATTG | 840 |
| Pach_9_ref | AACATTTTTGTTCCTCAGCCGAGCCGTTACTATTTTAAACAGGTCCATCTTCACGGATTG | 840 |
| Pachon2B | AACATTTTTGTTCCTCAGCCGAGCCGTTACTATTTTAAACAGGTCCATCTTCACGGATTG | 840 |
| Pachon...OG_ref | AACATTTTTGTTCCTCAGCCGAGCCGTTACTATTTTAAACAGGTCCATCTTCACGGATTG | 840 |
| Pachon_E2POG | AACATTTTTGTTCCTCAGCCGAGCCGTTACTATTTTAAACAGGTCCATCTTCACGGATTG | 840 |
| Pach_14 | AACATTTTTGTTCCTCAGCCGAGCCGTTACTATTTTAAACAGGTCCATCTTCACGGATTG | 840 |
| Pach_3 | AACATTTTTGTTCCTCAGCCGAGCCGTTACTATTTTAAACAGGTCCATCTTCACGGATTG | 840 |
| Pachon_H2POG | AACATTTTTGTTCCTCAGCCGAGCCGTTACTATTTTAAACAGGTCCATCTTCACGGATTG | 840 |
| Pach_7_ref | -----TTTTGTTCCTCAGCCGAGCCGTTACTATTTTAAACAGGTCCATCTTCACGGATTG | 827 |
| Pach_15 | AACATTTTTGTTCCTCAGCCGAGCCGTTACTATTTTAAACAGGTCCATCTTCACGGATTG | 840 |
| Pach_14_ref | AACATTTTTGTTCCTCAGCCGAGCCGTTACTATTTTAAACAGGTCCATCTTCACGGATTG | 840 |
| Pach_11_ref | AACATTTTTGTTCCTCAGCCGAGCCGTTACTATTTTAAACAGGTCCATCTTCACGGATTG | 840 |
| Pach_7 | -----TTTTGTTCCTCAGCCGAGCCGTTACTATTTTAAACAGGTCCATCTTCACGGATTG | 827 |

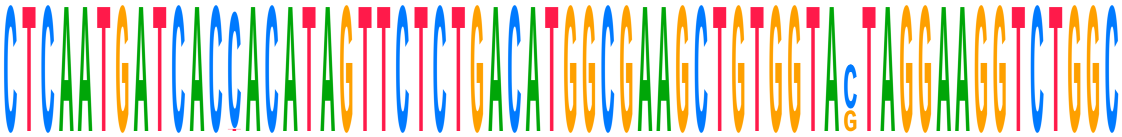

| | | |
|---|---|---|
| **Consensus** | **CTCAATGATCACCACATAGTTCTCTGACATGGCGAAGCTGTGGTACTAGGAAGGTCTGGC** | |
| Choy_1_ref | CTCAATGATCACCACATAGTTCTCTGACATGGCGAAGCTGTGGTAGTAGGAAGGTCTGGC | 898 |
| Choy_1 | CTCAATGATCACCACATAGTTCTCTGACATGGCGAAGCTGTGGTAGTAGGAAGGTCTGGC | 898 |
| Choy_10_ref | CTCAATGATCACCACATAGTTCTCTGACATGGCGAAGCTGTGGTAGTAGGAAGGTCTGGC | 892 |
| Choy_10 | CTCAATGATCACCACATAGTTCTCTGACATGGCGAAGCTGTGGTAGTAGGAAGGTCTGGC | 892 |
| Choy_11_ref | CTCAATGATCACCACATAGTTCTCTGACATGGCGAAGCTGTGGTAGTAGGAAGGTCTGGC | 882 |
| Choy_11 | CTCAATGATCACCACATAGTTCTCTGACATGGCGAAGCTGTGGTAGTAGGAAGGTCTGGC | 882 |
| Choy_12_ref | CTCAATGATCACCACATAGTTCTCTGACATGGCGAAGCTGTGGTAGTAGGAAGGTCTGGC | 862 |
| Choy_12 | CTCAATGATCACCACATAGTTCTCTGACATGGCGAAGCTGTGGTAGTAGGAAGGTCTGGC | 862 |
| Choy_13_ref | CTCAATGATCACCACATAGTTCTCTGACATGGCGAAGCTGTGGTAGTAGGAAGGTCTGGC | 900 |
| Choy_13 | CTCAATGATCACTACATAGTTCTCTGACATGGCGAAGCTGTGGTAGTAGGAAGGTCTGGC | 900 |
| Choy_14_ref | CTCAATGATCACCACATAGTTCTCTGACATGGCGAAGCTGTGGTAGTAGGAAGGTCTGGC | 892 |
| Choy_14 | CTCAATGATCACCACATAGTTCTCTGACATGGCGAAGCTGTGGTAGTAGGAAGGTCTGGC | 892 |
| Choy_5_ref | CTCAATGATCACCACATAGTTCTCTGACATGGCGAAGCTGTGGTAGTAGGAAGGTCTGGC | 900 |

| | | |
|---|---|---|
| Choy_5 | CTCAATGATCACCACATAGTTCTCTGACATGGCGAAGCTGTGGTAGTAGGAAGGTCTGGC | 900 |
| Choy_6_ref | CTCAATGATCACCACATAGTTCTCTGACATGGCGAAGCTGTGGTAGTAGGAAGGTCTGGC | 900 |
| Choy_6 | CTCAATGATCACCACATAGTTCTCTGACATGGCGAAGCTGTGGTAGTAGGAAGGTCTGGC | 900 |
| Choy_9_ref | CTCAATGATCACCACATAGTTCTCTGACATGGCGAAGCTGTGGTAGTAGGAAGGTCTGGC | 900 |
| Choy_9 | CTCAATGATCACCACATAGTTCTCTGACATGGCGAAGCTGTGGTAGTAGGAAGGTCTGGC | 900 |
| Pachon_wild_bco2a | | |
| Pachon_1Gross | CTCAATGATCACCACATAGTTCTCTGACATGGCGAAGCTGTGGTACTAGGAAGGTCTGGC | 900 |
| Pach_11 | CTCAATGATCACCACATAGTTCTCTGACATGGCGAAGCTGTGGTACTAGGAAGGTCTGGC | 900 |
| Pachon…ss_ref | CTCAATGATCACCACATAGTTCTCTGACATGGCGAAGCTGTGGTACTAGGAAGGTCTGGC | 900 |
| Pachon2B_ref | CTCAATGATCACCACATAGTTCTCTGACATGGCGAAGCTGTGGTACTAGGAAGGTCTGGC | 900 |
| Pachon…OG_ref | CTCAATGATCACCACATAGTTCTCTGACATGGCGAAGCTGTGGTACTAGGAAGGTCTGGC | 900 |
| Pach_12_ref | CTCAATGATCACCACATAGTTCTCTGACATGGCGAAGCTGTGGTACTAGGAAGGTCTGGC | 892 |
| Pach_12 | CTCAATGATCACCACATAGTTCTCTGACATGGCGAAGCTGTGGTACTAGGAAGGTCTGGC | 892 |
| Pach_17 | CTCAATGATCACCACATAGTTCTCTGACATGGCGAAGCTGTGGTACTAGGAAGGTCTGGC | 900 |
| Pachon_F2POG | CTCAATGATCACCACATAGTTCTCTGACATGGCGAAGCTGTGGTACTAGGAAGGTCTGGC | 900 |
| Pachon_3Gross | CTCAATGATCACCACATAGTTCTCTGACATGGCGAAGCTGTGGTACTAGGAAGGTCTGGC | 900 |
| Pach_15_ref | CTCAATGATCACCACATAGTTCTCTGACATGGCGAAGCTGTGGTACTAGGAAGGTCTGGC | 900 |
| Pachon…ro_ref | CTCAATGATCACCACATAGTTCTCTGACATGGCGAAGCTGTGGTACTAGGAAGGTCTGGC | 900 |
| Pachon…ss_ref | CTCAATGATCACCACATAGTTCTCTGACATGGCGAAGCTGTGGTACTAGGAAGGTCTGGC | 900 |
| Pachon_2Gross | CTCAATGATCACCACATAGTTCTCTGACATGGCGAAGCTGTGGTACTAGGAAGGTCTGGC | 900 |
| Pach_17_ref | CTCAATGATCACCACATAGTTCTCTGACATGGCGAAGCTGTGGTACTAGGAAGGTCTGGC | 900 |
| Pachon…ss_ref | CTCAATGATCACCACATAGTTCTCTGACATGGCGAAGCTGTGGTACTAGGAAGGTCTGGC | 900 |
| Pachon_ref_ref | CTCAATGATCACCACATAGTTCTCTGACATGGCGAAGCTGTGGTACTAGGAAGGTCTGGC | 900 |
| Pach_8 | CTCAATGATCACCACATAGTTCTCTGACATGGCGAAGCTGTGGTACTAGGAAGGTCTGGC | 900 |
| Pach_8_ref | CTCAATGATCACCACATAGTTCTCTGACATGGCGAAGCTGTGGTACTAGGAAGGTCTGGC | 900 |
| Pachon…OG_ref | CTCAATGATCACCACATAGTTCTCTGACATGGCGAAGCTGTGGTACTAGGAAGGTCTGGC | 900 |
| Pachon_6Boro | CTCAATGATCACCACATAGTTCTCTGACATGGCGAAGCTGTGGTACTAGGAAGGTCTGGC | 900 |
| Pachon_ref | CTCAATGATCACCACATAGTTCTCTGACATGGCGAAGCTGTGGTACTAGGAAGGTCTGGC | 900 |
| Pach_3_ref | CTCAATGATCACCACATAGTTCTCTGACATGGCGAAGCTGTGGTACTAGGAAGGTCTGGC | 900 |
| Pach_9 | CTCAATGATCACCACATAGTTCTCTGACATGGCGAAGCTGTGGTACTAGGAAGGTCTGGC | 900 |
| Pachon_G2POG | CTCAATGATCACCACATAGTTCTCTGACATGGCGAAGCTGTGGTACTAGGAAGGTCTGGC | 900 |
| Pachon…OG_ref | CTCAATGATCACCACATAGTTCTCTGACATGGCGAAGCTGTGGTACTAGGAAGGTCTGGC | 900 |
| Pach_9_ref | CTCAATGATCACCACATAGTTCTCTGACATGGCGAAGCTGTGGTACTAGGAAGGTCTGGC | 900 |
| Pachon2B | CTCAATGATCACCACATAGTTCTCTGACATGGCGAAGCTGTGGTACTAGGAAGGTCTGGC | 900 |
| Pachon…OG_ref | CTCAATGATCACCACATAGTTCTCTGACATGGCGAAGCTGTGGTACTAGGAAGGTCTGGC | 900 |
| Pachon_E2POG | CTCAATGATCACCACATAGTTCTCTGACATGGCGAAGCTGTGGTACTAGGAAGGTCTGGC | 900 |
| Pach_14 | CTCAATGATCACCACATAGTTCTCTGACATGGCGAAGCTGTGGTACTAGGAAGGTCTGGC | 900 |
| Pach_3 | CTCAATGATCACCACATAGTTCTCTGACATGGCGAAGCTGTGGTACTAGGAAGGTCTGGC | 900 |
| Pachon_H2POG | CTCAATGATCACCACATAGTTCTCTGACATGGCGAAGCTGTGGTACTAGGAAGGTCTGGC | 900 |
| Pach_7_ref | CTCAATGATCACCACATAGTTCTCTGACATGGCGAAGCTGTGGTACTAGGAAGGTCTGGC | 887 |
| Pach_15 | CTCAATGATCACCACATAGTTCTCTGACATGGCGAAGCTGTGGTACTAGGAAGGTCTGGC | 900 |
| Pach_14_ref | CTCAATGATCACCACATAGTTCTCTGACATGGCGAAGCTGTGGTACTAGGAAGGTCTGGC | 900 |
| Pach_11_ref | CTCAATGATCACCACATAGTTCTCTGACATGGCGAAGCTGTGGTACTAGGAAGGTCTGGC | 900 |
| Pach_7 | CTCAATGATCACCACATAGTTCTCTGACATGGCGAAGCTGTGGTACTAGGAAGGTCTGGC | 887 |

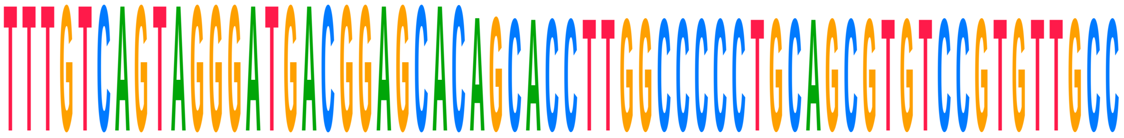


| | | |
|---|---|---|
| **Consensus** | **TTTGTCAGTAGGGATGACGGAGCAC-----------------------------------** |  |
| Choy_1_ref | TTTGTCAGTAGGGATGACGGAGCACAGCACCTTGGCCCCCTGCAGCGTGTCCGTGTTGCC | 958 |
| Choy_1 | TTTGTCAGTAGGGATGACGGAGCACAGCACCTTGGCCCCCTGCAGCGTGTCCGTGTTGCC | 958 |
| Choy_10_ref | TTTGTCAGTAGGGATGACGGAGCACAGCACCTTGGCCCCCTGCAGCGTGTCCGTGTTGCC | 952 |
| Choy_10 | TTTGTCAGTAGGGATGACGGAGCACAGCACCTTGGCCCCCTGCAGCGTGTCCGTGTTGCC | 952 |
| Choy_11_ref | TTTGTCAGTAGGGATGACGGAGCACAGCACCTTGGCCCCCTGCAGCGTGTCCGTGTTGCC | 942 |
| Choy_11 | TTTGTCAGTAGGGATGACGGAGCACAGCACCTTGGCCCCCTGCAGCGTGTCCGTGTTGCC | 942 |
| Choy_12_ref | TTTGTCAGTAGGGATGACGGAGCACAGCACCTTGGCCCCCTGCAGCGTGTCCGTGTTGCC | 922 |
| Choy_12 | TTTGTCAGTAGGGATGACGGAGCACAGCACCTTGGCCCCCTGCAGCGTGTCCGTGTTGCC | 922 |
| Choy_13_ref | TTTGTCAGTAGGGATGACGGAGCACAGCACCTTGGCCCCCTGCAGCGTGTCCGTGTTGCC | 960 |
| Choy_13 | TTTGTCAGTAGGGATGACGGAGCACAGCACCTTGGCCCCCTGCAGCGTGTCCGTGTTGCC | 960 |

```
Choy_14_ref          TTTGTCAGTAGGGATGACGGAGCACAGCACCTTGGCCCCCTGCAGCGTGTCCGTGTTGCC  952
Choy_14              TTTGTCAGTAGGGATGACGGAGCACAGCACCTTGGCCCCCTGCAGCGTGTCCGTGTTGCC  952
Choy_5_ref           TTTGTCAGTAGGGATGACGGAGCACAGCACCTTGGCCCCCTGCAGCGTGTCCGTGTTGCC  960
Choy_5               TTTGTCAGTAGGGATGACGGAGCACAGCACCTTGGCCCCCTGCAGCGTGTCCGTGTTGCC  960
Choy_6_ref           TTTGTCAGTAGGGATGACGGAGCACAGCACCTTGGCCCCCTGCAGCGTGTCCGTGTTGCC  960
Choy_6               TTTGTCAGTAGGGATGACGGAGCACAGCACCTTGGCCCCCTGCAGCGTGTCCGTGTTGCC  960
Choy_9_ref           TTTGTCAGTAGGGATGACGGAGCACAGCACCTTGGCCCCCTGCAGCGTGTCCGTGTTGCC  960
Choy_9               TTTGTCAGTAGGGATGACGGAGCACAGCACCTTGGCCCCCTGCAGCGTGTCCGTGTTGCC  960
Pachon_wild_bco2a
   Pachon_1Gross     TTTGTCAGTAGGGATGACGGAGCAC-----------------------------------  925
   Pach_11           TTTGTCAGTAGGGATGACGGAGCACAG---------------------------------  927
   Pachon...ss_ref   TTTGTCAGTAGGGATGACGGAGCAC-----------------------------------  925
   Pachon2B_ref      TTTGTCAGTAGGGATGACGGAGCAC-----------------------------------  925
   Pachon...OG_ref   TTTGTCAGTAGGGATGACGGAGCAC-----------------------------------  925
   Pach_12_ref       TTTGTCAGTAGGGATGACGGAGCAC-----------------------------------  917
   Pach_12           TTTGTCAGTAGGGATGACGGAGCAC-----------------------------------  917
   Pach_17           TTTGTCAGTAGGGATGACGGAGCAC-----------------------------------  925
   Pachon_F2POG      TTTGTCAGTAGGGATGACGGAGCAC-----------------------------------  925
   Pachon_3Gross     TTTGTCAGTAGGGATGACGGAGCAC-----------------------------------  925
   Pach_15_ref       TTTGTCAGTAGGGATGACGGAGCAC-----------------------------------  925
   Pachon...ro_ref   TTTGTCAGTAGGGATGACGGAGCAC-----------------------------------  925
   Pachon...ss_ref   TTTGTCAGTAGGGATGACGGAGCAC-----------------------------------  925
   Pachon_2Gross     TTTGTCAGTAGGGATGACGGAGCAC-----------------------------------  925
   Pach_17_ref       TTTGTCAGTAGGGATGACGGAGCAC-----------------------------------  925
   Pachon...ss_ref   TTTGTCAGTAGGGATGACGGAGCAC-----------------------------------  925
   Pachon_ref_ref    TTTGTCAGTAGGGATGACGGAGCAC-----------------------------------  925
   Pach_8            TTTGTCAGTAGGGATGACGGAGCAC-----------------------------------  925
   Pach_8_ref        TTTGTCAGTAGGGATGACGGAGCAC-----------------------------------  925
   Pachon...OG_ref   TTTGTCAGTAGGGATGACGGAGCAC-----------------------------------  925
   Pachon_6Boro      TTTGTCAGTAGGGATGACGGAGCAC-----------------------------------  925
   Pachon_ref        TTTGTCAGTAGGGATGACGGAGCAC-----------------------------------  925
   Pach_3_ref        TTTGTCAGTAGGGATGACGGAGCAC-----------------------------------  925
   Pach_9            TTTGTCAGTAGGGATGACGGAGCAC-----------------------------------  925
   Pachon_G2POG      TTTGTCAGTAGGGATGACGGAGCAC-----------------------------------  925
   Pachon...OG_ref   TTTGTCAGTAGGGATGACGGAGCAC-----------------------------------  925
   Pach_9_ref        TTTGTCAGTAGGGATGACGGAGCAC-----------------------------------  925
   Pachon2B          TTTGTCAGTAGGGATGACGGAGCAC-----------------------------------  925
   Pachon...OG_ref   TTTGTCAGTAGGGATGACGGAGCAC-----------------------------------  925
   Pachon_E2POG      TTTGTCAGTAGGGATGACGGAGCAC-----------------------------------  925
   Pach_14           TTTGTCAGTAGGGATGACGGAGCAC-----------------------------------  925
   Pach_3            TTTGTCAGTAGGGATGACGGAGCAC-----------------------------------  925
   Pachon_H2POG      TTTGTCAGTAGGGATGACGGAGCAC-----------------------------------  925
   Pach_7_ref        TTTGTCAGTAGGGATGACGGAGCA------------------------------------  911
   Pach_15           TTTGTCAGTAGGGATGACGGAGCAC-----------------------------------  925
   Pach_14_ref       TTTGTCAGTAGGGATGACGGAGCAC-----------------------------------  925
   Pach_11_ref       TTTGTCAGTAGGGATGACGGAGCACAG---------------------------------  927
   Pach_7            TTTGTCAGTAGGGATGACGGAGCA------------------------------------  911
```

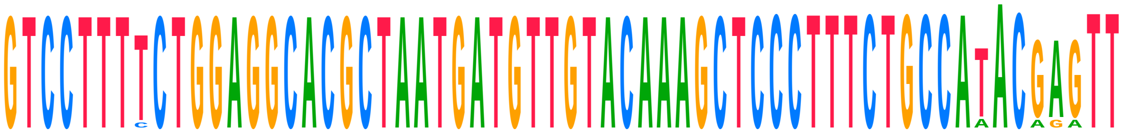

```
Consensus            -----------------------------------------------------A------
Choy_1_ref           GTCCTTTTCTGGAGGCACGCTAATGATGTTGTACAAAGCTCCCTTTCTGCCATACGAGTT  1018
Choy_1               GTCCTTTTCTGGAGGCACGCTAATGATGTTGTACAAAGCTCCCTTTCTGCCATACGAGTT  1018
Choy_10_ref          GTCCTTTTCTGGAGGCACGCTAATGATGTTGTACAAAGCTCCCTTTCTGCCA-ACGAGTT  1011
Choy_10              GTCCTTTCCTGGAGGCACGCTAATGATGTTGTACAAAGCTCCCTTTCTGCCA-ACGAGTT  1011
Choy_11_ref          GTCCTTTTCTGGAGGCACGCTAATGATGTTGTACAAAGCTCCCTTTCTGCCAAACGAGTT  1002
Choy_11              GTCCTTTTCTGGAGGCACGCTAATGATGTTGTACAAAGCTCCCTTTCTGCCAAACGAGTT  1002
Choy_12_ref          GTCCTTTTCTGGAGGCACGCTAATGATGTTGTACAAAGCTCCCTTTCTGCCATACGAGTT  982
```

```
Choy_12            GTCCTTTTCTGGAGGCACGCTAATGATGTTGTACAAAGCTCCCTTTCTGCCATACGAGTT   982
Choy_13_ref        GTCCTTTTCTGGAGGCACGCTAATGATGTTGTACAAAGCTCCCTTTCTGCCATACGAGTT   1020
Choy_13            GTCCTTTTCTGGAGGCACGCTAATGATGTTGTACAAAGCTCCCTTTCTGCCATACGAGTT   1020
Choy_14_ref        GTCCTTTTCTGGAGGCACGCTAATGATGTTGTACAAAGCTCCCTTTCTGCCATACGAGTT   1012
Choy_14            GTCCTTTTCTGGAGGCACGCTAATGATGTTGTACAAAGCTCCCTTTCTGCCATACGAGTT   1012
Choy_5_ref         GTCCTTTTCTGGAGGCACGCTAATGATGTTGTACAAAGCTCCCTTTCTGCCATACGAGTT   1020
Choy_5             GTCCTTTTCTGGAGGCACGCTAATGATGTTGTACAAAGCTCCCTTTCTGCCATACGAGTT   1020
Choy_6_ref         GTCCTTTTCTGGAGGCACGCTAATGATGTTGTACAAAGCTCCCTTTCTGCCATACGAGTT   1020
Choy_6             GTCCTTTTCTGGAGGCACGCTAATGATGTTGTACAAAGCTCCCTTTCTGCCATACGAGTT   1020
Choy_9_ref         GTCCTTTTCTGGAGGCACGCTAATGATGTTGTACAAAGCTCCCTTTCTGCCATACGAGTT   1020
Choy_9             GTCCTTTTCTGGAGGCACGCTAATGATGTTGTACAAAGCTCCCTTTCTGCCATACGAGTT   1020
Pachon_wild_bco2a
  Pachon_1Gross    -----------------------------------------------------A------   926
  Pach_11          -----------------------------------------CCTTTCTGCCATACGAGTT   946
  Pachon...ss_ref  -----------------------------------------------------A------   926
  Pachon2B_ref     -----------------------------------------------------A------   926
  Pachon...OG_ref  -----------------------------------------------------A------   926
  Pach_12_ref      -----------------------------------------------------A------   918
  Pach_12          -----------------------------------------------------A------   918
  Pach_17          -----------------------------------------------------A------   926
  Pachon_F2POG     -----------------------------------------------------A------   926
  Pachon_3Gross    -----------------------------------------------------A------   926
  Pach_15_ref      -----------------------------------------------------A------   926
  Pachon...ro_ref  -----------------------------------------------------A------   926
  Pachon...ss_ref  -----------------------------------------------------A------   926
  Pachon_2Gross    -----------------------------------------------------ACAGATT   932
  Pach_17_ref      -----------------------------------------------------A------   926
  Pachon...ss_ref  -----------------------------------------------------ACAGATT   932
  Pachon_ref_ref   -----------------------------------------------------A------   926
  Pach_8           -----------------------------------------------------A------   926
  Pach_8_ref       -----------------------------------------------------A------   926
  Pachon...OG_ref  -----------------------------------------------------A------   926
  Pachon_6Boro     -----------------------------------------------------A------   926
  Pachon_ref       -----------------------------------------------------A------   926
  Pach_3_ref       -----------------------------------------------------A------   926
  Pach_9           -----------------------------------------------------A------   926
  Pachon_G2POG     -----------------------------------------------------A------   926
  Pachon...OG_ref  -----------------------------------------------------A------   926
  Pach_9_ref       -----------------------------------------------------A------   926
  Pachon2B         -----------------------------------------------------A------   926
  Pachon...OG_ref  -----------------------------------------------------A------   926
  Pachon_E2POG     -----------------------------------------------------A------   926
  Pach_14          -----------------------------------------------------A------   926
  Pach_3           -----------------------------------------------------A------   926
  Pachon_H2POG     -----------------------------------------------------A------   926
  Pach_7_ref       ------------------------------------------------------------   911
  Pach_15          -----------------------------------------------------A------   926
  Pach_14_ref      -----------------------------------------------------A------   926
  Pach_11_ref      -----------------------------------------CCTTTCTGCCATACGAGTT   946
  Pach_7           ------------------------------------------------------------   911
```

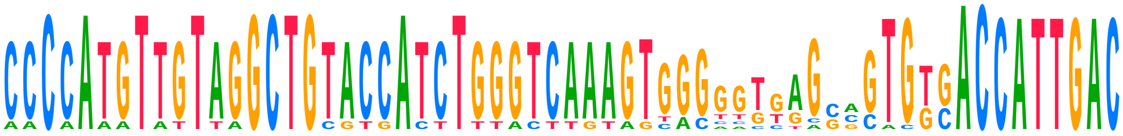

```
Consensus          -------------------------------------NNNNNNGNNGTGTGAC-------

Choy_1_ref         CCCCATGTTGTAGGCTGTACCATCTGGGTCAAAGTGGGGGTGAGCAGTGGCACCATTGAC   1078
Choy_1             CCCCATGTTGTAGGCTGTACCATCTGGGTCAAAGTGGGGGTGAGCAGTGGCACCATTGAC   1078
Choy_10_ref        CCCCATGTTGTAGGCTGTACCATCTGGGTCAAAGTGGGGGTGAGCAGTGGCACCATTGAC   1071
Choy_10            CCCCATGTTGTAGGCTGTACCATCTGGGTCAAAGTGGGGGTGAGCAGTGGCACCATTGAC   1071
```

| | | |
|---|---|---|
| Choy_11_ref | CCCCATGTTGTAGGCTGTACCATCTGGGTCAAAGTGGGGGTGAGCAGTGGCACCATTGAC | 1062 |
| Choy_11 | CCCCATGTTGTAGGCTGTACCATCTGGGTCAAAGTGGGGGTGAGCAGTGGCACCATTGAC | 1062 |
| Choy_12_ref | CCCCATGTTGTAGGCTGTACCATCTGGGTCAAAGTGGGGGTGAGCAGTGGCACCATTGAC | 1042 |
| Choy_12 | CCCCATGTTGTAGGCTGTACCATCTGGGTCAAAGTGGGGGTGAGCAGTGGCACCATTGAC | 1042 |
| Choy_13_ref | CCCCATGTTGTAGGCTGTACCATCTGGGTCAAAGTGGGGGTGAGCAGTGGCACCATTGAC | 1080 |
| Choy_13 | CCCCATGTTGTAGGCTGTACCATCTGGGTCAAAGTGGGGGTGAGCAGTGGCACCATTGAC | 1080 |
| Choy_14_ref | CCCCATGTTGTAGGCTGTACCATCTGGGTCAAAGTGGGGGTGAGCAGTGGCACCATTGAC | 1072 |
| Choy_14 | CCCCATGTTGTAGGCTGTACCATCTGGGTCAAAGTGGGGGTGAGCAGTGGCACCATTGAC | 1072 |
| Choy_5_ref | CCCCATGTTGTAGGCTGTACCATCTGGGTCAAAGTGGGGGTGAGCAGTGGCACCATTGAC | 1080 |
| Choy_5 | CCCCATGTTGTAGGCTGTACCATCTGGGTCAAAGTGGGGGTGAGCAGTGGCACCATTGAC | 1080 |
| Choy_6_ref | CCCCATGTTGTAGGCTGTACCATCTGGGTCAAAGTGGGGGTGAGCAGTGGCACCATTGAC | 1080 |
| Choy_6 | CCCCATGTTGTAGGCTGTACCATCTGGGTCAAAGTGGGGGTGAGCAGTGGCACCATTGAC | 1080 |
| Choy_9_ref | CCCCATGTTGTAGGCTGTACCATCTGGGTCAAAGTGGGGGTGAGCAGTGGCACCATTGAC | 1080 |
| Choy_9 | CCCCATGTTGTAGGCTGTACCATCTGGGTCAAAGTGGGGGTGAGCAGTGGCACCATTGAC | 1080 |
| Pachon_wild_bco2a | | |
| Pachon_1Gross | -------------------------------------------GGCGTGTGAC------- | 936 |
| Pach_11 | CCCCATGTTGTAGGCTGTACCATCTGGGTCAAAGTGGGGGTGAGGCCTGTGAC------- | 999 |
| Pachon…ss_ref | -------------------------------------------GGCGTGTGAC------- | 936 |
| Pachon2B_ref | ----------------------------------------GCTGGCGTGTGAC------- | 939 |
| Pachon…OG_ref | ------------------------------------------GGGCGTGTGAC------- | 937 |
| Pach_12_ref | ---------------------------------------------GGTGTGAC------- | 926 |
| Pach_12 | ---------------------------------------------GGTGTGAC------- | 926 |
| Pach_17 | -------------------------------------------GCCGTGTGAC------- | 936 |
| Pachon_F2POG | -------------------------------------GCACTGGGCGTGTGAC------- | 942 |
| Pachon_3Gross | ----------------------------------------GCACGCGTGTGAC------- | 939 |
| Pach_15_ref | ---------------------------------------------GGTGTGAC------- | 934 |
| Pachon…ro_ref | --------------------------------------GCGTGGGCGTGTGAC------- | 941 |
| Pachon…ss_ref | ----------------------------------------GCACGCGTGTGAC------- | 939 |
| Pachon_2Gross | AACAAAATATTTAGCTGCGTGACTTTTACTTGTATTACTTTTAATAGTGTGAC------- | 985 |
| Pach_17_ref | -------------------------------------------GCCGTGTGAC------- | 936 |
| Pachon…ss_ref | AACAAAATATTTAGCTGCGTGACTTTTACTTGTATTACTTTTAATAGTGTGAC------- | 985 |
| Pachon_ref_ref | ----------------------------------GCACAGTTAGAGCTGTGAC------- | 945 |
| Pach_8 | --------------------------------------------------GAC------- | 929 |
| Pach_8_ref | --------------------------------------------------GAC------- | 929 |
| Pachon…OG_ref | -------------------------------------GTTGTGGGCGTGTGAC------- | 942 |
| Pachon_6Boro | --------------------------------------GCGTGGGCGTGTGAC------- | 941 |
| Pachon_ref | ----------------------------------GCACAGTTAGAGCTGTGAC------- | 945 |
| Pach_3_ref | ---------------------------------------------GCTGTGAC------- | 934 |
| Pach_9 | -------------------------------------------GCACTGTGAC------- | 936 |
| Pachon_G2POG | ------------------------------------------GGGCGTGTGAC------- | 937 |
| Pachon…OG_ref | ---------------------------------------------GGTGTGAC------- | 934 |
| Pach_9_ref | -------------------------------------------GCACTGTGAC------- | 936 |
| Pachon2B | ----------------------------------------GCTGGCGTGTGAC------- | 939 |
| Pachon…OG_ref | -------------------------------------GCACTGGGCGTGTGAC------- | 942 |
| Pachon_E2POG | -------------------------------------GTTGTGGGCGTGTGAC------- | 942 |
| Pach_14 | ---------------------------------------------GCACTGAC------- | 934 |
| Pach_3 | ---------------------------------------------GCTGTGAC------- | 934 |
| Pachon_H2POG | ---------------------------------------------GGTGTGAC------- | 934 |
| Pach_7_ref | ------------------------------------------------------------ | 911 |
| Pach_15 | ---------------------------------------------GGTGTGAC------- | 934 |
| Pach_14_ref | ---------------------------------------------GCACTGAC------- | 934 |
| Pach_11_ref | CCCCATGTTGTAGGCTGTACCATCTGGGTCAAAGTGGGGGTGAGGCCTGTGAC------- | 999 |
| Pach_7 | ------------------------------------------------------------ | 911 |

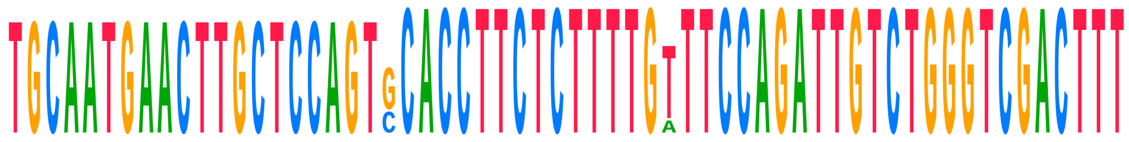

| | | |
|---|---|---|
| **Consensus** | **--------------------GCACCTTCTCTTTTGTTTCCAGATTGTCTGGGTCGACTTT** |  |
| Choy_1_ref | TGCAATGAACTTGCTCCAGTCCACCTTCTCTTTTGATTCCAGATTGTCTGGGTCGACTTT | 1138 |

| | | |
|---|---|---|
| Choy_1 | TGCAATGAACTTGCTCCAGTCCACCTTCTCTTTTGTTTCCAGATTGTCTGGGTCGACTTT | 1138 |
| Choy_10_ref | TGCAATGAACTTGCTCCAGTCCACCTTCTCTTTTGTTTCCAGATTGTCTGGGTCGACTTT | 1131 |
| Choy_10 | TGCAATGAACTTGCTCCAGTCCACCTTCTCTTTTGTTTCCAGATTGTCTGGGTCGACTTT | 1131 |
| Choy_11_ref | TGCAATGAACTTGCTCCAGTCCACCTTCTCTTTTGTTTCCAGATTGTCTGGGTCGACTTT | 1122 |
| Choy_11 | TGCAATGAACTTGCTCCAGTCCACCTTCTCTTTTGTTTCCAGATTGTCTGGGTCGACTTT | 1122 |
| Choy_12_ref | TGCAATGAACTTGCTCCAGTCCACCTTCTCTTTTGTTTCCAGATTGTCTGGGTCGACTTT | 1102 |
| Choy_12 | TGCAATGAACTTGCTCCAGTCCACCTTCTCTTTTGTTTCCAGATTGTCTGGGTCGACTTT | 1102 |
| Choy_13_ref | TGCAATGAACTTGCTCCAGTCCACCTTCTCTTTTGATTCCAGATTGTCTGGGTCGACTTT | 1140 |
| Choy_13 | TGCAATGAACTTGCTCCAGTCCACCTTCTCTTTTGTTTCCAGATTGTCTGGGTCGACTTT | 1140 |
| Choy_14_ref | TGCAATGAACTTGCTCCAGTCCACCTTCTCTTTTGATTCCAGATTGTCTGGGTCGACTTT | 1132 |
| Choy_14 | TGCAATGAACTTGCTCCAGTCCACCTTCTCTTTTGATTCCAGATTGTCTGGGTCGACTTT | 1132 |
| Choy_5_ref | TGCAATGAACTTGCTCCAGTCCACCTTCTCTTTTGATTCCAGATTGTCTGGGTCGACTTT | 1140 |
| Choy_5 | TGCAATGAACTTGCTCCAGTCCACCTTCTCTTTTGATTCCAGATTGTCTGGGTCGACTTT | 1140 |
| Choy_6_ref | TGCAATGAACTTGCTCCAGTCCACCTTCTCTTTTGATTCCAGATTGTCTGGGTCGACTTT | 1140 |
| Choy_6 | TGCAATGAACTTGCTCCAGTCCACCTTCTCTTTTGATTCCAGATTGTCTGGGTCGACTTT | 1140 |
| Choy_9_ref | TGCAATGAACTTGCTCCAGTCCACCTTCTCTTTTGTTTCCAGATTGTCTGGGTCGACTTT | 1140 |
| Choy_9 | TGCAATGAACTTGCTCCAGTCCACCTTCTCTTTTGTTTCCAGATTGTCTGGGTCGACTTT | 1140 |
| Pachon_wild_bco2a | | |
| Pachon_1Gross | --------------------GCACCTTCTCTTTTGTTTCCAGATTGTCTGGGTCGACTTT | 976 |
| Pach_11 | --------------------GCACCTTCTCTTTTGTTTCCAGATTGTCTGGGTCGACTTT | 1039 |
| Pachon...ss_ref | --------------------GCACCTTCTCTTTTGTTTCCAGATTGTCTGGGTCGACTTT | 976 |
| Pachon2B_ref | --------------------GCACCTTCTCTTTTGTTTCCAGATTGTCTGGGTCGACTTT | 979 |
| Pachon...OG_ref | --------------------GCACCTTCTCTTTTGTTTCCAGATTGTCTGGGTCGACTTT | 977 |
| Pach_12_ref | --------------------GCACCTTCTCTTTTGTTTCCAGATTGTCTGGGTCGACTTT | 966 |
| Pach_12 | --------------------GCACCTTCTCTTTTGTTTCCAGATTGTCTGGGTCGACTTT | 966 |
| Pach_17 | --------------------GCACCTTCTCTTTTGTTTCCAGATTGTCTGGGTCGACTTT | 976 |
| Pachon_F2POG | --------------------GCACCTTCTCTTTTGTTTCCAGATTGTCTGGGTCGACTTT | 982 |
| Pachon_3Gross | --------------------GCACCTTCTCTTTTGTTTCCAGATTGTCTGGGTCGACTTT | 979 |
| Pach_15_ref | --------------------GCACCTTCTCTTTTGTTTCCAGATTGTCTGGGTCGACTTT | 974 |
| Pachon...ro_ref | --------------------GCACCTTCTCTTTTGTTTCCAGATTGTCTGGGTCGACTTT | 981 |
| Pachon...ss_ref | --------------------GCACCTTCTCTTTTGTTTCCAGATTGTCTGGGTCGACTTT | 979 |
| Pachon_2Gross | --------------------GCACCTTCTCTTTTGTTTCCAGATTGTCTGGGTCGACTTT | 1025 |
| Pach_17_ref | --------------------GCACCTTCTCTTTTGTTTCCAGATTGTCTGGGTCGACTTT | 976 |
| Pachon...ss_ref | --------------------GCACCTTCTCTTTTGTTTCCAGATTGTCTGGGTCGACTTT | 1025 |
| Pachon_ref_ref | --------------------GCACCTTCTCTTTTGTTTCCAGATTGTCTGGGTCGACTTT | 985 |
| Pach_8 | --------------------GCACCTTCTCTTTTGTTTCCAGATTGTCTGGGTCGACTTT | 969 |
| Pach_8_ref | --------------------GCACCTTCTCTTTTGTTTCCAGATTGTCTGGGTCGACTTT | 969 |
| Pachon...OG_ref | --------------------GCACCTTCTCTTTTGTTTCCAGATTGTCTGGGTCGACTTT | 982 |
| Pachon_6Boro | --------------------GCACCTTCTCTTTTGTTTCCAGATTGTCTGGGTCGACTTT | 981 |
| Pachon_ref | --------------------GCACCTTCTCTTTTGTTTCCAGATTGTCTGGGTCGACTTT | 985 |
| Pach_3_ref | --------------------GCACCTTCTCTTTTGTTTCCAGATTGTCTGGGTCGACTTT | 974 |
| Pach_9 | --------------------GCACCTTCTCTTTTGTTTCCAGATTGTCTGGGTCGACTTT | 976 |
| Pachon_G2POG | --------------------GCACCTTCTCTTTTGTTTCCAGATTGTCTGGGTCGACTTT | 977 |
| Pachon...OG_ref | --------------------GCACCTTCTCTTTTGTTTCCAGATTGTCTGGGTCGACTTT | 974 |
| Pach_9_ref | --------------------GCACCTTCTCTTTTGTTTCCAGATTGTCTGGGTCGACTTT | 976 |
| Pachon2B | --------------------GCACCTTCTCTTTTGTTTCCAGATTGTCTGGGTCGACTTT | 979 |
| Pachon...OG_ref | --------------------GCACCTTCTCTTTTGTTTCCAGATTGTCTGGGTCGACTTT | 982 |
| Pachon_E2POG | --------------------GCACCTTCTCTTTTGTTTCCAGATTGTCTGGGTCGACTTT | 982 |
| Pach_14 | --------------------GCACCTTCTCTTTTGTTTCCAGATTGTCTGGGTCGACTTT | 974 |
| Pach_3 | --------------------GCACCTTCTCTTTTGTTTCCAGATTGTCTGGGTCGACTTT | 974 |
| Pachon_H2POG | --------------------GCACCTTCTCTTTTGTTTCCAGATTGTCTGGGTCGACTTT | 974 |
| Pach_7_ref | ---------------------CACCTTCTCTTTTGTTTCCAGATTGTCTGGGTCGACTTT | 950 |
| Pach_15 | --------------------GCACCTTCTCTTTTGTTTCCAGATTGTCTGGGTCGACTTT | 974 |
| Pach_14_ref | --------------------GCACCTTCTCTTTTGTTTCCAGATTGTCTGGGTCGACTTT | 974 |
| Pach_11_ref | --------------------GCACCTTCTCTTTTGTTTCCAGATTGTCTGGGTCGACTTT | 1039 |
| Pach_7 | ---------------------CACCTTCTCTTTTGTTTCCAGATTGTCTGGGTCGACTTT | 950 |

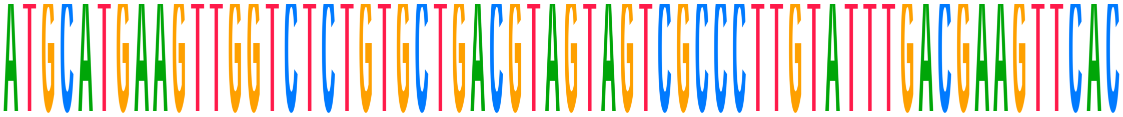

| | | |
|---|---|---|
| **Consensus** | **ATGCATGAAGTTGGTCTCTGTGCTGACGTAGTAGTCGCCCTTGTATTTGACGAAGTTCAC** | |
| Choy_1_ref | ATGCATGAAGTTGGTCTCTGTGCTGACGTAGTAGTCGCCCTTGTATTTGACGAAGTTCAC | 1198 |
| Choy_1 | ATGCATGAAGTTGGTCTCTGTGCTGACGTAGTAGTCGCCCTTGTATTTGACGAAGTTCAC | 1198 |
| Choy_10_ref | ATGCATGAAGTTGGTCTCTGTGCTGACGTAGTAGTCGCCCTTGTATTTGACGAAGTTCAC | 1191 |
| Choy_10 | ATGCATGAAGTTGGTCTCTGTGCTGACGTAGTAGTCGCCCTTGTATTTGACGAAGTTCAC | 1191 |
| Choy_11_ref | ATGCATGAAGTTGGTCTCTGTGCTGACGTAGTAGTCGCCCTTGTATTTGACGAAGTTCAC | 1182 |
| Choy_11 | ATGCATGAAGTTGGTCTCTGTGCTGACGTAGTAGTCGCCCTTGTATTTGACGAAGTTCAC | 1182 |
| Choy_12_ref | ATGCATGAAGTTGGTCTCTGTGCTGACGTAGTAGTCGCCCTTGTATTTGACGAAGTTCAC | 1162 |
| Choy_12 | ATGCATGAAGTTGGTCTCTGTGCTGACGTAGTAGTCGCCCTTGTATTTGACGAAGTTCAC | 1162 |
| Choy_13_ref | ATGCATGAAGTTGGTCTCTGTGCTGACGTAGTAGTCGCCCTTGTATTTGACGAAGTTCAC | 1200 |
| Choy_13 | ATGCATGAAGTTGGTCTCTGTGCTGACGTAGTAGTCGCCCTTGTATTTGACGAAGTTCAC | 1200 |
| Choy_14_ref | ATGCATGAAGTTGGTCTCTGTGCTGACGTAGTAGTCGCCCTTGTATTTGACGAAGTTCAC | 1192 |
| Choy_14 | ATGCATGAAGTTGGTCTCTGTGCTGACGTAGTAGTCGCCCTTGTATTTGACGAAGTTCAC | 1192 |
| Choy_5_ref | ATGCATGAAGTTGGTCTCTGTGCTGACGTAGTAGTCGCCCTTGTATTTGACGAAGTTCAC | 1200 |
| Choy_5 | ATGCATGAAGTTGGTCTCTGTGCTGACGTAGTAGTCGCCCTTGTATTTGACGAAGTTCAC | 1200 |
| Choy_6_ref | ATGCATGAAGTTGGTCTCTGTGCTGACGTAGTAGTCGCCCTTGTATTTGACGAAGTTCAC | 1200 |
| Choy_6 | ATGCATGAAGTTGGTCTCTGTGCTGACGTAGTAGTCGCCCTTGTATTTGACGAAGTTCAC | 1200 |
| Choy_9_ref | ATGCATGAAGTTGGTCTCTGTGCTGACGTAGTAGTCGCCCTTGTATTTGACGAAGTTCAC | 1200 |
| Choy_9 | ATGCATGAAGTTGGTCTCTGTGCTGACGTAGTAGTCGCCCTTGTATTTGACGAAGTTCAC | 1200 |
| Pachon_wild_bco2a | | |
| Pachon_1Gross | ATGCATGAAGTTGGTCTCTGTGCTGACGTAGTAGTCGCCCTTGTATTTGACGAAGTTCAC | 1036 |
| Pach_11 | ATGCATGAAGTTGGTCTCTGTGCTGACGTAGTAGTCGCCCTTGTATTTGACGAAGTTCAC | 1099 |
| Pachon...ss_ref | ATGCATGAAGTTGGTCTCTGTGCTGACGTAGTAGTCGCCCTTGTATTTGACGAAGTTCAC | 1036 |
| Pachon2B_ref | ATGCATGAAGTTGGTCTCTGTGCTGACGTAGTAGTCGCCCTTGTATTTGACGAAGTTCAC | 1039 |
| Pachon...OG_ref | ATGCATGAAGTTGGTCTCTGTGCTGACGTAGTAGTCGCCCTTGTATTTGACGAAGTTCAC | 1037 |
| Pach_12_ref | ATGCATGAAGTTGGTCTCTGTGCTGACGTAGTAGTCGCCCTTGTATTTGACGAAGTTCAC | 1026 |
| Pach_12 | ATGCATGAAGTTGGTCTCTGTGCTGACGTAGTAGTCGCCCTTGTATTTGACGAAGTTCAC | 1026 |
| Pach_17 | ATGCATGAAGTTGGTCTCTGTGCTGACGTAGTAGTCGCCCTTGTATTTGACGAAGTTCAC | 1036 |
| Pachon_F2POG | ATGCATGAAGTTGGTCTCTGTGCTGACGTAGTAGTCGCCCTTGTATTTGACGAAGTTCAC | 1042 |
| Pachon_3Gross | ATGCATGAAGTTGGTCTCTGTGCTGACGTAGTAGTCGCCCTTGTATTTGACGAAGTTCAC | 1039 |
| Pach_15_ref | ATGCATGAAGTTGGTCTCTGTGCTGACGTAGTAGTCGCCCTTGTATTTGACGAAGTTCAC | 1034 |
| Pachon...ro_ref | ATGCATGAAGTTGGTCTCTGTGCTGACGTAGTAGTCGCCCTTGTATTTGACGAAGTTCAC | 1041 |
| Pachon...ss_ref | ATGCATGAAGTTGGTCTCTGTGCTGACGTAGTAGTCGCCCTTGTATTTGACGAAGTTCAC | 1039 |
| Pachon_2Gross | ATGCATGAAGTTGGTCTCTGTGCTGACGTAGTAGTCGCCCTTGTATTTGACGAAGTTCAC | 1085 |
| Pach_17_ref | ATGCATGAAGTTGGTCTCTGTGCTGACGTAGTAGTCGCCCTTGTATTTGACGAAGTTCAC | 1036 |
| Pachon...ss_ref | ATGCATGAAGTTGGTCTCTGTGCTGACGTAGTAGTCGCCCTTGTATTTGACGAAGTTCAC | 1085 |
| Pachon_ref_ref | ATGCATGAAGTTGGTCTCTGTGCTGACGTAGTAGTCGCCCTTGTATTTGACGAAGTTCAC | 1045 |
| Pach_8 | ATGCATGAAGTTGGTCTCTGTGCTGACGTAGTAGTCGCCCTTGTATTTGACGAAGTTCAC | 1029 |
| Pach_8_ref | ATGCATGAAGTTGGTCTCTGTGCTGACGTAGTAGTCGCCCTTGTATTTGACGAAGTTCAC | 1029 |
| Pachon...OG_ref | ATGCATGAAGTTGGTCTCTGTGCTGACGTAGTAGTCGCCCTTGTATTTGACGAAGTTCAC | 1042 |
| Pachon_6Boro | ATGCATGAAGTTGGTCTCTGTGCTGACGTAGTAGTCGCCCTTGTATTTGACGAAGTTCAC | 1041 |
| Pachon_ref | ATGCATGAAGTTGGTCTCTGTGCTGACGTAGTAGTCGCCCTTGTATTTGACGAAGTTCAC | 1045 |
| Pach_3_ref | ATGCATGAAGTTGGTCTCTGTGCTGACGTAGTAGTCGCCCTTGTATTTGACGAAGTTCAC | 1034 |
| Pach_9 | ATGCATGAAGTTGGTCTCTGTGCTGACGTAGTAGTCGCCCTTGTATTTGACGAAGTTCAC | 1036 |
| Pachon_G2POG | ATGCATGAAGTTGGTCTCTGTGCTGACGTAGTAGTCGCCCTTGTATTTGACGAAGTTCAC | 1037 |
| Pachon...OG_ref | ATGCATGAAGTTGGTCTCTGTGCTGACGTAGTAGTCGCCCTTGTATTTGACGAAGTTCAC | 1034 |
| Pach_9_ref | ATGCATGAAGTTGGTCTCTGTGCTGACGTAGTAGTCGCCCTTGTATTTGACGAAGTTCAC | 1036 |
| Pachon2B | ATGCATGAAGTTGGTCTCTGTGCTGACGTAGTAGTCGCCCTTGTATTTGACGAAGTTCAC | 1039 |
| Pachon...OG_ref | ATGCATGAAGTTGGTCTCTGTGCTGACGTAGTAGTCGCCCTTGTATTTGACGAAGTTCAC | 1042 |
| Pachon_E2POG | ATGCATGAAGTTGGTCTCTGTGCTGACGTAGTAGTCGCCCTTGTATTTGACGAAGTTCAC | 1042 |
| Pach_14 | ATGCATGAAGTTGGTCTCTGTGCTGACGTAGTAGTCGCCCTTGTATTTGACGAAGTTCAC | 1034 |
| Pach_3 | ATGCATGAAGTTGGTCTCTGTGCTGACGTAGTAGTCGCCCTTGTATTTGACGAAGTTCAC | 1034 |
| Pachon_H2POG | ATGCATGAAGTTGGTCTCTGTGCTGACGTAGTAGTCGCCCTTGTATTTGACGAAGTTCAC | 1034 |
| Pach_7_ref | ATGCATGAAGTTGGTCTCTGTGCTGACGT------------------------------C | 980 |
| Pach_15 | ATGCATGAAGTTGGTCTCTGTGCTGACGTAGTAGTCGCCCTTGTATTTGACGAAGTTCAC | 1034 |
| Pach_14_ref | ATGCATGAAGTTGGTCTCTGTGCTGACGTAGTAGTCGCCCTTGTATTTGACGAAGTTCAC | 1034 |
| Pach_11_ref | ATGCATGAAGTTGGTCTCTGTGCTGACGTAGTAGTCGCCCTTGTATTTGACGAAGTTCAC | 1099 |
| Pach_7 | ATGCATGAAGTTGGTCTCTGTGCTGACGT------------------------------C | 980 |

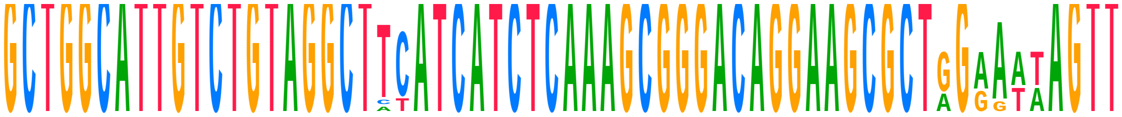

| | | |
|---|---|---|
| **Consensus** | **GCTGGCATTGTCTGTAGGCTTC------------------------------NANNAGTT** |  |
| Choy_1_ref | GCTGGCATTGTCTGTAGGCTTCATCATCTCAAAGCGGGACAGGAAGCGCTGGAATAAGTT | 1258 |
| Choy_1 | GCTGGCATTGTCTGTAGGCTTCATCATCTCAAAGCGGGACAGGAAGCGCTGGAATAAGTT | 1258 |
| Choy_10_ref | GCTGGCATTGTCTGTAGGCTTCATCATCTCAAAGCGGGACAGGAAGCGCTGGAATAAGTT | 1251 |
| Choy_10 | GCTGGCATTGTCTGTAGGCTTCATCATCTCAAAGCGGGACAGGAAGCGCTGGAATAAGTT | 1251 |
| Choy_11_ref | GCTGGCATTGTCTGTAGGCTTCATCATCTCAAAGCGGGACAGGAAGCGCTGGAATAAGTT | 1242 |
| Choy_11 | GCTGGCATTGTCTGTAGGCTTCATCATCTCAAAGCGGGACAGGAAGCGCTGGAATAAGTT | 1242 |
| Choy_12_ref | GCTGGCATTGTCTGTAGGCTTCATCATCTCAAAGCGGGACAGGAAGCGCTGGAATAAGTT | 1222 |
| Choy_12 | GCTGGCATTGTCTGTAGGCTTCATCATCTCAAAGCGGGACAGGAAGCGCTGGAATAAGTT | 1222 |
| Choy_13_ref | GCTGGCATTGTCTGTAGGCTTCATCATCTCAAAGCGGGACAGGAAGCGCTGGAATAAGTT | 1260 |
| Choy_13 | GCTGGCATTGTCTGTAGGCTTCATCATCTCAAAGCGGGACAGGAAGCGCTGGAATAAGTT | 1260 |
| Choy_14_ref | GCTGGCATTGTCTGTAGGCTTCATCATCTCAAAGCGGGACAGGAAGCGCTGGAATAAGTT | 1252 |
| Choy_14 | GCTGGCATTGTCTGTAGGCTTCATCATCTCAAAGCGGGACAGGAAGCGCTGGAATAAGTT | 1252 |
| Choy_5_ref | GCTGGCATTGTCTGTAGGCTTCATCATCTCAAAGCGGGACAGGAAGCGCTGGAATAAGTT | 1260 |
| Choy_5 | GCTGGCATTGTCTGTAGGCTTCATCATCTCAAAGCGGGACAGGAAGCGCTGGAATAAGTT | 1260 |
| Choy_6_ref | GCTGGCATTGTCTGTAGGCTTCATCATCTCAAAGCGGGACAGGAAGCGCTGGAATAAGTT | 1260 |
| Choy_6 | GCTGGCATTGTCTGTAGGCTTCATCATCTCAAAGCGGGACAGGAAGCGCTGGAATAAGTT | 1260 |
| Choy_9_ref | GCTGGCATTGTCTGTAGGCTTCATCATCTCAAAGCGGGACAGGAAGCGCTGGAATAAGTT | 1260 |
| Choy_9 | GCTGGCATTGTCTGTAGGCTTCATCATCTCAAAGCGGGACAGGAAGCGCTGGAATAAGTT | 1260 |
| Pachon_wild_bco2a | | |
| Pachon_1Gross | GCTGGCATTGTCTGTAGGCT----------------------------------ATAGTT | 1062 |
| Pach_11 | GCTGGCATTGTCTGTAGGCTTCA---------------------------AGAGATAGTT | 1132 |
| Pachon…ss_ref | GCTGGCATTGTCTGTAGGCT----------------------------------ATAGTT | 1062 |
| Pachon2B_ref | GCTGGCATTGTCTGTAGGCTTC--------------------------------ATAGTT | 1067 |
| Pachon…OG_ref | GCTGGCATTGTCTGTAGGCTT------------------------------------GTT | 1061 |
| Pach_12_ref | GCTGGCATTGTCTGTAGGCT----------------------------------ATAGTT | 1052 |
| Pach_12 | GCTGGCATTGTCTGTAGGCT----------------------------------ATAGTT | 1052 |
| Pach_17 | GCTGGCATTGTCTGTAGGCTTCA---------------------------AGGAATAGTT | 1069 |
| Pachon_F2POG | GCTGGCATTGTCTGTAGGCT-------------------------------------GTT | 1065 |
| Pachon_3Gross | GCTGGCATTGTCTGTAGGCT-------------------------------------GTT | 1062 |
| Pach_15_ref | GCTGGCATTGTCTGTAGGCTTCA---------------------------AGAAATAGTT | 1067 |
| Pachon…ro_ref | GCTGGCATTGTCTGTAGGCT--------------------------------GAATAGTT | 1069 |
| Pachon…ss_ref | GCTGGCATTGTCTGTAGGCT-------------------------------------GTT | 1062 |
| Pachon_2Gross | GCTGGCATTGTCTGTAGGCTTC--------------------------------ATAGTT | 1113 |
| Pach_17_ref | GCTGGCATTGTCTGTAGGCTTCA---------------------------AGGAATAGTT | 1069 |
| Pachon…ss_ref | GCTGGCATTGTCTGTAGGCTTC--------------------------------ATAGTT | 1113 |
| Pachon_ref_ref | GCTGGCATTGTCTGTAGGCTTC------------------------------GAATAGTT | 1075 |
| Pach_8 | GCTGGCATTGTCTGTAGGCTTT------------------------------GAATAGTT | 1059 |
| Pach_8_ref | GCTGGCATTGTCTGTAGGCTTT------------------------------GAATAGTT | 1059 |
| Pachon…OG_ref | GCTGGCATTGTCTGTAGGCTAT-----------------------------------GTT | 1067 |
| Pachon_6Boro | GCTGGCATTGTCTGTAGGCT--------------------------------GAATAGTT | 1069 |
| Pachon_ref | GCTGGCATTGTCTGTAGGCTTC------------------------------GAATAGTT | 1075 |
| Pach_3_ref | GCTGGCATTGTCTGTAGGCTCT------------------------------GAATAGTT | 1064 |
| Pach_9 | GCTGGCATTGTCTGTAGGCT---------------------------------GATAGTT | 1063 |
| Pachon_G2POG | GCTGGCATTGTCTGTAGGCTT------------------------------------GTT | 1061 |
| Pachon…OG_ref | GCTGGCATTGTCTGTAGGCT-------------------------------------GTT | 1057 |
| Pach_9_ref | GCTGGCATTGTCTGTAGGCT---------------------------------GATAGTT | 1063 |
| Pachon2B | GCTGGCATTGTCTGTAGGCTTC--------------------------------ATAGTT | 1067 |
| Pachon…OG_ref | GCTGGCATTGTCTGTAGGCT-------------------------------------GTT | 1065 |
| Pachon_E2POG | GCTGGCATTGTCTGTAGGCTAT-----------------------------------GTT | 1067 |
| Pach_14 | GCTGGCATTGTCTGTAGGCT--A---------------------------AGGAATAGTT | 1065 |
| Pach_3 | GCTGGCATTGTCTGTAGGCTCT------------------------------GAATAGTT | 1064 |
| Pachon_H2POG | GCTGGCATTGTCTGTAGGCT-------------------------------------GTT | 1057 |
| Pach_7_ref | GCTGGCATTGTCTGTAGGCT---------------------------------------- | 1000 |
| Pach_15 | GCTGGCATTGTCTGTAGGCTTCA---------------------------AGAAATAGTT | 1067 |
| Pach_14_ref | GCTGGCATTGTCTGTAGGCT--A---------------------------AGGAATAGTT | 1065 |
| Pach_11_ref | GCTGGCATTGTCTGTAGGCTTCA---------------------------AGAGATAGTT | 1132 |
| Pach_7 | GCTGGCATTGTCTGTAGGCT---------------------------------------- | 1000 |

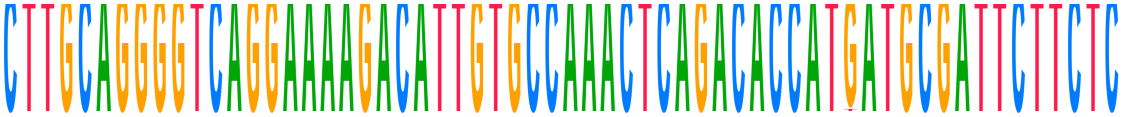

| | | |
|---|---|---|
| **Consensus** | **CTTGCAGGGGTCAGGAAAAGACATTGTGCCAAACTCAGACACCATGATGCGATTCTTCTC** | |
| Choy_1_ref | CTTGCAGGGGTCAGGAAAAGACATTGTGCCAAACTCAGACACCATGATGCGATTCTTCTC | 1318 |
| Choy_1 | CTTGCAGGGGTCAGGAAAAGACATTGTGCCAAACTCAGACACCATGATGCGATTCTTCTC | 1318 |
| Choy_10_ref | CTTGCAGGGGTCAGGAAAAGACATTGTGCCAAACTCAGACACCATGATGCGATTCTTCTC | 1311 |
| Choy_10 | CTTGCAGGGGTCAGGAAAAGACATTGTGCCAAACTCAGACACCATGATGCGATTCTTCTC | 1311 |
| Choy_11_ref | CTTGCAGGGGTCAGGAAAAGACATTGTGCCAAACTCAGACACCATGATGCGATTCTTCTC | 1302 |
| Choy_11 | CTTGCAGGGGTCAGGAAAAGACATTGTGCCAAACTCAGACACCATGATGCGATTCTTCTC | 1302 |
| Choy_12_ref | CTTGCAGGGGTCAGGAAAAGACATTGTGCCAAACTCAGACACCATGATGCGATTCTTCTC | 1282 |
| Choy_12 | CTTGCAGGGGTCAGGAAAAGACATTGTGCCAAACTCAGACACCATTATGCGATTCTTCTC | 1282 |
| Choy_13_ref | CTTGCAGGGGTCAGGAAAAGACATTGTGCCAAACTCAGACACCATGATGCGATTCTTCTC | 1320 |
| Choy_13 | CTTGCAGGGGTCAGGAAAAGACATTGTGCCAAACTCAGACACCATGATGCGATTCTTCTC | 1320 |
| Choy_14_ref | CTTGCAGGGGTCAGGAAAAGACATTGTGCCAAACTCAGACACCATGATGCGATTCTTCTC | 1312 |
| Choy_14 | CTTGCAGGGGTCAGGAAAAGACATTGTGCCAAACTCAGACACCATGATGCGATTCTTCTC | 1312 |
| Choy_5_ref | CTTGCAGGGGTCAGGAAAAGACATTGTGCCAAACTCAGACACCATGATGCGATTCTTCTC | 1320 |
| Choy_5 | CTTGCAGGGGTCAGGAAAAGACATTGTGCCAAACTCAGACACCATGATGCGATTCTTCTC | 1320 |
| Choy_6_ref | CTTGCAGGGGTCAGGAAAAGACATTGTGCCAAACTCAGACACCATGATGCGATTCTTCTC | 1320 |
| Choy_6 | CTTGCAGGGGTCAGGAAAAGACATTGTGCCAAACTCAGACACCATGATGCGATTCTTCTC | 1320 |
| Choy_9_ref | CTTGCAGGGGTCAGGAAAAGACATTGTGCCAAACTCAGACACCATGATGCGATTCTTCTC | 1320 |
| Choy_9 | CTTGCAGGGGTCAGGAAAAGACATTGTGCCAAACTCAGACACCATGATGCGATTCTTCTC | 1320 |
| Pachon_wild_bco2a | | |
| Pachon_1Gross | CTTGCAGGGGTCAGGAAAAGACATTGTGCCAAACTCAGACACCATGATGCGATTCTTCTC | 1122 |
| Pach_11 | CTTGCAGGGGTCAGGAAAAGACATTGTGCCAAACTCAGACACCATGATGCGATTCTTCTC | 1192 |
| Pachon...ss_ref | CTTGCAGGGGTCAGGAAAAGACATTGTGCCAAACTCAGACACCATGATGCGATTCTTCTC | 1122 |
| Pachon2B_ref | CTTGCAGGGGTCAGGAAAAGACATTGTGCCAAACTCAGACACCATGATGCGATTCTTCTC | 1127 |
| Pachon...OG_ref | CTTGCAGGGGTCAGGAAAAGACATTGTGCCAAACTCAGACACCATGATGCGATTCTTCTC | 1121 |
| Pach_12_ref | CTTGCAGGGGTCAGGAAAAGACATTGTGCCAAACTCAGACACCATGATGCGATTCTTCTC | 1112 |
| Pach_12 | CTTGCAGGGGTCAGGAAAAGACATTGTGCCAAACTCAGACACCATGATGCGATTCTTCTC | 1112 |
| Pach_17 | CTTGCAGGGGTCAGGAAAAGACATTGTGCCAAACTCAGACACCATGATGCGATTCTTCTC | 1129 |
| Pachon_F2POG | CTTGCAGGGGTCAGGAAAAGACATTGTGCCAAACTCAGACACCATGATGCGATTCTTCTC | 1125 |
| Pachon_3Gross | CTTGCAGGGGTCAGGAAAAGACATTGTGCCAAACTCAGACACCATGATGCGATTCTTCTC | 1122 |
| Pach_15_ref | CTTGCAGGGGTCAGGAAAAGACATTGTGCCAAACTCAGACACCATGATGCGATTCTTCTC | 1127 |
| Pachon...ro_ref | CTTGCAGGGGTCAGGAAAAGACATTGTGCCAAACTCAGACACCATGATGCGATTCTTCTC | 1129 |
| Pachon...ss_ref | CTTGCAGGGGTCAGGAAAAGACATTGTGCCAAACTCAGACACCATGATGCGATTCTTCTC | 1122 |
| Pachon_2Gross | CTTGCAGGGGTCAGGAAAAGACATTGTGCCAAACTCAGACACCATGATGCGATTCTTCTC | 1173 |
| Pach_17_ref | CTTGCAGGGGTCAGGAAAAGACATTGTGCCAAACTCAGACACCATGATGCGATTCTTCTC | 1129 |
| Pachon...ss_ref | CTTGCAGGGGTCAGGAAAAGACATTGTGCCAAACTCAGACACCATGATGCGATTCTTCTC | 1173 |
| Pachon_ref_ref | CTTGCAGGGGTCAGGAAAAGACATTGTGCCAAACTCAGACACCATGATGCGATTCTTCTC | 1135 |
| Pach_8 | CTTGCAGGGGTCAGGAAAAGACATTGTGCCAAACTCAGACACCATGATGCGATTCTTCTC | 1119 |
| Pach_8_ref | CTTGCAGGGGTCAGGAAAAGACATTGTGCCAAACTCAGACACCATGATGCGATTCTTCTC | 1119 |
| Pachon...OG_ref | CTTGCAGGGGTCAGGAAAAGACATTGTGCCAAACTCAGACACCATGATGCGATTCTTCTC | 1127 |
| Pachon_6Boro | CTTGCAGGGGTCAGGAAAAGACATTGTGCCAAACTCAGACACCATGATGCGATTCTTCTC | 1129 |
| Pachon_ref | CTTGCAGGGGTCAGGAAAAGACATTGTGCCAAACTCAGACACCATGATGCGATTCTTCTC | 1135 |
| Pach_3_ref | CTTGCAGGGGTCAGGAAAAGACATTGTGCCAAACTCAGACACCATGATGCGATTCTTCTC | 1124 |
| Pach_9 | CTTGCAGGGGTCAGGAAAAGACATTGTGCCAAACTCAGACACCATGATGCGATTCTTCTC | 1123 |
| Pachon_G2POG | CTTGCAGGGGTCAGGAAAAGACATTGTGCCAAACTCAGACACCATGATGCGATTCTTCTC | 1121 |
| Pachon...OG_ref | CTTGCAGGGGTCAGGAAAAGACATTGTGCCAAACTCAGACACCATGATGCGATTCTTCTC | 1117 |
| Pach_9_ref | CTTGCAGGGGTCAGGAAAAGACATTGTGCCAAACTCAGACACCATGATGCGATTCTTCTC | 1123 |
| Pachon2B | CTTGCAGGGGTCAGGAAAAGACATTGTGCCAAACTCAGACACCATGATGCGATTCTTCTC | 1127 |
| Pachon...OG_ref | CTTGCAGGGGTCAGGAAAAGACATTGTGCCAAACTCAGACACCATGATGCGATTCTTCTC | 1125 |
| Pachon_E2POG | CTTGCAGGGGTCAGGAAAAGACATTGTGCCAAACTCAGACACCATGATGCGATTCTTCTC | 1127 |
| Pach_14 | CTTGCAGGGGTCAGGAAAAGACATTGTGCCAAACTCAGACACCATGATGCGATTCTTCTC | 1125 |
| Pach_3 | CTTGCAGGGGTCAGGAAAAGACATTGTGCCAAACTCAGACACCATGATGCGATTCTTCTC | 1124 |
| Pachon_H2POG | CTTGCAGGGGTCAGGAAAAGACATTGTGCCAAACTCAGACACCATGATGCGATTCTTCTC | 1117 |
| Pach_7_ref | -TTGCAGGGGTCAGGAAAAGACATTGTGCCAAACTCAGACACCATGATGCGATTCTTCTC | 1059 |
| Pach_15 | CTTGCAGGGGTCAGGAAAAGACATTGTGCCAAACTCAGACACCATGATGCGATTCTTCTC | 1127 |
| Pach_14_ref | CTTGCAGGGGTCAGGAAAAGACATTGTGCCAAACTCAGACACCATGATGCGATTCTTCTC | 1125 |
| Pach_11_ref | CTTGCAGGGGTCAGGAAAAGACATTGTGCCAAACTCAGACACCATGATGCGATTCTTCTC | 1192 |
| Pach_7 | -TTGCAGGGGTCAGGAAAAGACATTGTGCCAAACTCAGACACCATGATGCGATTCTTCTC | 1059 |

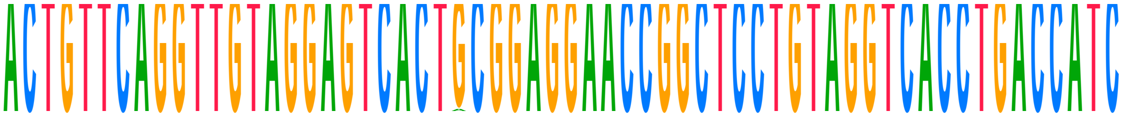

| | | |
|---|---|---|
| **Consensus** | **ACTGTTCAGGTTGTAGGAGTCACTGCGGAGGAACCGGCTCCTGTAGGTCACCTGACCATC** | |
| Choy_1_ref | ACTGTTCAGGTTGTAGGAGTCACTGCGGAGGAACCGGCTCCTGTAGGTCACCTGACCATC | 1378 |
| Choy_1 | ACTGTTCAGGTTGTAGGAGTCACTGCGGAGGAACCGGCTCCTGTAGGTCACCTGACCATC | 1378 |
| Choy_10_ref | ACTGTTCAGGTTGTAGGAGTCACTGCGGAGGAACCGGCTCCTGTAGGTCACCTGACCATC | 1371 |
| Choy_10 | ACTGTTCAGGTTGTAGGAGTCACTGCGGAGGAACCGGCTCCTGTAGGTCACCTGACCATC | 1371 |
| Choy_11_ref | ACTGTTCAGGTTGTAGGAGTCACTGCGGAGGAACCGGCTCCTGTAGGTCACCTGACCATC | 1362 |
| Choy_11 | ACTGTTCAGGTTGTAGGAGTCACTACGGAGGAACCGGCTCCTGTAGGTCACCTGACCATC | 1362 |
| Choy_12_ref | ACTGTTCAGGTTGTAGGAGTCACTGCGGAGGAACCGGCTCCTGTAGGTCACCTGACCATC | 1342 |
| Choy_12 | ACTGTTCAGGTTGTAGGAGTCACTGCGGAGGAACCGGCTCCTGTAGGTCACCTGACCATC | 1342 |
| Choy_13_ref | ACTGTTCAGGTTGTAGGAGTCACTGCGGAGGAACCGGCTCCTGTAGGTCACCTGACCATC | 1380 |
| Choy_13 | ACTGTTCAGGTTGTAGGAGTCACTGCGGAGGAACCGGCTCCTGTAGGTCACCTGACCATC | 1380 |
| Choy_14_ref | ACTGTTCAGGTTGTAGGAGTCACTGCGGAGGAACCGGCTCCTGTAGGTCACCTGACCATC | 1372 |
| Choy_14 | ACTGTTCAGGTTGTAGGAGTCACTGCGGAGGAACCGGCTCCTGTAGGTCACCTGACCATC | 1372 |
| Choy_5_ref | ACTGTTCAGGTTGTAGGAGTCACTGCGGAGGAACCGGCTCCTGTAGGTCACCTGACCATC | 1380 |
| Choy_5 | ACTGTTCAGGTTGTAGGAGTCACTGCGGAGGAACCGGCTCCTGTAGGTCACCTGACCATC | 1380 |
| Choy_6_ref | ACTGTTCAGGTTGTAGGAGTCACTGCGGAGGAACCGGCTCCTGTAGGTCACCTGACCATC | 1380 |
| Choy_6 | ACTGTTCAGGTTGTAGGAGTCACTGCGGAGGAACCGGCTCCTGTAGGTCACCTGACCATC | 1380 |
| Choy_9_ref | ACTGTTCAGGTTGTAGGAGTCACTGCGGAGGAACCGGCTCCTGTAGGTCACCTGACCATC | 1380 |
| Choy_9 | ACTGTTCAGGTTGTAGGAGTCACTGCGGAGGAACCGGCTCCTGTAGGTCACCTGACCATC | 1380 |
| Pachon_wild_bco2a | | |
| Pachon_1Gross | ACTGTTCAGGTTGTAGGAGTCACTGCGGAGGAACCGGCTCCTGTAGGTCACCTGACCATC | 1182 |
| Pach_11 | ACTGTTCAGGTTGTAGGAGTCACTGCGGAGGAACCGGCTCCTGTAGGTCACCTGACCATC | 1252 |
| Pachon...ss_ref | ACTGTTCAGGTTGTAGGAGTCACTGCGGAGGAACCGGCTCCTGTAGGTCACCTGACCATC | 1182 |
| Pachon2B_ref | ACTGTTCAGGTTGTAGGAGTCACTGCGGAGGAACCGGCTCCTGTAGGTCACCTGACCATC | 1187 |
| Pachon...OG_ref | ACTGTTCAGGTTGTAGGAGTCACTGCGGAGGAACCGGCTCCTGTAGGTCACCTGACCATC | 1181 |
| Pach_12_ref | ACTGTTCAGGTTGTAGGAGTCACTGCGGAGGAACCGGCTCCTGTAGGTCACCTGACCATC | 1172 |
| Pach_12 | ACTGTTCAGGTTGTAGGAGTCACTGCGGAGGAACCGGCTCCTGTAGGTCACCTGACCATC | 1172 |
| Pach_17 | ACTGTTCAGGTTGTAGGAGTCACTGCGGAGGAACCGGCTCCTGTAGGTCACCTGACCATC | 1189 |
| Pachon_F2POG | ACTGTTCAGGTTGTAGGAGTCACTGCGGAGGAACCGGCTCCTGTAGGTCACCTGACCATC | 1185 |
| Pachon_3Gross | ACTGTTCAGGTTGTAGGAGTCACTGCGGAGGAACCGGCTCCTGTAGGTCACCTGACCATC | 1182 |
| Pach_15_ref | ACTGTTCAGGTTGTAGGAGTCACTGCGGAGGAACCGGCTCCTGTAGGTCACCTGACCATC | 1187 |
| Pachon...ro_ref | ACTGTTCAGGTTGTAGGAGTCACTGCGGAGGAACCGGCTCCTGTAGGTCACCTGACCATC | 1189 |
| Pachon...ss_ref | ACTGTTCAGGTTGTAGGAGTCACTGCGGAGGAACCGGCTCCTGTAGGTCACCTGACCATC | 1182 |
| Pachon_2Gross | ACTGTTCAGGTTGTAGGAGTCACTGCGGAGGAACCGGCTCCTGTAGGTCACCTGACCATC | 1233 |
| Pach_17_ref | ACTGTTCAGGTTGTAGGAGTCACTGCGGAGGAACCGGCTCCTGTAGGTCACCTGACCATC | 1189 |
| Pachon...ss_ref | ACTGTTCAGGTTGTAGGAGTCACTGCGGAGGAACCGGCTCCTGTAGGTCACCTGACCATC | 1233 |
| Pachon_ref_ref | ACTGTTCAGGTTGTAGGAGTCACTGCGGAGGAACCGGCTCCTGTAGGTCACCTGACCATC | 1195 |
| Pach_8 | ACTGTTCAGGTTGTAGGAGTCACTGCGGAGGAACCGGCTCCTGTAGGTCACCTGACCATC | 1179 |
| Pach_8_ref | ACTGTTCAGGTTGTAGGAGTCACTGCGGAGGAACCGGCTCCTGTAGGTCACCTGACCATC | 1179 |
| Pachon...OG_ref | ACTGTTCAGGTTGTAGGAGTCACTGCGGAGGAACCGGCTCCTGTAGGTCACCTGACCATC | 1187 |
| Pachon_6Boro | ACTGTTCAGGTTGTAGGAGTCACTGCGGAGGAACCGGCTCCTGTAGGTCACCTGACCATC | 1189 |
| Pachon_ref | ACTGTTCAGGTTGTAGGAGTCACTGCGGAGGAACCGGCTCCTGTAGGTCACCTGACCATC | 1195 |
| Pach_3_ref | ACTGTTCAGGTTGTAGGAGTCACTGCGGAGGAACCGGCTCCTGTAGGTCACCTGACCATC | 1184 |
| Pach_9 | ACTGTTCAGGTTGTAGGAGTCACTGCGGAGGAACCGGCTCCTGTAGGTCACCTGACCATC | 1183 |
| Pachon_G2POG | ACTGTTCAGGTTGTAGGAGTCACTGCGGAGGAACCGGCTCCTGTAGGTCACCTGACCATC | 1181 |
| Pachon...OG_ref | ACTGTTCAGGTTGTAGGAGTCACTGCGGAGGAACCGGCTCCTGTAGGTCACCTGACCATC | 1177 |
| Pach_9_ref | ACTGTTCAGGTTGTAGGAGTCACTGCGGAGGAACCGGCTCCTGTAGGTCACCTGACCATC | 1183 |
| Pachon2B | ACTGTTCAGGTTGTAGGAGTCACTGCGGAGGAACCGGCTCCTGTAGGTCACCTGACCATC | 1187 |
| Pachon...OG_ref | ACTGTTCAGGTTGTAGGAGTCACTGCGGAGGAACCGGCTCCTGTAGGTCACCTGACCATC | 1185 |
| Pachon_E2POG | ACTGTTCAGGTTGTAGGAGTCACTGCGGAGGAACCGGCTCCTGTAGGTCACCTGACCATC | 1187 |
| Pach_14 | ACTGTTCAGGTTGTAGGAGTCACTGCGGAGGAACCGGCTCCTGTAGGTCACCTGACCATC | 1185 |
| Pach_3 | ACTGTTCAGGTTGTAGGAGTCACTGCGGAGGAACCGGCTCCTGTAGGTCACCTGACCATC | 1184 |
| Pachon_H2POG | ACTGTTCAGGTTGTAGGAGTCACTGCGGAGGAACCGGCTCCTGTAGGTCACCTGACCATC | 1177 |
| Pach_7_ref | ACTGTTCAGGTTGTAGGAGTCACTGCGGAGGAACCGGCTCCTGTAGGTCACCTGACCATC | 1119 |
| Pach_15 | ACTGTTCAGGTTGTAGGAGTCACTGCGGAGGAACCGGCTCCTGTAGGTCACCTGACCATC | 1187 |
| Pach_14_ref | ACTGTTCAGGTTGTAGGAGTCACTGCGGAGGAACCGGCTCCTGTAGGTCACCTGACCATC | 1185 |
| Pach_11_ref | ACTGTTCAGGTTGTAGGAGTCACTGCGGAGGAACCGGCTCCTGTAGGTCACCTGACCATC | 1252 |
| Pach_7 | ACTGTTCAGGTTGTAGGAGTCACTGCGGAGGAACCGGCTCCTGTAGGTCACCTGACCATC | 1119 |

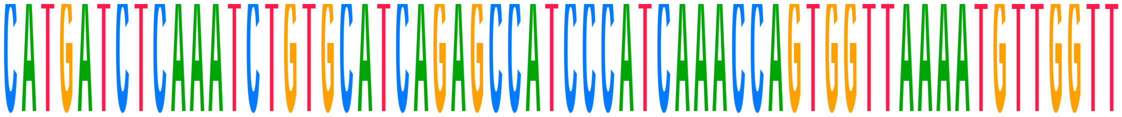

| | | |
|---|---|---|
| **Consensus** | **CATGATCTCAAATCTGTGCATCAGAGCCATCCCATCAAACCAGTGGTTAAAATGTTGGTT** | |
| Choy_1_ref | CATGATCTCAAATCTGTGCATCAGAGCCATCCCATCAAACCAGTGGTTAAAATGTTGGTT | 1438 |
| Choy_1 | CATGATCTCAAATCTGTGCATCAGAGCCATCCCATCAAACCAGTGGTTAAAATGTTGGTT | 1438 |
| Choy_10_ref | CATGATCTCAAATCTGTGCATCAGAGCCATCCCATCAAACCAGTGGTTAAAATGTTGGTT | 1431 |
| Choy_10 | CATGATCTCAAATCTGTGCATCAGAGCCATCCCATCAAACCAGTGGTTAAAATGTTGGTT | 1431 |
| Choy_11_ref | CATGATCTCAAATCTGTGCATCAGAGCCATCCCATCAAACCAGTGGTTAAAA-------- | 1414 |
| Choy_11 | CATGATCTCAAATCTGTGCATCAGAGCCATCCCATCAAACCAGTGGTTAAAA-------- | 1414 |
| Choy_12_ref | CATGATCTCAAATCTGTGCATCAGAGCCATCCCATCAAACCAGTGGTTAAAATGTTGGTT | 1402 |
| Choy_12 | CATGATCTCAAATCTGTGCATCAGAGCCATCCCATCAAACCAGTGGTTAAAATGTTGGTT | 1402 |
| Choy_13_ref | CATGATCTCAAATCTGTGCATCAGAGCCATCCCATCAAACCAGTGGTTAAAATGTTGGTT | 1440 |
| Choy_13 | CATGATCTCAAATCTGTGCATCAGAGCCATCCCATCAAACCAGTGGTTAAAATGTTGGTT | 1440 |
| Choy_14_ref | CATGATCTCAAATCTGTGCATCAGAGCCATCCCATCAAACCAGTGGTTAAAATGTTGGTT | 1432 |
| Choy_14 | CATGATCTCAAATCTGTGCATCAGAGCCATCCCATCAAACCAGTGGTTAAAATGTTGGTT | 1432 |
| Choy_5_ref | CATGATCTCAAATCTGTGCATCAGAGCCATCCCATCAAACCAGTGGTTAAAATGTTGGTT | 1440 |
| Choy_5 | CATGATCTCAAATCTGTGCATCAGAGCCATCCCATCAAACCAGTGGTTAAAATGTTGGTT | 1440 |
| Choy_6_ref | CATGATCTCAAATCTGTGCATCAGAGCCATCCCATCAAACCAGTGGTTAAAATGTTGGTT | 1440 |
| Choy_6 | CATGATCTCAAATCTGTGCATCAGAGCCATCCCATCAAACCAGTGGTTAAAATGTTGGTT | 1440 |
| Choy_9_ref | CATGATCTCAAATCTGTGCATCAGAGCCATCCCATCAAACCAGTGGTTAAAATGTTGGTT | 1440 |
| Choy_9 | CATGATCTCAAATCTGTGCATCAGAGCCATCCCATCAAACCAGTGGTTAAAATGTTGGTT | 1440 |
| Pachon_wild_bco2a | | |
| Pachon_1Gross | CATGATCTCAAATCTGTGCATCAGAGCCATCCCATCAAACCAGTGGTTAAAATGTTGGTT | 1242 |
| Pach_11 | CATGATCTCAAATCTGTGCATCAGAGCCATCCCATCAAACCAGTGGTTAAAATGTTGGTT | 1312 |
| Pachon...ss_ref | CATGATCTCAAATCTGTGCATCAGAGCCATCCCATCAAACCAGTGGTTAAAATGTTGGTT | 1242 |
| Pachon2B_ref | CATGATCTCAAATCTGTGCATCAGAGCCATCCCATCAAACCAGTGGTTAAAATGTTGGTT | 1247 |
| Pachon...OG_ref | CATGATCTCAAATCTGTGCATCAGAGCCATCCCATCAAACCAGTGGTTAAAATGTTGGTT | 1241 |
| Pach_12_ref | CATGATCTCAAATCTGTGCATCAGAGCCATCCCATCAAACCAGTGGTTAAAATGTTGGTT | 1232 |
| Pach_12 | CATGATCTCAAATCTGTGCATCAGAGCCATCCCATCAAACCAGTGGTTAAAATGTTGGTT | 1232 |
| Pach_17 | CATGATCTCAAATCTGTGCATCAGAGCCATCCCATCAAACCAGTGGTTAAAA-------- | 1241 |
| Pachon_F2POG | CATGATCTCAAATCTGTGCATCAGAGCCATCCCATCAAACCAGTGGTTAAAATGTTGGTT | 1245 |
| Pachon_3Gross | CATGATCTCAAATCTGTGCATCAGAGCCATCCCATCAAACCAGTGGTTAAAATGTTGGTT | 1242 |
| Pach_15_ref | CATGATCTCAAATCTGTGCATCAGAGCCATCCCATCAAACCAGTGGTTAAAATGTTGGTT | 1247 |
| Pachon...ro_ref | CATGATCTCAAATCTGTGCATCAGAGCCATCCCATCAAACCAGTGGTTAAAATGTTGGTT | 1249 |
| Pachon...ss_ref | CATGATCTCAAATCTGTGCATCAGAGCCATCCCATCAAACCAGTGGTTAAAATGTTGGTT | 1242 |
| Pachon_2Gross | CATGATCTCAAATCTGTGCATCAGAGCCATCCCATCAAACCAGTGGTTAAAATGTTGGTT | 1293 |
| Pach_17_ref | CATGATCTCAAATCTGTGCATCAGAGCCATCCCATCAAACCAGTGGTTAAAA-------- | 1241 |
| Pachon...ss_ref | CATGATCTCAAATCTGTGCATCAGAGCCATCCCATCAAACCAGTGGTTAAAATGTTGGTT | 1293 |
| Pachon_ref_ref | CATGATCTCAAATCTGTGCATCAGAGCCATCCCATCAAACCAGTGGTTAAAATGTTGGTT | 1255 |
| Pach_8 | CATGATCTCAAATCTGTGCATCAGAGCCATCCCATCAAACCAGTGGTTAAAATGTTGGTT | 1239 |
| Pach_8_ref | CATGATCTCAAATCTGTGCATCAGAGCCATCCCATCAAACCAGTGGTTAAAATGTTGGTT | 1239 |
| Pachon...OG_ref | CATGATCTCAAATCTGTGCATCAGAGCCATCCCATCAAACCAGTGGTTAAAATGTTGGTT | 1247 |
| Pachon_6Boro | CATGATCTCAAATCTGTGCATCAGAGCCATCCCATCAAACCAGTGGTTAAAATGTTGGTT | 1249 |
| Pachon_ref | CATGATCTCAAATCTGTGCATCAGAGCCATCCCATCAAACCAGTGGTTAAAATGTTGGTT | 1255 |
| Pach_3_ref | CATGATCTCAAATCTGTGCATCAGAGCCATCCCATCAAACCAGTGGTTAAAATGTTGGTT | 1244 |
| Pach_9 | CATGATCTCAAATCTGTGCATCAGAGCCATCCCATCAAACCAGTGGTTAAAATGTTGGTT | 1243 |
| Pachon_G2POG | CATGATCTCAAATCTGTGCATCAGAGCCATCCCATCAAACCAGTGGTTAAAATGTTGGTT | 1241 |
| Pachon...OG_ref | CATGATCTCAAATCTGTGCATCAGAGCCATCCCATCAAACCAGTGGTTAAAATGTTGGTT | 1237 |
| Pach_9_ref | CATGATCTCAAATCTGTGCATCAGAGCCATCCCATCAAACCAGTGGTTAAAATGTTGGTT | 1243 |
| Pachon2B | CATGATCTCAAATCTGTGCATCAGAGCCATCCCATCAAACCAGTGGTTAAAATGTTGGTT | 1247 |
| Pachon...OG_ref | CATGATCTCAAATCTGTGCATCAGAGCCATCCCATCAAACCAGTGGTTAAAATGTTGGTT | 1245 |
| Pachon_E2POG | CATGATCTCAAATCTGTGCATCAGAGCCATCCCATCAAACCAGTGGTTAAAATGTTGGTT | 1247 |
| Pach_14 | CATGATCTCAAATCTGTGCATCAGAGCCATCCCATCAAACCAGTGGTTAAAATGTTGGTT | 1245 |
| Pach_3 | CATGATCTCAAATCTGTGCATCAGAGCCATCCCATCAAACCAGTGGTTAAAATGTTGGTT | 1244 |
| Pachon_H2POG | CATGATCTCAAATCTGTGCATCAGAGCCATCCCATCAAACCAGTGGTTAAAATGTTGGTT | 1237 |
| Pach_7_ref | CATGATCTCAAATCTGTGCATCAGAGCCATCCCATCAAACCAGTGGTTAAAATGTTGGTT | 1179 |
| Pach_15 | CATGATCTCAAATCTGTGCATCAGAGCCATCCCATCAAACCAGTGGTTAAAATGTTGGTT | 1247 |
| Pach_14_ref | CATGATCTCAAATCTGTGCATCAGAGCCATCCCATCAAACCAGTGGTTAAAATGTTGGTT | 1245 |
| Pach_11_ref | CATGATCTCAAATCTGTGCATCAGAGCCATCCCATCAAACCAGTGGTTAAAATGTTGGTT | 1312 |
| Pach_7 | CATGATCTCAAATCTGTGCATCAGAGCCATCCCATCAAACCAGTGGTTAAAATGTTGGTT | 1179 |

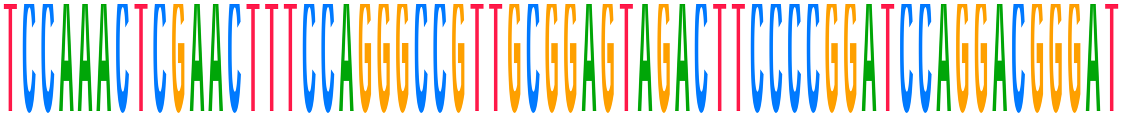

| | | |
|---|---|---|
| **Consensus** | **TCCAAACTCGAACTTTCCAGGGCCGTTGCGGAGTAGACTTCCCCGGATCCAGGACGGGAT** | |
| Choy_1_ref | TCCAAACTCGAACTTTCCAGGGC------------------------------------- | 1461 |
| Choy_1 | TCCAAACTCGAACTTTCCAGGGC------------------------------------- | 1461 |
| Choy_10_ref | TCCAAACTCGAACTTTCCAGGGCCGTTGCGGAGTAGACTTCCCCGGATCCAGGACGGGAT | 1491 |
| Choy_10 | TCCAAACTCGAACTTTCCAGGGCCGTTGCGGAGTAGACTTCCCCGGATCCAGGACGGGAT | 1491 |
| Choy_11_ref | --CAAACTCGAACTTTCCAGGGCCGTTGCGGAGTAGACTTCCCCGGATCCAGGA-GGGAT | 1471 |
| Choy_11 | --CAAACTCGAACTTTCCAGGGCCGTTGCGGAGTAGACTTCCCCGGATCCAGGA-GGGAT | 1471 |
| Choy_12_ref | TCCAAACTCGAACTTTCCAGGGCCGTTGCGGAGTAGACTTCCCCGGATCCAGGACGGGAT | 1462 |
| Choy_12 | TCCAAACTCGAACTTTCCAGGGCCGTTGCGGAGTAGACTTCCCCGGATCCAGGACGGGAT | 1462 |
| Choy_13_ref | TCCAAACTCGAACTTTCCAGGGCCGTTGCGGAGTAGACTTCCCCGGATCCAGGACGGGAT | 1500 |
| Choy_13 | TCCAAACTCGAACTTTCCAGGGCCGTTGCGGAGTAGACTTCCCCGGATCCAGGACGGGAT | 1500 |
| Choy_14_ref | TCCAAACTCGAACTTTCCAGGGCCGTTGCGGAGTAGACTTCCCCGGATCCAGGACGGGAT | 1492 |
| Choy_14 | TCCAAACTCGAACTTTCCAGGGCCGTTGCGGAGTAGACTTCCCCGGATCCAGGACGGGAT | 1492 |
| Choy_5_ref | TCCAAACTCGAACTTTCCAGGGCCGTTGCGGAGTAGACTTCCCCGGATCCAGGACGGGAT | 1500 |
| Choy_5 | TCCAAACTCGAACTTTCCAGGGCCGTTGCGGAGTAGACTTCCCCGGATCCAGGACGGGAT | 1500 |
| Choy_6_ref | TCCAAACTCGAACTTTCCAGGGCCGTTGCGGAGTAGACTTCCCCGGATCCAGGACGGGAT | 1500 |
| Choy_6 | TCCAAACTCGAACTTTCCAGGGCCGTTGCGGAGTAGACTTCCCCGGATCCAGGACGGGAT | 1500 |
| Choy_9_ref | TCCAAACTCGAACTTTCCAGGGCCGTTGCGGAGTAGACTTCCCCGGATCCAGGACGGGAT | 1500 |
| Choy_9 | TCCAAACTCGAACTTTCCAGGGCCGTTGCGGAGTAGACTTCCCCGGATCCAGGACGGGAT | 1500 |
| Pachon_wild_bco2a | | |
| Pachon_1Gross | TCCAAACTCGAACTTTCCAGGGCCGTTGCGGAGTAGACTTCCCCGGATCCAGGACGGGAT | 1302 |
| Pach_11 | TCCAAACTCGAA---TCCAGGGCCGTTGCGGAGTAGACTTCCCCGGATCCAGGACGGGAT | 1369 |
| Pachon…ss_ref | TCCAAACTCGAACTTTCCAGGGCCGTTGCGGAGTAGACTTCCCCGGATCCAGGACGGGAT | 1302 |
| Pachon2B_ref | TCCAAACTCGAACTTTCCAGGGCCGTTGCGGAGTAGACTTCCCCGGATCCAGGACGGGAT | 1307 |
| Pachon…OG_ref | TCCAAACTCGAACTTTCCAGGGCCGTTGCGGAGTAGACTTCCCCGGATCCAGGACGGGAT | 1301 |
| Pach_12_ref | TCCAAACTCGAACTTTCCAGGGCCGTTGCGGAGTAGACTTCCCCGGATCCAGGACGGGAT | 1292 |
| Pach_12 | TCCAAACTCGAACTTTCCAGGGCCGTTGCGGAGTAGACTTCCCCGGATCCAGGACGGGAT | 1292 |
| Pach_17 | ---------------------GCCGTTGCGGAGTAGACTTCCCCGGATCCAGGACGGGAT | 1280 |
| Pachon_F2POG | TCCAAACTCGAACTTTCCAGGGCCGTTGCGGAGTAGACTTCCCCGGATCCAGGACGGGAT | 1305 |
| Pachon_3Gross | TCCAAACTCGAACTTTCCAGGGCCGTTGCGGAGTAGACTTCCCCGGATCCAGGACGGGAT | 1302 |
| Pach_15_ref | TCCAAACTCGAACTTTCCAGGGCCGTTGCGGAGTAGACTTCCCCGGATCCAGGACGGGAT | 1307 |
| Pachon…ro_ref | TCCAAACTCGAACTTTCCAGGGCCGTTGCGGAGTAGACTTCCCCGGATCCAGGACGGGAT | 1309 |
| Pachon…ss_ref | TCCAAACTCGAACTTTCCAGGGCCGTTGCGGAGTAGACTTCCCCGGATCCAGGACGGGAT | 1302 |
| Pachon_2Gross | TCCAAACTCGAACTTTCCAGGGCCGTTGCGGAGTAGACTTCCCCGGATCCAGGACGGGAT | 1353 |
| Pach_17_ref | ---------------------GCCGTTGCGGAGTAGACTTCCCCGGATCCAGGACGGGAT | 1280 |
| Pachon…ss_ref | TCCAAACTCGAACTTTCCAGGGCCGTTGCGGAGTAGACTTCCCCGGATCCAGGACGGGAT | 1353 |
| Pachon_ref_ref | TCCAAACTCGAACTTTCCAGGGCCGTTGCGGAGTAGACTTCCCCGGATCCAGGACGGGAT | 1315 |
| Pach_8 | TCCAAACTCGAACTTTCCAGGGCCGTTGCGGAGTAGACTTCCCCGGATCCAGGACGGGAT | 1299 |
| Pach_8_ref | TCCAAACTCGAACTTTCCAGGGCCGTTGCGGAGTAGACTTCCCCGGATCCAGGACGGGAT | 1299 |
| Pachon…OG_ref | TCCAAACTCGAACTTTCCAGGGCCGTTGCGGAGTAGACTTCCCCGGATCCAGGACGGGAT | 1307 |
| Pachon_6Boro | TCCAAACTCGAACTTTCCAGGGCCGTTGCGGAGTAGACTTCCCCGGATCCAGGACGGGAT | 1309 |
| Pachon_ref | TCCAAACTCGAACTTTCCAGGGCCGTTGCGGAGTAGACTTCCCCGGATCCAGGACGGGAT | 1315 |
| Pach_3_ref | TCCAAACTCGAACTTTCCAGGGCCGTTGCGGAGTAGACTTCCCCGGATCCAGGACGGGAT | 1304 |
| Pach_9 | TCCAAACTCGAACTTTCCAGGGCCGTTGCGGAGTAGACTTCCCCGGATCCAGGACGGGAT | 1303 |
| Pachon_G2POG | TCCAAACTCGAACTTTCCAGGGCCGTTGCGGAGTAGACTTCCCCGGATCCAGGACGGGAT | 1301 |
| Pachon…OG_ref | TCCAAACTCGAACTTTCCAGGGCCGTTGCGGAGTAGACTTCCCCGGATCCAGGACGGGAT | 1297 |
| Pach_9_ref | TCCAAACTCGAACTTTCCAGGGCCGTTGCGGAGTAGACTTCCCCGGATCCAGGACGGGAT | 1303 |
| Pachon2B | TCCAAACTCGAACTTTCCAGGGCCGTTGCGGAGTAGACTTCCCCGGATCCAGGACGGGAT | 1307 |
| Pachon…OG_ref | TCCAAACTCGAACTTTCCAGGGCCGTTGCGGAGTAGACTTCCCCGGATCCAGGACGGGAT | 1305 |
| Pachon_E2POG | TCCAAACTCGAACTTTCCAGGGCCGTTGCGGAGTAGACTTCCCCGGATCCAGGACGGGAT | 1307 |
| Pach_14 | TCCAAACTCGAACTTTCCAGGGCCGTTGCGGAGTAGACTTCCCCGGATCCAGGACGGGAT | 1305 |
| Pach_3 | TCCAAACTCGAACTTTCCAGGGCCGTTGCGGAGTAGACTTCCCCGGATCCAGGACGGGAT | 1304 |
| Pachon_H2POG | TCCAAACTCGAACTTTCCAGGGCCGTTGCGGAGTAGACTTCCCCGGATCCAGGACGGGAT | 1297 |
| Pach_7_ref | TCCAAACTCGAACTTTCCAGGGCCGTTGCGGAGTAGACTTCCCCGGATCCAGGACGGGAT | 1239 |
| Pach_15 | TCCAAACTCGAACTTTCCAGGGCCGTTGCGGAGTAGACTTCCCCGGATCCAGGACGGGAT | 1307 |
| Pach_14_ref | TCCAAACTCGAACTTTCCAGGGCCGTTGCGGAGTAGACTTCCCCGGATCCAGGACGGGAT | 1305 |
| Pach_11_ref | TCCAAACTCGAA---TCCAGGGCCGTTGCGGAGTAGACTTCCCCGGATCCAGGACGGGAT | 1369 |
| Pach_7 | TCCAAACTCGAACTTTCCAGGGCCGTTGCGGAGTAGACTTCCCCGGATCCAGGACGGGAT | 1239 |

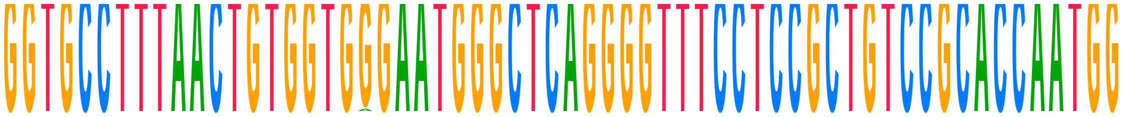

| | | |
|---|---|---|
| **Consensus** | **GGTGCCTTTAACTGTGGTGGGAATGGGCTCAGGGGTTTCCTCCGCTGTCCGCACCAATGG** | |
| Choy_1_ref | -----------------------------------TTTCCTCCGCTGTCCGCACCAATGG | 1486 |
| Choy_1 | -----------------------------------TTTCCTCCGCTGTCCGCACCAATGG | 1486 |
| Choy_10_ref | GGTGCCTTTAACTGTGGTGGGAATGGGCTCAGGGGTTTCCTCCGCTGTCCGCACCAATGG | 1551 |
| Choy_10 | GGTGCCTTTAACTGTGGTGGGAATGGGCTCAGGGGTTTCCTCCGCTGTCCGCACCAATGG | 1551 |
| Choy_11_ref | GGTGCCTTTAACTGTGGTGGGAATGGGCTCAGGGGTTTCCTCCGCTGTCCGCACCAATGG | 1531 |
| Choy_11 | GGTGCCTTTAACTGTGGTGGGAATGGGCTCAGGGGTTTCCTCCGCTGTCCGCACCAATGG | 1531 |
| Choy_12_ref | GGTGCCTTTAA------------------------------CCGCTGTCCGCACCAATGG | 1492 |
| Choy_12 | GGTGCCTTTAA------------------------------CCGCTGTCCGCACCAATGG | 1492 |
| Choy_13_ref | GGTGCCTTTAACTGTGGTGGGAATGGGCTCAGGGGTTTCCTCCGCTGTCCGCACCAATGG | 1560 |
| Choy_13 | GGTGCCTTTAACTGTGGTGGGAATGGGCTCAGGGGTTTCCTCCGCTGTCCGCACCAATGG | 1560 |
| Choy_14_ref | GGTGCCTTTAACTGTGGTGGGAATGGGCTCAGGGGTTTCCTCCGCTGTCCGCACCAATGG | 1552 |
| Choy_14 | GGTGCCTTTAACTGTGGTGGGAATGGGCTCAGGGGTTTCCTCCGCTGTCCGCACCAATGG | 1552 |
| Choy_5_ref | GGTGCCTTTAACTGTGGTGGGAATGGGCTCAGGGGTTTCCTCCGCTGTCCGCACCAATGG | 1560 |
| Choy_5 | GGTGCCTTTAACTGTGGTGGGAATGGGCTCAGGGGTTTCCTCCGCTGTCCGCACCAATGG | 1560 |
| Choy_6_ref | GGTGCCTTTAACTGTGGTGGGAATGGGCTCAGGGGTTTCCTCCGCTGTCCGCACCAATGG | 1560 |
| Choy_6 | GGTGCCTTTAACTGTGGTGGGAATGGGCTCAGGGGTTTCCTCCGCTGTCCGCACCAATGG | 1560 |
| Choy_9_ref | GGTGCCTTT-ACTGTGGTGGGAATGGGC-------------------------------- | 1527 |
| Choy_9 | GGTGCCTTT-ACTGTGGTGGGAATGGGC-------------------------------- | 1527 |
| Pachon_wild_bco2a | | |
| Pachon_1Gross | GGTGCCTTTAACTGTGGTGGGAATGGGCTCAGGGGTTTCCTCCGCTGTCCGCACCAATGG | 1362 |
| Pach_11 | GGTGCCTTTAACTGTGGTGGGAATGGGCTCAGGGGTTTCCTCCGCTGTCCGCACCAATGG | 1429 |
| Pachon...ss_ref | GGTGCCTTTAACTGTGGTGGGAATGGGCTCAGGGGTTTCCTCCGCTGTCCGCACCAATGG | 1362 |
| Pachon2B_ref | GGTGCCTTTAACTGTGGTGGGAATGGGCTCAGGGGTTTCCTCCGCTGTCCGCACCAATGG | 1367 |
| Pachon...OG_ref | GGTGCCTTTAACTGTGGTGGGAATGGGCTCAGGGGTTTCCTCCGCTGTCCGCACCAATGG | 1361 |
| Pach_12_ref | GGTGCCTTTAACTGTGGTGGGAATGGGCTCAGGGGTTTCCTCCGCTGTCCGCACCAATGG | 1352 |
| Pach_12 | GGTGCCTTTAACTGTGGTGGGAATGGGCTCAGGGGTTTCCTCCGCTGTCCGCACCAATGG | 1352 |
| Pach_17 | GGTGCCTTTAACTGTGGTGGGAAT----------GTTTCCTCCGCTGTCCGCACCAATGG | 1330 |
| Pachon_F2POG | GGTGCCTTTAACTGTGGTGAGAATGGGCTCAGGGGTTTCCTCCGCTGTCCGCACCAATGG | 1365 |
| Pachon_3Gross | GGTGCCTTTAACTGTGGTGGGAATGGGCTCAGGGGTTTCCTCCGCTGTCCGCACCAATGG | 1362 |
| Pach_15_ref | GGTGCCTTTAACTGTGGTGGGAATGGGCTCAGGGGTTTCCTCCGCTGTCCGCACCAATGG | 1367 |
| Pachon...ro_ref | GGTGCCTTTAACTGTGGTGGGAATGGGCTCAGGGGTTTCCTCCGCTGTCCGCACCAATGG | 1369 |
| Pachon...ss_ref | GGTGCCTTTAACTGTGGTGGGAATGGGCTCAGGGGTTTCCTCCGCTGTCCGCACCAATGG | 1362 |
| Pachon_2Gross | GGTGCCTTTAACTGTGGTGGGAATGGGCTCAGGGGTTTCCTCCGCTGTCCGCACCAATGG | 1413 |
| Pach_17_ref | GGTGCCTTTAACTGTGGTGGGAAT----------GTTTCCTCCGCTGTCCGCACCAATGG | 1330 |
| Pachon...ss_ref | GGTGCCTTTAACTGTGGTGGGAATGGGCTCAGGGGTTTCCTCCGCTGTCCGCACCAATGG | 1413 |
| Pachon_ref_ref | GGTGCCTTTAACTGTGGTGGGAATGGGCTCAGGGGTTTCCTCCGCTGTCCGCACCAATGG | 1375 |
| Pach_8 | GGTGCCTTTAACTGTGGTGGGAATGGGCTCAGGGGTTTCCTCCGCTGTCCGCACCAATGG | 1359 |
| Pach_8_ref | GGTGCCTTTAACTGTGGTGGGAATGGGCTCAGGGGTTTCCTCCGCTGTCCGCACCAATGG | 1359 |
| Pachon...OG_ref | GGTGCCTTTAACTGTGGTGGGAATGGGCTCAGGGGTTTCCTCCGCTGTCCGCACCAATGG | 1367 |
| Pachon_6Boro | GGTGCCTTTAACTGTGGTGGGAATGGGCTCAGGGGTTTCCTCCGCTGTCCGCACCAATGG | 1369 |
| Pachon_ref | GGTGCCTTTAACTGTGGTGGGAATGGGCTCAGGGGTTTCCTCCGCTGTCCGCACCAATGG | 1375 |
| Pach_3_ref | GGTGCCTTTAACTGTGGTGGGAATGGGCTCAGGGGTTTCCTCCGCTGTCCGCACCAATGG | 1364 |
| Pach_9 | GGTGCCTTTAACTGTGGTGGGAATGGGCTCAGGGGTTTCCTCCGCTGTCCGCACCAATGG | 1363 |
| Pachon_G2POG | GGTGCCTTTAACTGTGGTGGGAATGGGCTCAGGGGTTTCCTCCGCTGTCCGCACCAATGG | 1361 |
| Pachon...OG_ref | GGTGCCTTTAACTGTGGTGGGAATGGGCTCAGGGGTTTCCTCCGCTGTCCGCACCAATGG | 1357 |
| Pach_9_ref | GGTGCCTTTAACTGTGGTGGGAATGGGCTCAGGGGTTTCCTCCGCTGTCCGCACCAATGG | 1363 |
| Pachon2B | GGTGCCTTTAACTGTGGTGGGAATGGGCTCAGGGGTTTCCTCCGCTGTCCGCACCAATGG | 1367 |
| Pachon...OG_ref | GGTGCCTTTAACTGTGGTGGGAATGGGCTCAGGGGTTTCCTCCGCTGTCCGCACCAATGG | 1365 |
| Pachon_E2POG | GGTGCCTTTAACTGTGGTGGGAATGGGCTCAGGGGTTTCCTCCGCTGTCCGCACCAATGG | 1367 |
| Pach_14 | GGTGCCT-------------GAATGGGCTCAGGGGTTTCCTCCGCTGTCCGCACCAATGG | 1352 |
| Pach_3 | GGTGCCTTTAACTGTGGTGGGAATGGGCTCAGGGGTTTCCTCCGCTGTCCGCACCAATGG | 1364 |
| Pachon_H2POG | GGTGCCTTTAACTGTGGTGGGAATGGGCTCAGGGGTTTCCTCCGCTGTCCGCACCAATGG | 1357 |
| Pach_7_ref | GGTGCCTTTAACTGTGGTGGGAATGGGCTCAGGGGTTTCCTCCGCTGTCCGCACCAATGG | 1299 |
| Pach_15 | GGTGCCTTTAACTGTGGTGGGAATGGGCTCAGGGGTTTCCTCCGCTGTCCGCACCAATGG | 1367 |
| Pach_14_ref | GGTGCCT-------------GAATGGGCTCAGGGGTTTCCTCCGCTGTCCGCACCAATGG | 1352 |
| Pach_11_ref | GGTGCCTTTAACTGTGGTGGGAATGGGCTCAGGGGTTTCCTCCGCTGTCCGCACCAATGG | 1429 |
| Pach_7 | GGTGCCTTTAACTGTGGTGGGAATGGGCTCAGGGGTTTCCTCCGCTGTCCGCACCAATGG | 1299 |

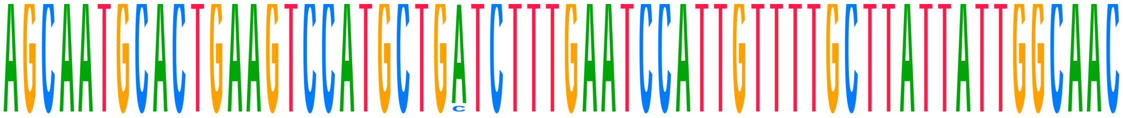

| | | |
|---|---|---|
| **Consensus** | **AGCAATGCACTGAAGTCCATGCTGATCTTTGAATCCATTGTTTTGCTTATTATTGGCAAC** | |
| Choy_1_ref | AGCAATGCACTGAAGTCCATGCTGATCTTTGAATCCATTGTTTTGCTTATTATTGGCAAC | 1546 |
| Choy_1 | AGCAATGCACTGAAGTCCATGCTGATCTTTGAATCCATTGTTTTGCTTATTATTGGCAAC | 1546 |
| Choy_10_ref | AGCAATGCACTGAAGTCCATGCTGATCTTTGAATCCATTGTTTTGCTTATTATTGGCAAC | 1611 |
| Choy_10 | AGCAATGCACTGAAGTCCATGCTGATCTTTGAATCCATTGTTTTGCTTATTATTGGCAAC | 1611 |
| Choy_11_ref | AGCAATGCACTGAAGTCCATGCTGATCTTTGAATCCATTGTTTTGCTTATTATTGGCAAC | 1591 |
| Choy_11 | AGCAATGCACTGAAGTCCATGCTGCTCTTTGAATCCATTGTTTTGCTTATTATTGGCAAC | 1591 |
| Choy_12_ref | AGCAATGCACTGAAGTCCATGCTGATCTTTGAATCCATTGTTTTGCTTATTATTGGCAAC | 1552 |
| Choy_12 | AGCAATGCACTGAAGTCCATGCTGATCTTTGAATCCATTGTTTTGCTTATTATTGGCAAC | 1552 |
| Choy_13_ref | AGCAATGCACTGAAGTCCATGCTGATCTTTGAATCCATTGTTTTGCTTATTATTGGCAAC | 1620 |
| Choy_13 | AGCAATGCACTGAAGTCCATGCTGCTCTTTGAATCCATTGTTTTGCTTATTATTGGCAAC | 1620 |
| Choy_14_ref | AGCAATGCACTGAAGTCCATGCTGATCTTTGAATCCATTGTTTTGCTTATTATTGGCAAC | 1612 |
| Choy_14 | AGCAATGCACTGAAGTCCATGCTGCTCTTTGAATCCATTGTTTTGCTTATTATTGGCAAC | 1612 |
| Choy_5_ref | AGCAATGCACTGAAGTCCATGCTGATCTTTGAATCCATTGTTTTGCTTATTATTGGCAAC | 1620 |
| Choy_5 | AGCAATGCACTGAAGTCCATGCTGATCTTTGAATCCATTGTTTTGCTTATTATTGGCAAC | 1620 |
| Choy_6_ref | AGCAATGCACTGAAGTCCATGCTGATCTTTGAATCCATTGTTTTGCTTATTATTGGCAAC | 1620 |
| Choy_6 | AGCAATGCACTGAAGTCCATGCTGATCTTTGAATCCATTGTTTTGCTTATTATTGGCAAC | 1620 |
| Choy_9_ref | ---------------TCCATGCTGATCTTTGAATCCATTGTTTTGCTTATTATTGGCAAC | 1572 |
| Choy_9 | ---------------TCCATGCTGATCTTTGAATCCATTGTTTTGCTTATTATTGGCAAC | 1572 |
| Pachon_wild_bco2a | | |
| Pachon_1Gross | AGCAATGCACTGAAGTCCATGCTGATCTTTGAATCCATTGTTTTGCTTATTATTGGCAAC | 1422 |
| Pach_11 | AGCAATGCACTGAAGTCCATGCTGATCTTTGAATCCATTGTTTTGCTTATTATTGGCAAC | 1489 |
| Pachon...ss_ref | AGCAATGCACTGAAGTCCATGCTGATCTTTGAATCCATTGTTTTGCTTATTATTGGCAAC | 1422 |
| Pachon2B_ref | AGCAATGCACTGAAGTCCATGCTGATCTTTGAATCCATTGTTTTGCTTATTATTGGCAAC | 1427 |
| Pachon...OG_ref | AGCAATGCACTGAAGTCCATGCTGATCTTTGAATCCATTGTTTTGCTTATTATTGGCAAC | 1421 |
| Pach_12_ref | AGCAATGCACTGAAGTCCATGCTGATCTTTGAATCCATTGTTTTGCTTATTATTGGCAAC | 1412 |
| Pach_12 | AGCAATGCACTGAAGTCCATGCTGATCTTTGAATCCATTGTTTTGCTTATTATTGGCAAC | 1412 |
| Pach_17 | AGCAATGCACTGAAGTCCATGCTGATCTTTGAATCCATTGTTTTGCTTATTATTGGCAAC | 1390 |
| Pachon_F2POG | AGCAATGCACTGAAGTCCATGCTGATCTTTGAATCCATTGTTTTGCTTATTATTGGCAAC | 1425 |
| Pachon_3Gross | AGCAATGCACTGAAGTCCATGCTGATCTTTGAATCCATTGTTTTGCTTATTATTGGCAAC | 1422 |
| Pach_15_ref | AGCAATGCACTGAAGTCCATGCTGATCTTTGAATCCATTGTTTTGCTTATTATTGGCAAC | 1427 |
| Pachon...ro_ref | AGCAATGCACTGAAGTCCATGCTGATCTTTGAATCCATTGTTTTGCTTATTATTGGCAAC | 1429 |
| Pachon...ss_ref | AGCAATGCACTGAAGTCCATGCTGATCTTTGAATCCATTGTTTTGCTTATTATTGGCAAC | 1422 |
| Pachon_2Gross | AGCAATGCACTGAAGTCCATGCTGATCTTTGAATCCATTGTTTTGCTTATTATTGGCAAC | 1473 |
| Pach_17_ref | AGCAATGCACTGAAGTCCATGCTGATCTTTGAATCCATTGTTTTGCTTATTATTGGCAAC | 1390 |
| Pachon...ss_ref | AGCAATGCACTGAAGTCCATGCTGATCTTTGAATCCATTGTTTTGCTTATTATTGGCAAC | 1473 |
| Pachon_ref_ref | AGCAATGCACTGAAGTCCATGCTGATCTTTGAATCCATTGTTTTGCTTATTATTGGCAAC | 1435 |
| Pach_8 | AGCAATGCACTGAAGTCCATGCTGATCTTTGAATCCATTGTTTTGCTTATTATTGGCAAC | 1419 |
| Pach_8_ref | AGCAATGCACTGAAGTCCATGCTGATCTTTGAATCCATTGTTTTGCTTATTATTGGCAAC | 1419 |
| Pachon...OG_ref | AGCAATGCACTGAAGTCCATGCTGATCTTTGAATCCATTGTTTTGCTTATTATTGGCAAC | 1427 |
| Pachon_6Boro | AGCAATGCACTGAAGTCCATGCTGATCTTTGAATCCATTGTTTTGCTTATTATTGGCAAC | 1429 |
| Pachon_ref | AGCAATGCACTGAAGTCCATGCTGATCTTTGAATCCATTGTTTTGCTTATTATTGGCAAC | 1435 |
| Pach_3_ref | AGCAATGCACTGAAGTCCATGCTGATCTTTGAATCCATTGTTTTGCTTATTATTGGCAAC | 1424 |
| Pach_9 | AGCAATGCACTGAAGTCCATGCTGATCTTTGAATCCATTGTTTTGCTTATTATTGGCAAC | 1423 |
| Pachon_G2POG | AGCAATGCACTGAAGTCCATGCTGATCTTTGAATCCATTGTTTTGCTTATTATTGGCAAC | 1421 |
| Pachon...OG_ref | AGCAATGCACTGAAGTCCATGCTGATCTTTGAATCCATTGTTTTGCTTATTATTGGCAAC | 1417 |
| Pach_9_ref | AGCAATGCACTGAAGTCCATGCTGATCTTTGAATCCATTGTTTTGCTTATTATTGGCAAC | 1423 |
| Pachon2B | AGCAATGCACTGAAGTCCATGCTGATCTTTGAATCCATTGTTTTGCTTATTATTGGCAAC | 1427 |
| Pachon...OG_ref | AGCAATGCACTGAAGTCCATGCTGATCTTTGAATCCATTGTTTTGCTTATTATTGGCAAC | 1425 |
| Pachon_E2POG | AGCAATGCACTGAAGTCCATGCTGATCTTTGAATCCATTGTTTTGCTTATTATTGGCAAC | 1427 |
| Pach_14 | AGCAATGCACTGAAGTCCATGCTGATCTTTGAATCCATTGTTTTGCTTATTATTGGCAAC | 1412 |
| Pach_3 | AGCAATGCACTGAAGTCCATGCTGATCTTTGAATCCATTGTTTTGCTTATTATTGGCAAC | 1424 |
| Pachon_H2POG | AGCAATGCACTGAAGTCCATGCTGATCTTTGAATCCATTGTTTTGCTTATTATTGGCAAC | 1417 |
| Pach_7_ref | AGCAATGCACTGAAGTCCATGCTGATCTTTGAATCCATTGTTTTGCTTATTATTGGCAAC | 1359 |
| Pach_15 | AGCAATGCACTGAAGTCCATGCTGATCTTTGAATCCATTGTTTTGCTTATTATTGGCAAC | 1427 |
| Pach_14_ref | AGCAATGCACTGAAGTCCATGCTGATCTTTGAATCCATTGTTTTGCTTATTATTGGCAAC | 1412 |
| Pach_11_ref | AGCAATGCACTGAAGTCCATGCTGATCTTTGAATCCATTGTTTTGCTTATTATTGGCAAC | 1489 |
| Pach_7 | AGCAATGCACTGAAGTCCATGCTGATCTTTGAATCCATTGTTTTGCTTATTATTGGCAAC | 1359 |

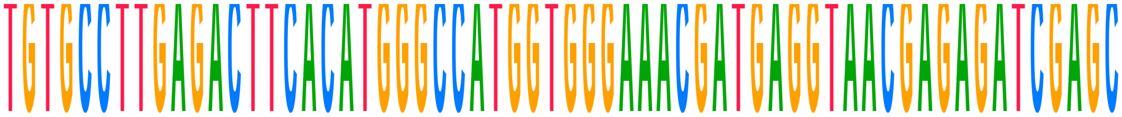

| | | |
|---|---|---|
| **Consensus** | **TGTGCCTTGAGACTTCACATGGGCCATGGTGGGAAACGATGAGGTAACGAGAGATCGAGC** | |
| Choy_1_ref | TGTGCCTTGAGACTTCACATGGGCCATGGTGGGAAACGATGAGGTAACGAGAGATCGAGC | 1606 |
| Choy_1 | TGTGCCTTGAGACTTCACATGGGCCATGGTGGGAAACGATGAGGTAACGAGAGATCGAGC | 1606 |
| Choy_10_ref | TGTGCCTTGAGACTTCACATGGGCCATGGTGGGAAACGATGAGGTAACGAGAGATCGAGC | 1671 |
| Choy_10 | TGTGCCTTGAGACTTCACATGGGCCATGGTGGGAAACGATGAGGTAACGAGAGATCGAGC | 1671 |
| Choy_11_ref | TGTGCCTTGAGACTTCACATGGGCCATGGTGGGAAACGATGAGGTAACGAGAGATCGAGC | 1651 |
| Choy_11 | TGTGCCTTGAGACTTCACATGGGCCATGGTGGGAAACGATGAGGTAACGAGAGATCGAGC | 1651 |
| Choy_12_ref | TGTGCCTTGAGACTTCACATGGGCCATGGTGGGAAACGATGAGGTAACGAGAGATCGAGC | 1612 |
| Choy_12 | TGTGCCTTGAGACTTCACATGGGCCATGGTGGGAAACGATGAGGTAACGAGAGATCGAGC | 1612 |
| Choy_13_ref | TGTGCCTTGAGACTTCACATGGGCCATGGTGGGAAACGATGAGGTAACGAGAGATCGAGC | 1680 |
| Choy_13 | TGTGCCTTGAGACTTCACATGGGCCATGGTGGGAAACGATGAGGTAACGAGAGATCGAGC | 1680 |
| Choy_14_ref | TGTGCCTTGAGACTTCACATGGGCCATGGTGGGAAACGATGAGGTAACGAGAGATCGAGC | 1672 |
| Choy_14 | TGTGCCTTGAGACTTCACATGGGCCATGGTGGGAAACGATGAGGTAACGAGAGATCGAGC | 1672 |
| Choy_5_ref | TGTGCCTTGAGACTTCACATGGGCCATGGTGGGAAACGATGAGGTAACGAGAGATCGAGC | 1680 |
| Choy_5 | TGTGCCTTGAGACTTCACATGGGCCATGGTGGGAAACGATGAGGTAACGAGAGATCGAGC | 1680 |
| Choy_6_ref | TGTGCCTTGAGACTTCACATGGGCCATGGTGGGAAACGATGAGGTAACGAGAGATCGAGC | 1680 |
| Choy_6 | TGTGCCTTGAGACTTCACATGGGCCATGGTGGGAAACGATGAGGTAACGAGAGATCGAGC | 1680 |
| Choy_9_ref | TGTGCCTTGAGACTTCACATGGGCCATGGTGGGAAACGATGAGGTAACGAGAGATCGAGC | 1632 |
| Choy_9 | TGTGCCTTGAGACTTCACATGGGCCATGGTGGGAAACGATGAGGTAACGAGAGATCGAGC | 1632 |
| Pachon_wild_bco2a | | |
| Pachon_1Gross | TGTGCCTTGAGACTTCACATGGGCCATGGTGGGAAACGATGAGGTAACGAGAGATCGAGC | 1482 |
| Pach_11 | TGTGCCTTGAGACTTCACATGGGCCATGGTGGGAAACGATGAGGTAACGAGAGATCGAGC | 1549 |
| Pachon...ss_ref | TGTGCCTTGAGACTTCACATGGGCCATGGTGGGAAACGATGAGGTAACGAGAGATCGAGC | 1482 |
| Pachon2B_ref | TGTGCCTTGAGACTTCACATGGGCCATGGTGGGAAACGATGAGGTAACGAGAGATCGAGC | 1487 |
| Pachon...OG_ref | TGTGCCTTGAGACTTCACATGGGCCATGGTGGGAAACGATGAGGTAACGAGAGATCGAGC | 1481 |
| Pach_12_ref | TGTGCCTTGAGACTTCACATGGGCCATGGTGGGAAACGATGAGGTAACGAGAGATCGAGC | 1472 |
| Pach_12 | TGTGCCTTGAGACTTCACATGGGCCATGGTGGGAAACGATGAGGTAACGAGAGATCGAGC | 1472 |
| Pach_17 | TGTGCCTTGAGACTTCACATGGGCCATGGTGGGAAACGATGAGGTAACGAGAGATCGAGC | 1450 |
| Pachon_F2POG | TGTGCCTTGAGACTTCACATGGGCCATGGTGGGAAACGATGAGGTAACGAGAGATCGAGC | 1485 |
| Pachon_3Gross | TGTGCCTTGAGACTTCACATGGGCCATGGTGGGAAACGATGAGGTAACGAGAGATCGAGC | 1482 |
| Pach_15_ref | TGTGCCTTGAGACTTCACATGGGCCATGGTGGGAAACGATGAGGTAACGAGAGATCGAGC | 1487 |
| Pachon...ro_ref | TGTGCCTTGAGACTTCACATGGGCCATGGTGGGAAACGATGAGGTAACGAGAGATCGAGC | 1489 |
| Pachon...ss_ref | TGTGCCTTGAGACTTCACATGGGCCATGGTGGGAAACGATGAGGTAACGAGAGATCGAGC | 1482 |
| Pachon_2Gross | TGTGCCTTGAGACTTCACATGGGCCATGGTGGGAAACGATGAGGTAACGAGAGATCGAGC | 1533 |
| Pach_17_ref | TGTGCCTTGAGACTTCACATGGGCCATGGTGGGAAACGATGAGGTAACGAGAGATCGAGC | 1450 |
| Pachon...ss_ref | TGTGCCTTGAGACTTCACATGGGCCATGGTGGGAAACGATGAGGTAACGAGAGATCGAGC | 1533 |
| Pachon_ref_ref | TGTGCCTTGAGACTTCACATGGGCCATGGTGGGAAACGATGAGGTAACGAGAGATCGAGC | 1495 |
| Pach_8 | TGTGCCTTGAGACTTCACATGGGCCATGGTGGGAAACGATGAGGTAACGAGAGATCGAGC | 1479 |
| Pach_8_ref | TGTGCCTTGAGACTTCACATGGGCCATGGTGGGAAACGATGAGGTAACGAGAGATCGAGC | 1479 |
| Pachon...OG_ref | TGTGCCTTGAGACTTCACATGGGCCATGGTGGGAAACGATGAGGTAACGAGAGATCGAGC | 1487 |
| Pachon_6Boro | TGTGCCTTGAGACTTCACATGGGCCATGGTGGGAAACGATGAGGTAACGAGAGATCGAGC | 1489 |
| Pachon_ref | TGTGCCTTGAGACTTCACATGGGCCATGGTGGGAAACGATGAGGTAACGAGAGATCGAGC | 1495 |
| Pach_3_ref | TGTGCCTTGAGACTTCACATGGGCCATGGTGGGAAACGATGAGGTAACGAGAGATCGAGC | 1484 |
| Pach_9 | TGTGCCTTGAGACTTCACATGGGCCATGGTGGGAAACGATGAGGTAACGAGAGATCGAGC | 1483 |
| Pachon_G2POG | TGTGCCTTGAGACTTCACATGGGCCATGGTGGGAAACGATGAGGTAACGAGAGATCGAGC | 1481 |
| Pachon...OG_ref | TGTGCCTTGAGACTTCACATGGGCCATGGTGGGAAACGATGAGGTAACGAGAGATCGAGC | 1477 |
| Pach_9_ref | TGTGCCTTGAGACTTCACATGGGCCATGGTGGGAAACGATGAGGTAACGAGAGATCGAGC | 1483 |
| Pachon2B | TGTGCCTTGAGACTTCACATGGGCCATGGTGGGAAACGATGAGGTAACGAGAGATCGAGC | 1487 |
| Pachon...OG_ref | TGTGCCTTGAGACTTCACATGGGCCATGGTGGGAAACGATGAGGTAACGAGAGATCGAGC | 1485 |
| Pachon_E2POG | TGTGCCTTGAGACTTCACATGGGCCATGGTGGGAAACGATGAGGTAACGAGAGATCGAGC | 1487 |
| Pach_14 | TGTGCCTTGAGACTTCACATGGGCCATGGTGGGAAACGATGAGGTAACGAGAGATCGAGC | 1472 |
| Pach_3 | TGTGCCTTGAGACTTCACATGGGCCATGGTGGGAAACGATGAGGTAACGAGAGATCGAGC | 1484 |
| Pachon_H2POG | TGTGCCTTGAGACTTCACATGGGCCATGGTGGGAAACGATGAGGTAACGAGAGATCGAGC | 1477 |
| Pach_7_ref | TGTGCCTTGAGACTTCACATGGGCCATGGTGGGAAACGATGAGGTAACGAGAGATCGAGC | 1419 |
| Pach_15 | TGTGCCTTGAGACTTCACATGGGCCATGGTGGGAAACGATGAGGTAACGAGAGATCGAGC | 1487 |
| Pach_14_ref | TGTGCCTTGAGACTTCACATGGGCCATGGTGGGAAACGATGAGGTAACGAGAGATCGAGC | 1472 |
| Pach_11_ref | TGTGCCTTGAGACTTCACATGGGCCATGGTGGGAAACGATGAGGTAACGAGAGATCGAGC | 1549 |
| Pach_7 | TGTGCCTTGAGACTTCACATGGGCCATGGTGGGAAACGATGAGGTAACGAGAGATCGAGC | 1419 |

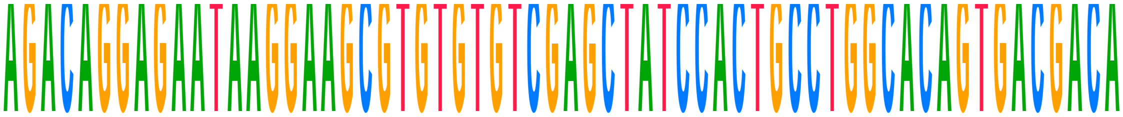

| Name | Sequence | Position |
|---|---|---|
| **Consensus** | **AGACAGGAGAATAAGGAAGCGTGTGTGTCGAGCTATCCACTGCCTGGCACAGTGACGACA** | |
| Choy_1_ref | AGACAGGAGAATAAGGAAGCGTGTGTGTCGAGCTATCCACTGCCTGGCACAGTGACGACA | 1666 |
| Choy_1 | AGACAGGAGAATAAGGAAGCGTGTGTGTCGAGCTATCCACTGCCTGGCACAGTGACGACA | 1666 |
| Choy_10_ref | AGACAGGAGAATAAGGAAGCGTGTGTGTCGAGCTATCCACTGCCTGGCACAGTGACGACA | 1731 |
| Choy_10 | AGACAGGAGAATAAGGAAGCGTGTGTGTCGAGCTATCCACTGCCTGGCACAGTGACGACA | 1731 |
| Choy_11_ref | AGACAGGAGAATAAGGAAGCGTGTGTGTCGAGCTATCCACTGCCTGGCACAGTGACGACA | 1711 |
| Choy_11 | AGACAGGAGAATAAGGAAGCGTGTGTGTCGAGCTATCCACTGCCTGGCACAGTGACGACA | 1711 |
| Choy_12_ref | AGACAGGAGAATAAGGAAGCGTGTGTGTCGAGCTATCCACTGCCTGGCACAGTGACGACA | 1672 |
| Choy_12 | AGACAGGAGAATAAGGAAGCGTGTGTGTCGAGCTATCCACTGCCTGGCACAGTGACGACA | 1672 |
| Choy_13_ref | AGACAGGAGAATAAGGAAGCGTGTGTGTCGAGCTATCCACTGCCTGGCACAGTGACGACA | 1740 |
| Choy_13 | AGACAGGAGAATAAGGAAGCGTGTGTGTCGAGCTATCCACTGCCTGGCACAGTGACGACA | 1740 |
| Choy_14_ref | AGACAGGAGAATAAGGAAGCGTGTGTGTCGAGCTATCCACTGCCTGGCACAGTGACGACA | 1732 |
| Choy_14 | AGACAGGAGAATAAGGAAGCGTGTGTGTCGAGCTATCCACTGCCTGGCACAGTGACGACA | 1732 |
| Choy_5_ref | AGACAGGAGAATAAGGAAGCGTGTGTGT----CTATCCACTGCCTGGCACAGTGACGACA | 1736 |
| Choy_5 | AGACAGGAGAATAAGGAAGCGTGTGTGT----CTATCCACTGCCTGGCACAGTGACGACA | 1736 |
| Choy_6_ref | AGACAGGAGAATAAGGAAGCGTGTGTGTC---CTATCCACTGCCTGGCACAGTGACGACA | 1737 |
| Choy_6 | AGACAGGAGAATAAGGAAGCGTGTGTGTC---CTATCCACTGCCTGGCACAGTGACGACA | 1737 |
| Choy_9_ref | AGACAGGAGAATAAGGAAGCGTGTGTGTCGAGCTATCCACTGCCTGGCACAGTGACGACA | 1692 |
| Choy_9 | AGACAGGAGAATAAGGAAGCGTGTGTGTCGAGCTATCCACTGCCTGGCACAGTGACGACA | 1692 |
| Pachon_wild_bco2a | | |
| Pachon_1Gross | AGACAGGAGAATAAGGAAGCGTGTGTGTCGAGCTATCCACTGCCTGGCACAGTGACGACA | 1542 |
| Pach_11 | AGACAGGAGAATAAGGAAGCGTGTGTGTCGAGCTATCCACTGCCTGGCACAGTGACGACA | 1609 |
| Pachon...ss_ref | AGACAGGAGAATAAGGAAGCGTGTGTGTCGAGCTATCCACTGCCTGGCACAGTGACGACA | 1542 |
| Pachon2B_ref | AGACAGGAGAATAAGGAAGCGTGTGTGTCGAGCTATCCACTGCCTGGCACAGTGACGACA | 1547 |
| Pachon...OG_ref | AGACAGGAGAATAAGGAAGCGTGTGTGTCGAGCTATCCACTGCCTGGCACAGTGACGACA | 1541 |
| Pach_12_ref | AGACAGGAGAATAAGGAAGCGTGTGTGTCGAGCTATCCACTGCCTGGCACAGTGACGACA | 1532 |
| Pach_12 | AGACAGGAGAATAAGGAAGCGTGTGTGTCGAGCTATCCACTGCCTGGCACAGTGACGACA | 1532 |
| Pach_17 | AGACAGGAGAATAAGGAAGCGTGTGTGTCGAGCTATCCACTGCCT------GTGACGACA | 1504 |
| Pachon_F2POG | AGACAGGAGAATAAGGAAGCGTGTGTGTCGAGCTATCCACTGCCTGGCACAGTGACGACA | 1545 |
| Pachon_3Gross | AGACAGGAGAATAAGGAAGCGTGTGTGTCGAGCTATCCACTGCCTGGCACAGTGACGACA | 1542 |
| Pach_15_ref | AGACAGGAGAATAAGGAAGCGTGTGTGTCGAGCTATCCACTGCCTGGCACAGTGACGACA | 1547 |
| Pachon...ro_ref | AGACAGGAGAATAAGGAAGCGTGTGTGTCGAGCTATCCACTGCCTGGCACAGTGACGACA | 1549 |
| Pachon...ss_ref | AGACAGGAGAATAAGGAAGCGTGTGTGTCGAGCTATCCACTGCCTGGCACAGTGACGACA | 1542 |
| Pachon_2Gross | AGACAGGAGAATAAGGAAGCGTGTGTGTCGAGCTATCCACTGCCTGGCACAGTGACGACA | 1593 |
| Pach_17_ref | AGACAGGAGAATAAGGAAGCGTGTGTGTCGAGCTATCCACTGCCT------GTGACGACA | 1504 |
| Pachon...ss_ref | AGACAGGAGAATAAGGAAGCGTGTGTGTCGAGCTATCCACTGCCTGGCACAGTGACGACA | 1593 |
| Pachon_ref_ref | AGACAGGAGAATAAGGAAGCGTGTGTGTCGAGCTATCCACTGCCTGGCACAGTGACGACA | 1555 |
| Pach_8 | AGACAGGAGAATAAGGAAGCGTGTGTGTCGAGCTATCCACTGCCTGGCACAGTGACGACA | 1539 |
| Pach_8_ref | AGACAGGAGAATAAGGAAGCGTGTGTGTCGAGCTATCCACTGCCTGGCACAGTGACGACA | 1539 |
| Pachon...OG_ref | AGACAGGAGAATAAGGAAGCGTGTGTGTCGAGCTATCCACTGCCTGGCACAGTGACGACA | 1547 |
| Pachon_6Boro | AGACAGGAGAATAAGGAAGCGTGTGTGTCGAGCTATCCACTGCCTGGCACAGTGACGACA | 1549 |
| Pachon_ref | AGACAGGAGAATAAGGAAGCGTGTGTGTCGAGCTATCCACTGCCTGGCACAGTGACGACA | 1555 |
| Pach_3_ref | AGACAGGAGAATAAGGAAGCGTGTGTGTCGAGCTATCCACTGCCTGGCACAGTGACGACA | 1544 |
| Pach_9 | AGACAGGAGAATAAGGAAGCGTGTGTGTCGAGCTATCCACTGCCTGGCACAGTGACGACA | 1543 |
| Pachon_G2POG | AGACAGGAGAATAAGGAAGCGTGTGTGTCGAGCTATCCACTGCCTGGCACAGTGACGACA | 1541 |
| Pachon...OG_ref | AGACAGGAGAATAAGGAAGCGTGTGTGTCGAGCTATCCACTGCCTGGCACAGTGACGACA | 1537 |
| Pach_9_ref | AGACAGGAGAATAAGGAAGCGTGTGTGTCGAGCTATCCACTGCCTGGCACAGTGACGACA | 1543 |
| Pachon2B | AGACAGGAGAATAAGGAAGCGTGTGTGTCGAGCTATCCACTGCCTGGCACAGTGACGACA | 1547 |
| Pachon...OG_ref | AGACAGGAGAATAAGGAAGCGTGTGTGTCGAGCTATCCACTGCCTGGCACAGTGACGACA | 1545 |
| Pachon_E2POG | AGACAGGAGAATAAGGAAGCGTGTGTGTCGAGCTATCCACTGCCTGGCACAGTGACGACA | 1547 |
| Pach_14 | AGACAGGAGA----------------------CTATCCACTGCCTGGCACAGTGACGACA | 1511 |
| Pach_3 | AGACAGGAGAATAAGGAAGCGTGTGTGTCGAGCTATCCACTGCCTGGCACAGTGACGACA | 1544 |
| Pachon_H2POG | AGACAGGAGAATAAGGAAGCGTGTGTGTCGAGCTATCCACTGCCTGGCACAGTGACGACA | 1537 |
| Pach_7_ref | AGACAGGAGAATAAGGAAGCGTGTGTGTCGAGCTATCCACTGCCTGGCACAGTGACGACA | 1479 |
| Pach_15 | AGACAGGAGAATAAGGAAGCGTGTGTGTCGAGCTATCCACTGCCTGGCACAGTGACGACA | 1547 |
| Pach_14_ref | AGACAGGAGA----------------------CTATCCACTGCCTGGCACAGTGACGACA | 1511 |
| Pach_11_ref | AGACAGGAGAATAAGGAAGCGTGTGTGTCGAGCTATCCACTGCCTGGCACAGTGACGACA | 1609 |
| Pach_7 | AGACAGGAGAATAAGGAAGCGTGTGTGTCGAGCTATCCACTGCCTGGCACAGTGACGACA | 1479 |

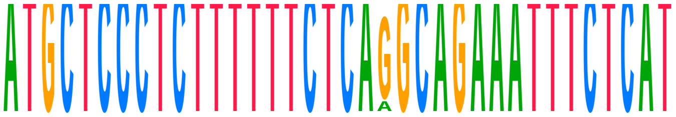


| Name | Sequence | Position |
|---|---|---|
| **Consensus** | **ATGCTCCCTCTTTTTTCTCAGGCAGAAATTTCTCAT** | |
| Choy_1_ref | ATGCTCCCTCTTTTTTCTCAGGCAGAAATTTCTCAT | 1702 |
| Choy_1 | ATGCTCCCTCTTTTTTCTCAGGCAGAAATTTCTCAT | 1702 |
| Choy_10_ref | ATGCTCCCTCTTTTTTCTCAGGCAGAAATTTCTCAT | 1767 |
| Choy_10 | ATGCTCCCTCTTTTTTCTCAAGCAGAAATTTCTCAT | 1767 |
| Choy_11_ref | ATGCTCCCTCTTTTTTCTCAGGCAGAAATTTCTCAT | 1747 |
| Choy_11 | ATGCTCCCTCTTTTTTCTCAAGCAGAAATTTCTCAT | 1747 |
| Choy_12_ref | ATGCTCCCTCTTTTTTCTCAGGCAGAAATTTCTCAT | 1708 |
| Choy_12 | ATGCTCCCTCTTTTTTCTCAAGCAGAAATTTCTCAT | 1708 |
| Choy_13_ref | ATGCTCCCTCTTTTTTCTCAGGCAGAAATTTCTCAT | 1776 |
| Choy_13 | ATGCTCCCTCTTTTTTCTCAGGCAGAAATTTCTCAT | 1776 |
| Choy_14_ref | ATGCTCCCTCTTTTTTCTCAGGCAGAAATTTCTCAT | 1768 |
| Choy_14 | ATGCTCCCTCTTTTTTCTCAAGCAGAAATTTCTCAT | 1768 |
| Choy_5_ref | ATGCTCCCTCTTTTTTCTCAGGCAGAAATTTCTCAT | 1772 |
| Choy_5 | ATGCTCCCTCTTTTTTCTCAAGCAGAAATTTCTCAT | 1772 |
| Choy_6_ref | ATGCTCCCTCTTTTTTCTCAGGCAGAAATTTCTCAT | 1773 |
| Choy_6 | ATGCTCCCTCTTTTTTCTCAAGCAGAAATTTCTCAT | 1773 |
| Choy_9_ref | ATGCTCCCTCTTTTTTCTCAGGCAGAAATTTCTCAT | 1728 |
| Choy_9 | ATGCTCCCTCTTTTTTCTCAGGCAGAAATTTCTCAT | 1728 |
| Pachon_wild_bco2a | | |
| Pachon_1Gross | ATGCTCCCTCTTTTTTCTCAGGCAGAAATTTCTCAT | 1578 |
| Pach_11 | ATGCTCCCTCTTTTTTCTCAGGCAGAAATTTCTCAT | 1645 |
| Pachon…ss_ref | ATGCTCCCTCTTTTTTCTCAGGCAGAAATTTCTCAT | 1578 |
| Pachon2B_ref | ATGCTCCCTCTTTTTTCTCAGGCAGAAATTTCTCAT | 1583 |
| Pachon…OG_ref | ATGCTCCCTCTTTTTTCTCAGGCAGAAATTTCTCAT | 1577 |
| Pach_12_ref | ATGCTCCCTCTTTTTTCTCAGGCAGAAATTTCTCAT | 1568 |
| Pach_12 | ATGCTCCCTCTTTTTTCTCAGGCAGAAATTTCTCAT | 1568 |
| Pach_17 | ATGCTCCCTCTTTTTTCTCAGGCAGAAATTTCTCAT | 1540 |
| Pachon_F2POG | ATGCTCCCTCTTTTTTCTCAGGCAGAAATTTCTCAT | 1581 |
| Pachon_3Gross | ATGCTCCCTCTTTTTTCTCAGGCAGAAATTTCTCAT | 1578 |
| Pach_15_ref | ATGCTCCCTCTTTTTTCTCAGGCAGAAATTTCTCAT | 1583 |
| Pachon…ro_ref | ATGCTCCCTCTTTTTTCTCAGGCAGAAATTTCTCAT | 1585 |
| Pachon…ss_ref | ATGCTCCCTCTTTTTTCTCAGGCAGAAATTTCTCAT | 1578 |
| Pachon_2Gross | ATGCTCCCTCTTTTTTCTCAGGCAGAAATTTCTCAT | 1629 |
| Pach_17_ref | ATGCTCCCTCTTTTTTCTCAGGCAGAAATTTCTCAT | 1540 |
| Pachon…ss_ref | ATGCTCCCTCTTTTTTCTCAGGCAGAAATTTCTCAT | 1629 |
| Pachon_ref_ref | ATGCTCCCTCTTTTTTCTCAGGCAGAAATTTCTCAT | 1591 |
| Pach_8 | ATGCTCCCTCTTTTTTCTCAGGCAGAAATTTCTCAT | 1575 |
| Pach_8_ref | ATGCTCCCTCTTTTTTCTCAGGCAGAAATTTCTCAT | 1575 |
| Pachon…OG_ref | ATGCTCCCTCTTTTTTCTCAGGCAGAAATTTCTCAT | 1583 |
| Pachon_6Boro | ATGCTCCCTCTTTTTTCTCAGGCAGAAATTTCTCAT | 1585 |
| Pachon_ref | ATGCTCCCTCTTTTTTCTCAGGCAGAAATTTCTCAT | 1591 |
| Pach_3_ref | ATGCTCCCTCTTTTTTCTCAGGCAGAAATTTCTCAT | 1580 |
| Pach_9 | ATGCTCCCTCTTTTTTCTCAGGCAGAAATTTCTCAT | 1579 |
| Pachon_G2POG | ATGCTCCCTCTTTTTTCTCAGGCAGAAATTTCTCAT | 1577 |
| Pachon…OG_ref | ATGCTCCCTCTTTTTTCTCAGGCAGAAATTTCTCAT | 1573 |
| Pach_9_ref | ATGCTCCCTCTTTTTTCTCAGGCAGAAATTTCTCAT | 1579 |
| Pachon2B | ATGCTCCCTCTTTTTTCTCAGGCAGAAATTTCTCAT | 1583 |
| Pachon…OG_ref | ATGCTCCCTCTTTTTTCTCAGGCAGAAATTTCTCAT | 1581 |
| Pachon_E2POG | ATGCTCCCTCTTTTTTCTCAGGCAGAAATTTCTCAT | 1583 |
| Pach_14 | ATGCTCCCTCTTTTTTCTCAGGCAGAAATTTCT--- | 1544 |
| Pach_3 | ATGCTCCCTCTTTTTTCTCAGGCAGAAATTTCTCAT | 1580 |
| Pachon_H2POG | ATGCTCCCTCTTTTTTCTCAGGCAGAAATTTCTCAT | 1573 |
| Pach_7_ref | ATGCTCCCTCTTTTTTCTCAGGCAGAAATTTCTCAT | 1515 |
| Pach_15 | ATGCTCCCTCTTTTTTCTCAGGCAGAAATTTCTCAT | 1583 |
| Pach_14_ref | ATGCTCCCTCTTTTTTCTCAGGCAGAAATTTCT--- | 1544 |
| Pach_11_ref | ATGCTCCCTCTTTTTTCTCAGGCAGAAATTTCTCAT | 1645 |
| Pach_7 | ATGCTCCCTCTTTTTTCTCAGGCAGAAATTTCTCAT | 1515 |

**Sequence Logo:** 50% GC base composition

**Consensus Threshold:** >50%

**Compare to:** the consensus

Bases that don't match the reference are marked with yellow highlighting.

**Created:** May 9, 2024

**Last Modified:** May 9, 2024

Aligned using Global alignment (Needleman-Wunsch)

### Supplemental Figure 4

**Pachón cavefish consensus**
**Río Choy Surface consensus**

```
1   MRNFCLRKKREHCRHCARQWIARHTRFLILLSARSLVTSSFPTMAHVKSQ  50
1   MRNFCLRKKREHCRHCARQWIARHTRFLILLSARSLVTSSFPTMAHVKSQ  50

                                                W96R in L1 Cave
51  GTVANNKQNNGFKDQHGLQCIAPLVRTAEETPEPIPTTVKGTIPSWIRGS  100
51  GTVANNKQNNGFKDQHGLQCIAPLVRTAEETPEPIPTTVKGTIPSWIRGS  100

101 LLRNGPGKFEFGNQHFNHWFDGMALMHRFEIMDGQVTYRSRFLRSDSYNL  150
101 LLRNGPGKFEFGNQHFNHWFDGMALMHRFEIMDGQVTYRSRFLRSDSYNL  150

              Pachón short end |              YLK
151 NSEKNRIMVSEFGTMSFPDPCKN-----------YXPTDNASVNFVKYKG  189
151 NSEKNRIMVSEFGTMSFPDPCKNLFQRFLSRFEMMKPTDNASVNFVKYKG  200

                Pachón long end |
190 DYYVSTETNFMHKVDPDNLETKEKV-------------------------  214
201 DYYVSTETNFMHKVDPDNLETKEKVDWSKFIAVNGATAHPHFDPDGTAYN  250

214 --------------------------------------------------  214
251 MGNSYGRKGALYNIISVPPEKDGNTDTLQGAKVLCSVIPTDKARPSYYHS  300

                                   V332L in L1 Cave
214 --------------------------------------------------  214
301 FAMSENYVVIIEQSVKMDLFKIVTARLRNKNVSNGVYWDPKQETVFHLVN  350

215 RHTXVL--------------------------------------------  220
351 KHTGEVSPVKYHTTPLSTFHQINAFEEDGFLMMDLCGSDDGKAINTYLIQ  400

220 --------------------------------------------------  220
401 NLRKTGEALDEVYNSMGRAFPRRYVLPLNITNETPLQQNLNTRPNSSATA  450

220 -------------------------------------------------R  221
451 VCHSSEKVFCTSEDLHDEDLLEYGGLEFPQIYYSKYNTKPYRYFYGCGFR  500

222 H-------------------------------------------------  222
501 HLVGDSLIKMDLNNKKLKAWHQPGFYPSEPVFVPSPEAEEEDDGVILSVV  550
```

222 -------------------------------PY---------* 225
551 LTPTEDNSTFLLVLDAKTFEEMGRAEVPVNIPYGFHGVFNAA 592

Supplemental Figure 5

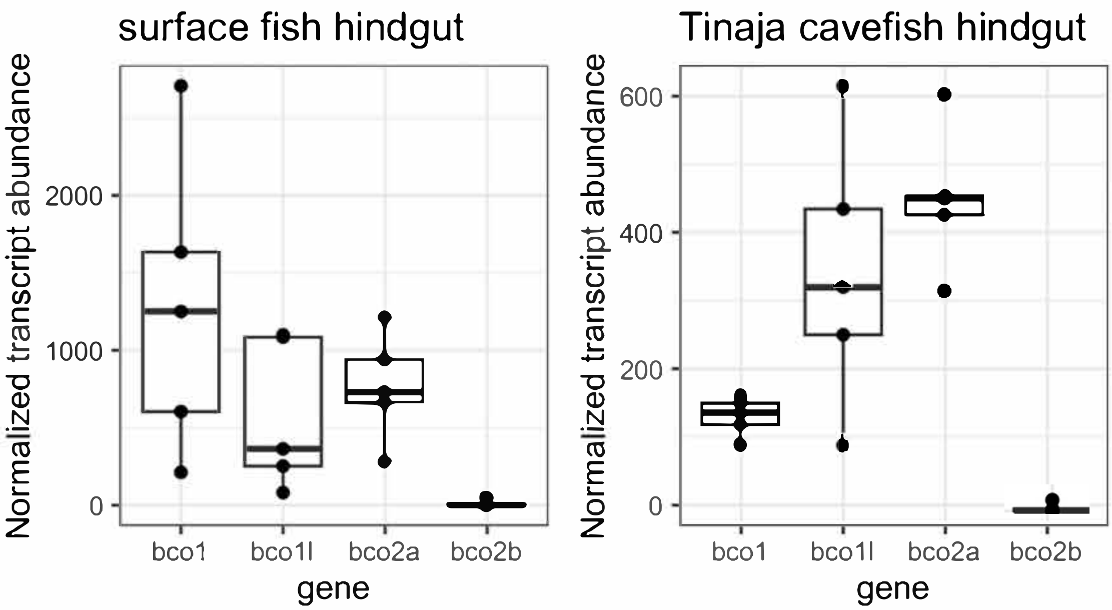
